# An updated compendium and reevaluation of the evidence for nuclear transcription factor occupancy over the mitochondrial genome

**DOI:** 10.1101/2024.06.04.597442

**Authors:** Georgi K. Marinov, Vivekanandan Ramalingam, William J. Greenleaf, Anshul Kundaje

**Affiliations:** Department of Genetics, Stanford University, Stanford, CA 94305, USA; Department of Computer Science, Stanford University, Stanford, CA 94305, USA; Center for Personal Dynamic Regulomes, Stanford University, Stanford, California 94305, USA; Department of Applied Physics, Stanford University, Stanford, California 94305, USA; Chan Zuckerberg Biohub, San Francisco, California, USA

## Abstract

In most eukaryotes, mitochondrial organelles contain their own genome, usually circular, which is the remnant of the genome of the ancestral bacterial endosymbiont that gave rise to modern mitochondria. Mitochondrial genomes are dramatically reduced in their gene content due to the process of endosymbiotic gene transfer to the nucleus; as a result most mitochondrial proteins are encoded in the nucleus and imported into mitochondria. This includes the components of the dedicated mitochondrial transcription and replication systems and regulatory factors, which are entirely distinct from the information processing systems in the nucleus. However, since the 1990s several nuclear transcription factors have been reported to act in mitochondria, and previously we identified 8 human and 3 mouse transcription factors (TFs) with strong localized enrichment over the mitochondrial genome using ChIP-seq (Chromatin Immunoprecipitation) datasets from the second phase of the ENCODE (Encyclopedia of DNA Elements) Project Consortium. Here, we analyze the greatly expanded in the intervening decade ENCODE compendium of TF ChIP-seq datasets (a total of 6,153 ChIP experiments for 942 proteins, of which 763 are sequence-specific TFs) combined with interpretative deep learning models of TF occupancy to create a comprehensive compendium of nuclear TFs that show evidence of association with the mitochondrial genome. We find some evidence for chrM occupancy for 50 nuclear TFs and two other proteins, with bZIP TFs emerging as most likely to be playing a role in mitochondria. However, we also observe that in cases where the same TF has been assayed with multiple antibodies and ChIP protocols, evidence for its chrM occupancy is not always reproducible. In the light of these findings, we discuss the evidential criteria for establishing chrM occupancy and reevaluate the overall compendium of putative mitochondrial-acting nuclear TFs.

## Introduction

Mitochondria contain their own genome ^2^ (mtDNA/chrM), usually circular in topology and highly compact, especially in metazoans. The mammalian chrM is around 16-17 kbp in size and encodes for 13 proteins (all of them components of electron transport chains), 22 tRNAs and two rRNAs ^3,4^. It has a peculiar compared to the nuclear genome organization and is replicated, transcribed and regulated by its own dedicated set of information processing factors. The 13 genes and two rRNAs are densely packed, with only one notable non-coding region (NCR) – the so called displacement or D-loop ^5^. The D-loop is major the site of replication and transcription initiation, the latter producing long polycistronic transcripts from both strands (referred to as the H- for “heavy” and L- for “light”), and contains three promoters – a light strand (LSP) and two heavy strand (HSP1 and HSP2) promoters ^6,7^. Transcription is carried out by the mitochondrial-specific nuclear-encoded POLRMT RNA polymerase ^8^, with several additional nuclear genome-encoded factors – TFAM ^9–11^, TFB1M, and TFB2M ^12,13^ – also involved in the process of initiation ^14^.

The modern organization of the mitochondrial genome is the result of a combination of extreme reduction and massive endosymbiotic gene transfer (EGT). Mitochondria originated very early in eukaryote evolution as a result of the establishment of endosymbiosis with their prokaryotic ancestor ^15–20^, which was most likely a member of the *α*-proteobacteria clade ^20–23^. Subsequently, the endosymbiont lost the vast majority of its genes, either outright, or through transfer to the nuclear genome, form which then their products were imported back into the mitochondrion.

In addition to the well-characterized bona fide mitochondrial transcription and replication factors, since the 1990s there have been reports suggesting that nuclear transcription factors may also moonlight as direct regulators of events in mitochondria ^24–27^. These include the glucocorticoid receptor GR ^28–31^, a 43kDa isoform of the thyroid hormone T_3_ receptor T_3_R*α*1 called p43 ^32–35^, the CREB TF ^36–39^, the tumor suppressor transcription factor p53 ^40–43^, the estrogen receptor ER ^44,45^, STAT3 ^46^, AP-1 and PPAR*γ*2 ^47–49^, as well as MEF2D in mouse ^50^.

However, direct *in vivo* evidence for occupancy of mtDNA by nuclear factors was provided only for a handful of them by these original studies (e.g. CREB ^37^ and p53 ^43^, and it was limited to only the D-loop region. The advent of high resolution techniques for genome-wide profiling of DNA-protein interactions such as ChIP-seq (Chromatin Immunoprecipitation coupled with deep sequencing ^51–54^) eventually enabled the direct examination of evidence for mtDNA occupancy by a large number of nuclear TFs.

At the end of the second phase of the ENCODE Project, we and others ^26,27^ carried out a comprehensive survey of the existing at the time ChIP-seq datasets generated by the ENCODE, mouseENCODE and modENCODE efforts ^55–61^ in human, mouse, the worm *C. elegans* and the fly *D. melanogaster*. Eight human TFs were identified as showing strong evidence for mtDNA occupancy (JUN, JUND, CEBPB, MAX, MafF, MafK, NFE2 and RFX5), three mouse TFs (MafK, MafF and Usf2), and no fly or worm ones. Furthermore, Blumberg et al. ^27^ demonstrated directly the localization to mitochondria of JUN and JUND in HepG2 cells using immuno-gold labeling and electron microscopy while Marinov et al. ^26^ showed MAFK localizing to mitochondria using immunocytochemical staining. Examination of available ChIP-seq data for TFs previously proposed to act in mitochondria (GR, ER*α*, CREB, STAT3, p53) found no putative occupancy sites.

However, these studies did not reveal any obvious mechanisms through which these nuclear TFs might act to regulate mitochondrial transcription, as all the identified ChIP-seq peaks are away from the D-loop in the middle of the transcriptional units. The D-loop itself shows up as “enriched” in practically all ChIP-seq datasets, but this is almost certainly an experimental artifact as the ChIP signal there does not show the characteristics of proper occupancy sites. Thus the question about the potential role of nuclear TFs in mitochondria remains open and unresolved ^24,25^.

With the completion of the third ^62^ and fourth phases of the ENCODE project, a vastly expanded collection of ChIP-seq datasets has now become available, encompassing an order-of-magnitude larger sampling of the human TF repertoire. Furthermore, many TFs have now been assayed using multiple different reagents or using endogenous tagging, thus potentially providing distinct lines of evidence for mtDNA occupancy, and powerful deep learning-based tools for analyzing the sequence patterns driving TF occupancy and predicting it from sequence have been developed ^63^. In this study, we take advantage of these resources, survey the expanded ENCODE collection, and identify 50 nuclear TFs plus two other chromatin proteins exhibiting more or less robustly supported peaks over chrM. On the other hand, the picture revealed by the expanded collection is more complicated than previously perceived as in many cases occupancy profiles are not replicated in all cases where multiple immune reagents have been used to assay the same TF. We discuss the currently most reliable set of mtDNA-associated nuclear TFs, as well as the evidential criteria for establishing chrM occupancy using ChIP and other experimental methods.

## Results

### Evaluating the evidence for mitochondrial occupancy of nuclear TFs in the expanded ENCODE ChIP-seq collection

According to the latest census of human transcription factors ^1^, the human genome encodes 1,742 sequence-specific TFs, belonging to *∼*60 different families defined by their DNA binding domains (DBDs).

Our previous analysis of the dataset collection generated as part of the second phase of the ENCODE Project encompassed a total of 151 transcription factors, which represents less than 10% of the total. After the fourth phase of ENCODE, the number of available datasets is now greatly expanded and covers *∼*44% of the known TFs.

In order to evaluate putative physical associations of nuclear TFs with the mitochondrial genome (and also reevaluate previous observations), we examined 6,153 ChIP-seq datasets for 942 targets, of which 763 are sequence-specific TFs ^1^ (Figure 1A). The additional 179 are non-sequence specific chromatin proteins, such as histone modifying enzymes and chromatin remodellers.

**Figure 1:**
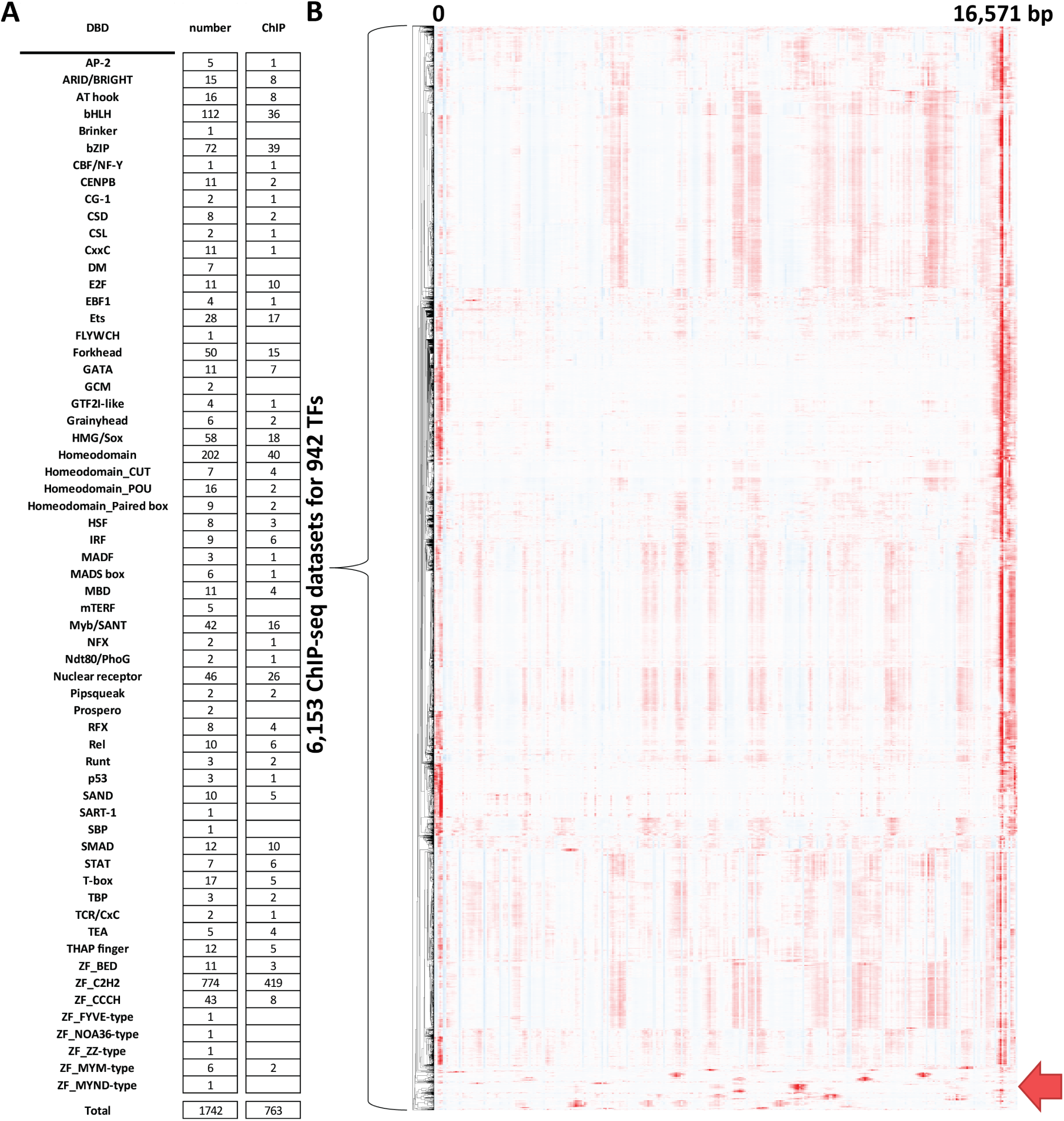
Global assessment of the evidence for association of human nuclear TFs with the mitochondrial genome.. (A) Summary of the known human TFs and available ENCODE TF ChIP datasets. The TF classification of human TFs of Lambert et al. 2018 ^1^ was followed. (B) Hierarchical clustering of ChIP-seq profiles over the mitochondrial genome for 6,513 TF ChIP-seq datasets. Datasets that show evidence for non-artefactual association with mitochondrial DNA are highlighted at the bottom.

The most recent ENCODE collection is also not merely a quantitative expansion. Multiple TFs have been assayed with different antibodies, which is extremely valuable because, especially when the antibodies are polyclonal, the possibility that mtDNA ChIP-seq peaks are the result of non-specific pull down can never be excluded (although that would still mean that some previously not known to do so protein is specifically occupying those regions of the mitochondrial genome). Many of the newly available TFs, but also some that have been mapped previously, have now been assayed using endogenous tagging with FLAG, GFP or HA tags, most commonly using CRISPR epitope tagging (CETCH-seq ^64^) and site-specific recombination ^65^. We note that in CETCH-eq tagging is carried out at the stop codon of the gene, i.e. at its C-terminus, thus it is unlikely that it would affect import into mitochondria, which is classically mediated by N-terminal targeting sequences ^66,67^; however, that can nevertheless not be completely excluded as a possibility.

We updated our pipeline for analyzing mitochondrial occupancy in several ways in order to allow for more comprehensive characterization of the current TF compendium.

First, we previously mapped reads against a joint nuclear and mitochondrial genomic index, and excluded all reads mapping to multiple locations within that combined genomic space. A substantial portion of the human chrM in the context of the hg38 assembly is affected by such mappability limitations (Supplementary Figure 1). However, because of the very high copy number of mitochondrial genomes in a given cell relative to the diploid nuclear genome ^68^, peaks observed over chrM are nearly always much stronger than even the very top nuclear ChIP-seq peaks, as previously demonstrated ^26^. Furthermore, other hallmarks of TF occupancy, such as chromatin accessibility peaks and high levels of histone marks associated with active regulatory elements, which would be expected if chrM peaks arose from mitochondrial sequences that have been inserted in the nuclear genome (so called NUMTs ^69^), are not seen over the peaks observed over the mitochondrial genome. This makes it highly unlikely that they arise from nuclear TF occupancy over NUMTs. For these reasons, we now evaluate putative mitochondrial occupancy based on read alignments generated entirely in mitochondrial space.

Second, in our past work we sought corroborating evidence for the observed ChIP-seq profiles in the presence or absence of the cognate sequence recognition motifs for each TF. However, these motifs are often very short and degenerate, and thus only a small fraction of them is actually occupied in cells. In this work, we leverage the power of interpretative deep learning models to generate more reliable and specific predictions of TF occupancy profiles over chrM, which we then compare to experimental measurements. We use the state-of-the-art BPNet ^70^ profile models, which take as input genomic sequence and the forward- and reverse-strand ChIP-seq profiles, and then learn to predict these profiles as a function of genomic sequence. As part of the overall ENCODE effort, we have trained such models over the nuclear genome for all TFs for which data is available, and we used these models to predict chrM ChIP-seq profiles (see Methods for details).

Figure 1B shows the observed chrM profiles for all 6,153 datasets. As discussed in our previous work ^26^, the D-loop region appears as strongly “enriched” in nearly all ChIP datasets; this is certainly an artifact in almost all cases because the observed forward- and reverse-strand profiles do not exhibit the expected from true occupancy asymmetry around a punctate binding site ^71–74^. It is most likely that the unique triple-stranded structure of the D-loop results in preferential enrichment in sequencing libraries. We also observe a few regions of weakly elevated signal in the middle of chrM, which are also present in the majority of datasets, and are also unlikely to represent true occupancy events.

Disregarding these signals, we find some evidence for chrM occupancy for 50 sequence-specific TFs, which we discuss in detail below. In addition, two of the 179 non-sequence specific chromatin proteins also showed evidence for putative association with mtDNA.

### bZIP TFs

The TF family with the largest and most notable set of members with strong chrM peaks is the bZIP (Basic Leucine Zipper) domain-containing proteins. In humans, 72 such TFs are annotated in the genome, and for 39 of them there is ENCODE ChIP-seq data.

Remarkably, nearly half of them – 19/39 – exhibit evidence fo mtDNA occupancy (Figures 2–6A-C and Supplementary Figures 2–22). These TFs are ATF2, ATF3, ATF4, ATF7, CREB1, FOS, FOSL1, FOSL2, CEBPB, CEBPG, JUN, JUND, MAFF, MAFG, MAFK, NFE2, NFE2L1, NFE2L2, and NRL.

**Figure 2:**
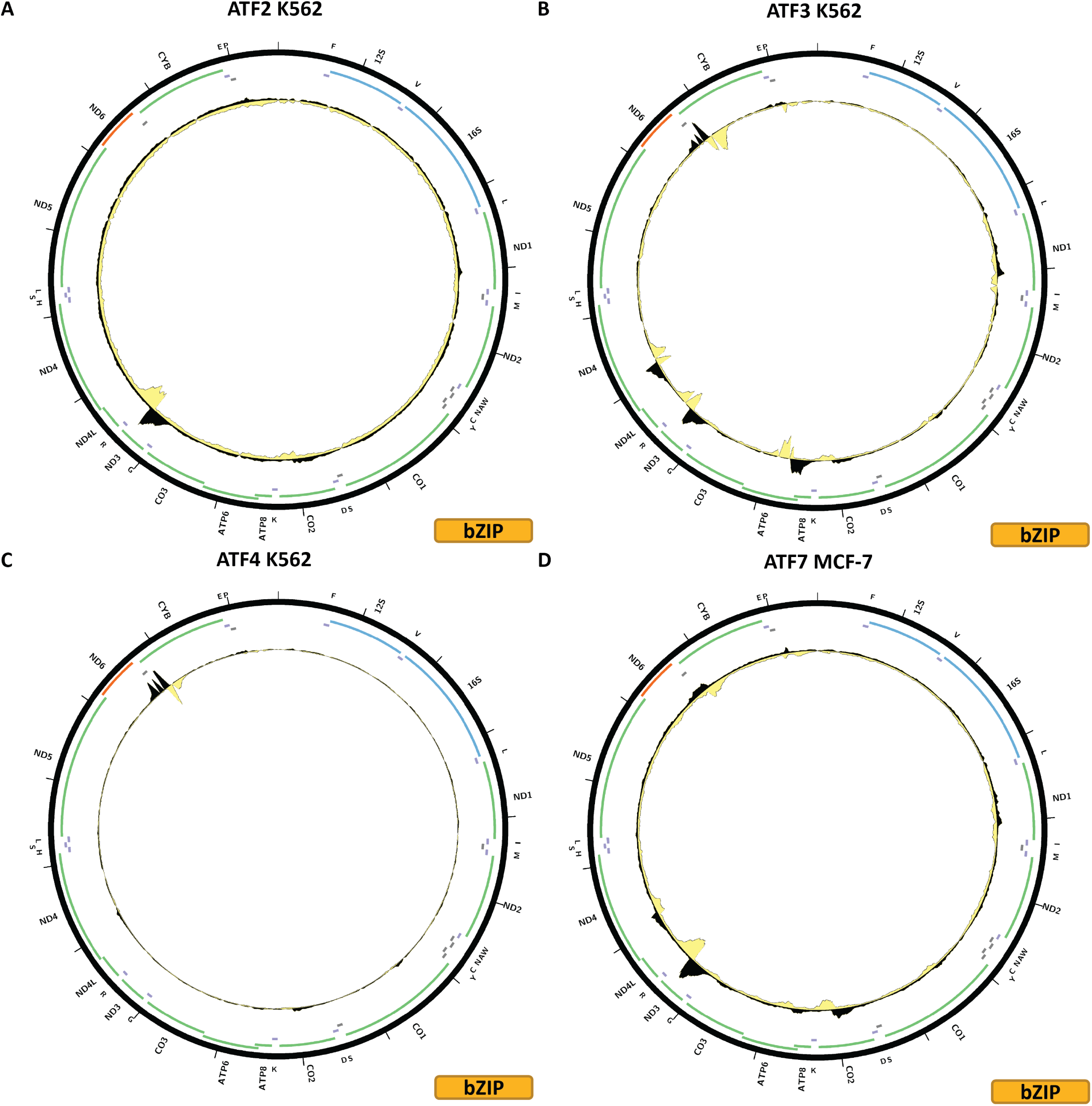
Evidence for mitochondrial genome occupancy by nuclear transcription factors. Black and yellow tracks show the forward- and reverse-strand ChIP-seq coverage over chrM. (A) ATF2 (bZIP); (B) ATF3 (bZIP); (C) ATF4 (bZIP); (D) ATF7 (bZIP).

Figure 2A shows the chrM ChIP-seq profile for the ATF2 TF in the K562 cell line, revealing a strong peak, with the classic strand asymmetry features of true sequence-specific TF occupancy, in the region around the *MT-ND3* and *MT-ND4L* genes and the *MT-TR* tRNA gene. The BPNet-predicted profile corroborates the existence of such a peak, although BPNet also predicts high ChIP-seq signal in several other locations, which is not observed in the actual data (Supplementary Figure 2B). The same peak is observed in HepG2 cells using the same antibody (Supplementary Figure 2E), and in site-recombination-tagged experiment in the HEK293 cell line (Supplementary Figure 2D). It is not observed in H1-hESC using a different antibody (Supplementary Figure 2C), and also in a CRISPR-mediated endogenous tagging experiment in HepG2 cells (Supplementary Figure 2F). Thus for ATF2 we see evidence for occupancy using two different reagents (an *α*-ATF2 antibody and GFP-tagging), but not with another *α*-ATF2 antibody and FLAG-tagging.

Figure 2B shows the chrM ChIP-seq profile for the ATF3 TF in the K562 cell line. In this case, at least four strong peaks are observed – the same one around *MT-ND3* /*MT-ND4L* seen for ATF2, but also one over *MT-ATP6*, one over *MT-ND4*, and another over *MT-CYB*. All these peaks are corroborated by BPNet predictions (Supplementary Figure 3B), although here too BPNet predicts additional occupancy peaks. However, none of the experiments in other cell lines – H1-hESC, HCT116, HepG2, A549, GM12878, liver, and K562 again – carried out with a different antibody exhibit these peaks (Supplementary Figure 3C–I), and neither does CRISPR-mediated FLAG-tagging (Supplementary Figure 4J).

Figure 2C shows the chrM ChIP-seq profile for the ATF4 TF in the K562 cell line. In this case we observe a single strong peak, at the same location as the ATF3 peak over the *MT-CYB* gene. Unfortunately, we were not able to train a good model for this TF, thus we do not have BPNet predictions over chrM for it (Supplementary Figure 4B). This peak is not seen in a CRISPR-mediated FLAG-tagging experiment in HepG2 cells (Supplementary Figure 4C).

Figure 2D shows the chrM ChIP-seq profile for the ATF7 TF in the MCF-7 cell line. Its profile over chrM is similar to that of ATF3 (Figure 2B), with four peaks. BPNet models corroborate the strong peak over *MT-ND3* /*MT-ND4L*, but are less concordant elsewhere in the genome. These peaks are also seen in GM12878 and K562 ChIP-seq experiments generated with the same antibody (Supplementary Figure 5C-D), but not in HepG2 ChIP-seq carried out with a different antibody (Supplementary Figure 5E).

Figure 3A shows the chrM ChIP-seq profile for the CREB1 TF in the HepG2 cell line. CREB1 is notable for having been previously proposed to localize to mitochondria and play a functional role there ^37,39^, and specifically to bind to the D-loop ^38^. Just as in our previous effort ^26^, we see no evidence that is unlikely to be an artifact for D-loop occupancy, but we observe a strong peak over the *MT-ND1* gene, another one over *MT-CO3* and several weaker others elsewhere in the genome. These match BPNet predictions qualitatively, but the magnitudes of observed and predicted signals differ significantly (Supplementary Figure 6B). The putative occupancy profiles are replicated in MCF-7 cells using the same antibody (Supplementary Figure 6D), in CRISPR FLAG-tagged HepG2 and GM23338 cells (Supplementary Figure 6C and E), and also in K562 cells using a different antibody (Supplementary Figure 6F). However, the latter antibody was also used in datasets in GM12878, H1-hESC and Ishikawa cells (Supplementary Figure 6G-I) resulting in a flat profile over chrM, as is the case with a CETCH-seq experiment in K562 cells (Supplementary Figure 6J).

**Figure 3:**
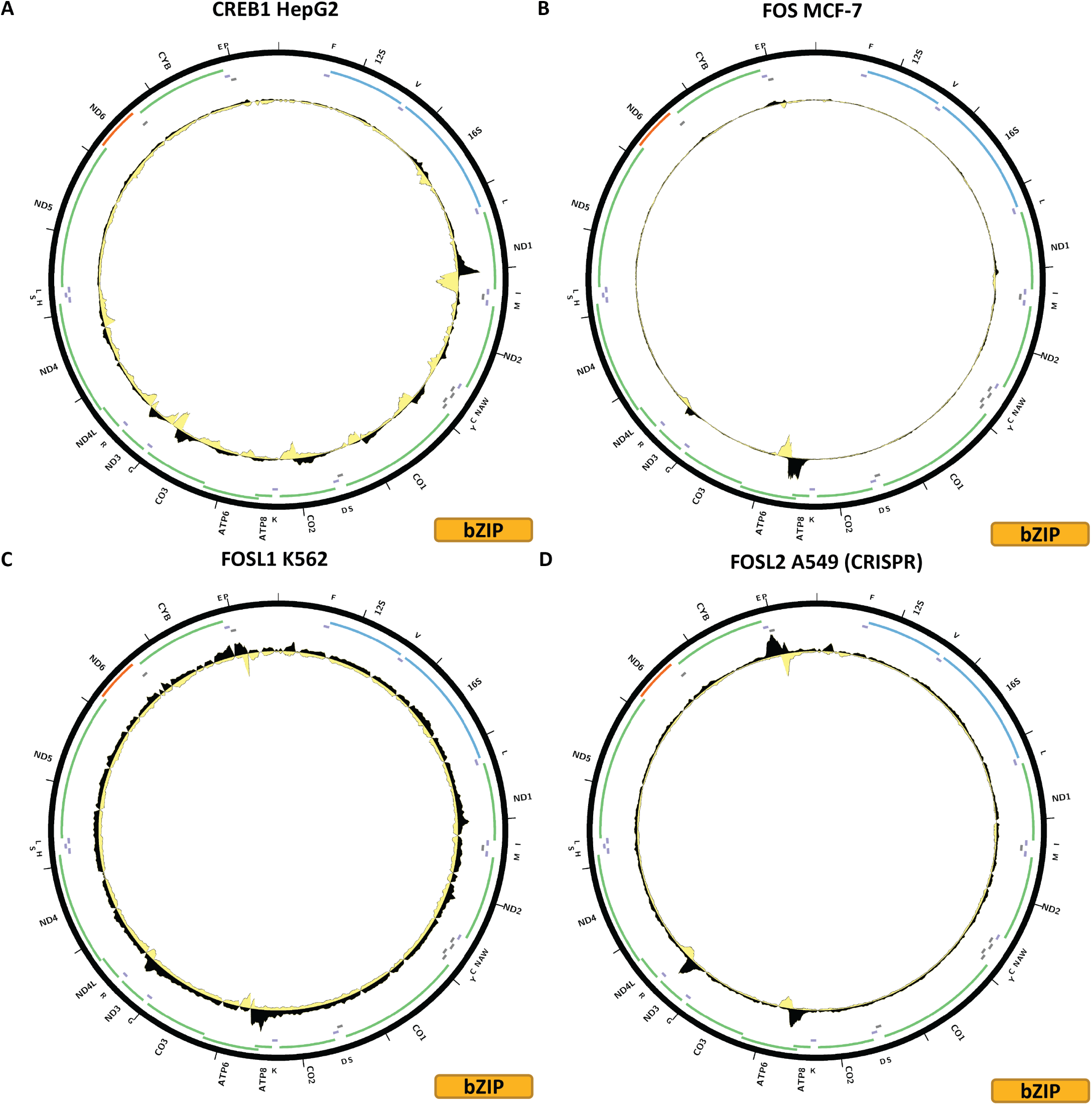
Evidence for mitochondrial genome occupancy by nuclear transcription factors. Black and yellow tracks show the forward- and reverse-strand ChIP-seq coverage over chrM. (A) CREB1 (bZIP); (B) FOS (bZIP); (C) FOSL1 (bZIP); (D) FOSL2 (bZIP).

Figure 3B shows the chrM ChIP-seq profile for the FOS TF in the MCF-7 cell line. Two peaks are observed – a strong one over the *MT-ATP6* gene and a weaker one at the same location as the ATF2, ATF3 and ATF7 peaks over *MT-ND3* /*MT-ND4L*. Both are matched by BPNet predictions while several other BPNet-predicted peaks are not observed in the data (Supplementary Figure 7B). The pattern is replicated in K562 cells using the same antibody (Supplementary Figure 7C), but not in IMR90, endothelial cells of the umbilical vein, GM12878 or HeLaS3, all using the same antibody (Supplementary Figure 7D–G).

Figure 3C shows the chrM ChIP-seq profile for the FOSL1 TF in the K562 cell line; this experiment uses GFP-tagged FOSL1. The same two peaks as for FOS are observed, but neither is particularly strong. BPNet predicts a large number of peaks all over mtDNA, which are not seen in the data (Supplementary Figure 8B). These peaks are not replicated by K562 CETCH-seq, HepG2 CETCH-seq, and ChIP-seq using an *α*-FOSL1 antibody in H1-hESC and HCT116 (Supplementary Figure 8C–F).

Figure 3D shows the chrM ChIP-seq profile for the FOSL2 TF in the A549 cell line, using CRISPR-tagged cells. Again, the same two peaks are observed as for FOS and FOSL1. These are also predicted by BPNet (Supplementary Figure 9B), but the strongest BPNet prediction – over *MT-CYB* is not observed in the data. However, other experiments using two different antibodies in A549, MCF-7, HepG2 and SK-N-SH as well as CETCH-seq in MCF-7 and HepG2 do not shows these peaks (Supplementary Figure 9C–J).

Figure 4A shows the chrM ChIP-seq profile for the CEBPB TF in the K562 cell line, using CRISPR-tagged cells. A strong peak is observed over the *MT-ND4* gene, and a weaker one over *MT-CO2*, as well as a few other weak peaks. These are corroborated by BPNet (Supplementary Figure 10B); in fact BPNet predicts two distinct binding sites over *MT-ND4* and two peaks associated with strand asymmetry are also seen in the CETCH-seq data. BPNet also predicts numerous other peaks that are not observed experimentally. A large number of different additional experiments are available for CEBPB (Supplementary Figure 10C–J and Supplementary Figure 11), using two different antibody lots. The observed putative mtDNA occupancy is replicated in IMR90 (Supplementary Figure 10D), HeLaS3 (Supplementary Figure 10F), HepG2 (Supplementary Figure 10J), A549 (Supplementary Figure 11E), but not in other experiments for HepG2 and A549 or the other cell lines – MCF-7, HCT116, Ishikawa, H1-hESC, GM12878 and non-tagged K562 – that have been assayed.

**Figure 4:**
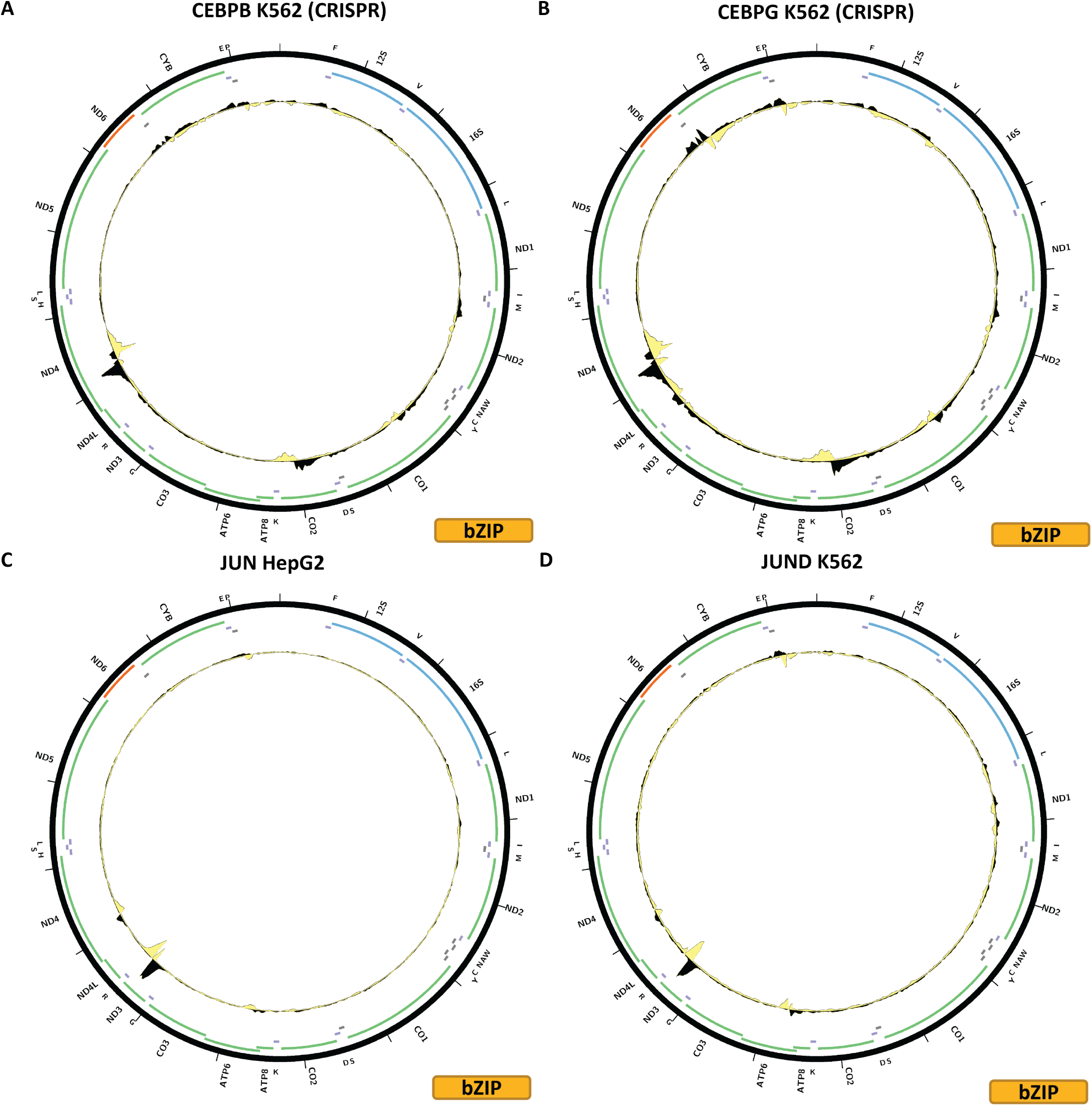
Evidence for mitochondrial genome occupancy by nuclear transcription factors. Black and yellow tracks show the forward- and reverse-strand ChIP-seq coverage over chrM. (A) CEBPB (bZIP); (B) CEBPG (bZIP); (C) JUN (bZIP); (D) JUND (bZIP).

Figure 4B shows the chrM ChIP-seq profile for the CEBPG TF in the K562 cell line, using CRISPR-tagged cells. A similar pattern to CEBPB is observed, but with a stronger peak over the *MT-CYB* gene. BPNet predictions match the observed profile (Supplementary Figure 12B). However, CRISRP-tagged MCF-7 and HepG2 cells do not show these peaks (Supplementary Figure 12C-D).

Figure 4C shows the chrM ChIP-seq profile for the JUN TF in the HepG2 cell line. Two strong peaks are observed for JUN – one over the same *MT-ND3 MT-ND4L* region as seen for many other bZIP factors, and another over *ND4*. A weaker peak is seen over *MT-ATP6*. BPNet predicts all these peaks, as well as many others (Supplementary Figure 13B). These peaks are also observed in endothelial cells of the umbilical veins (Supplementary Figure 13C), where it is to be noted that the *MT-ATP6* peak is stronger than the *MT-ND4L* one, but not in any of the other cell lines assayed – MCF-7, HeLa-S3, A549, K562, H1-hESC and in a HepG2 CETCH-seq sample (Supplementary Figure 13D–I). All of these ChIP-seq experiments were carried out with the same antibody. JUN was one of the factors whose presence in mitochondria was conclusively confirmed through immuno-gold electron microscopy previously ^27^, thus the discrepancy between HepG2 ChIP-seq and CETCH-seq and it being observed only in two seemingly unrelated cell lines and not in any of the others are particularly puzzling observations.

Figure 4D shows the chrM ChIP-seq profile for the JUND TF in the K562 cell line. The same three peaks are observed as for JUN, and these are also corroborated by BPNet predictions (Supplementary Figure 14B). Here too we observed discordance in the available datasets as these peaks are also seen in HepG2 (Supplementary Figure 14D) and SK-N-SH (Supplementary Figure 14H) cells, but not in any of the other ENCODE experiments for JUND – HeLaS3, GM12878, HCT116, H1-hESC, liver, MCF-7, T47D, A549, and most puzzling, additional datasets in K562, HepG2 and SK-N-SH (Supplementary Figure 14C,E-G,I and Supplementary Figure 15). All of these experiments were carried out with the same Santa Cruz Biotech sc-74 antibody, except for the K562 experiments, both of which used GFP-tagged JUND. however, this antibody is polyclonal and there is no information available whether the same lot was used. It is possible the discrepancy arises as a result of lot differences; the other possibility is that the experimental protocols used are not the same as the discordant samples arise from two different production labs. As is the case with JUN, JUND’s presence in mitochondria was previously verified by immunogold electron microscopy in HepG2 cells ^27^.

Figure 5A shows the chrM ChIP-seq profile for the MAFF TF in the K562 cell line. Several peaks are observed – over the *MT-CO1* and *MT-ND5* genes as well as over the tRNA cluster between *MT-ND4* and *MT-ND5*. These are also predicted by BPNet (Supplementary Figure 16B) together with multiple other peaks not observed in the ChIP data. The first two peaks are also observed in HepG2 and HeLa-S3 cells (all experiments carried out with the same antibody) but the latter is not.

**Figure 5:**
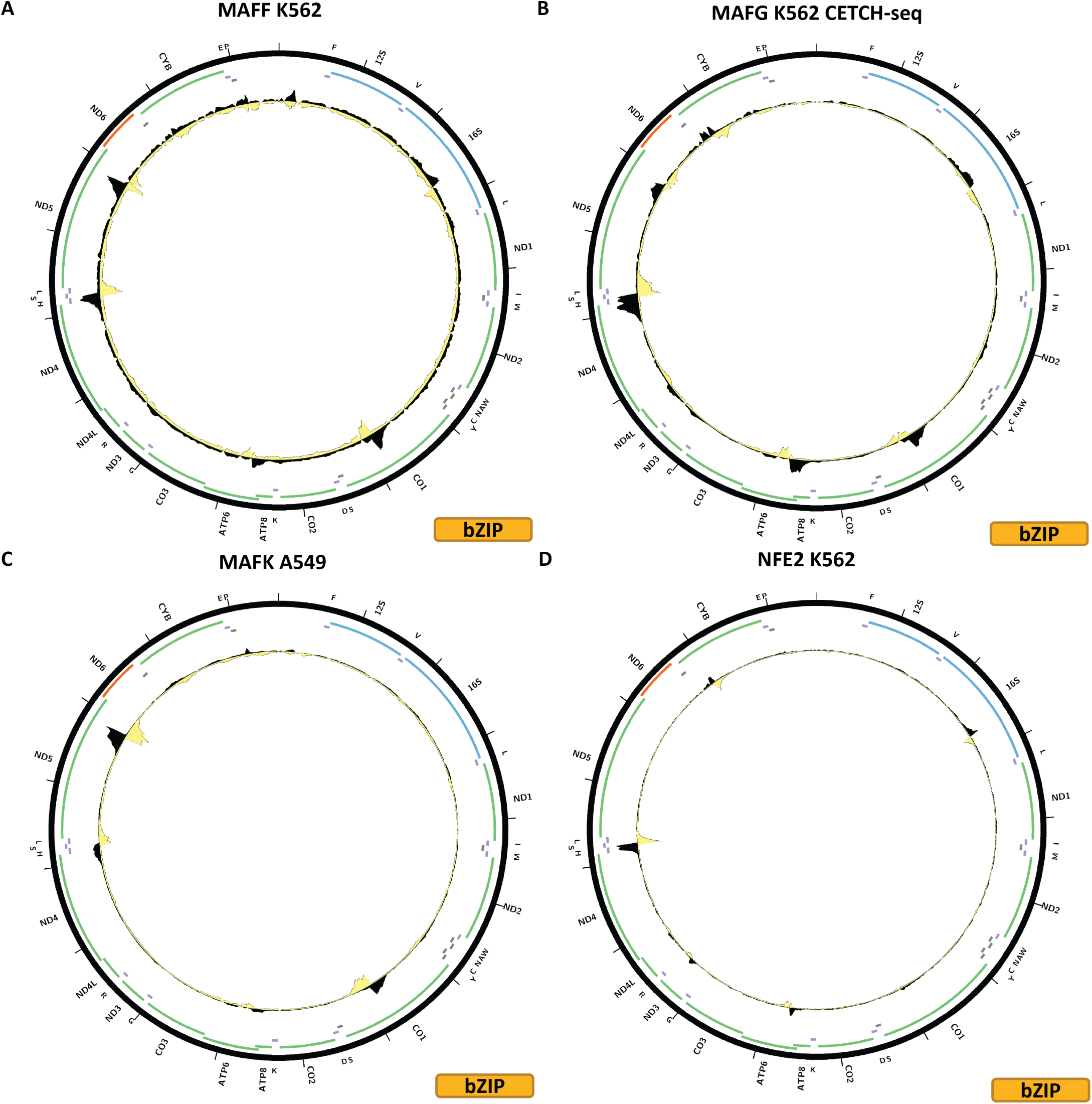
Evidence for mitochondrial genome occupancy by nuclear transcription factors. Black and yellow tracks show the forward- and reverse-strand ChIP-seq coverage over chrM. (A) MAFF (bZIP); (B) MAFG (bZIP); (C) MAFK (bZIP); (D) NFE2 (bZIP).

Figure 5B shows the chrM CETCH-seq profile for the MAFG TF in the K562 cell line. The three peaks observed for MAFF are also present in the MAFG profile, but in addition peaks are present over the *MT-CYB* and *MT-ATP6* genes as well as a weaker one over the 16S rRNA. These peaks are corroborated by BPNet predictions (Supplementary Figure 17B), but are not seen in a HepG2 CETCH-seq experiment (Supplementary Figure 17C).

Figure 5C shows the chrM ChIP-seq profile for the MAFK TF in the A549 cell line. The same peaks are observed as those for MAFF, and they are corroborated by BPNet predictions (Supplementary Figure 18B). They are also observed in GM12878, IMR-90, HeLa-S3, K562, MCF-7 and HepG2 cells, but not in H1-hESC cells (Supplementary Figure 18C–I). Of note, they are seen in datasets generated with two different antibodies, and MAFK was previously shown to localize to mitochondria using immunocytochemical staining.

Figure 5D shows the chrM ChIP-seq profile for the NFE TF in the K562 cell line. Five peaks are observed – over the 16S rRNA gene, over *MT-ATP6*, over *MT-ND3*, over the tRNA cluster between *MT-ND4* and *MT-ND5*, and over *MT-CYB*. These are all sites where peaks are seen also for other bZIP factors. Most of them are predicted by BPNet (Supplementary Figure 19B), and they also seen in K562 CETCH-seq experiment (Supplementary Figure 19C). On the other hand, CETCH-seq in HepG2 (Supplementary Figure 19D) and ChIP-seq in GM12878 generated using a different antibody (Supplementary Figure 19E) show no peaks.

Figure 6A shows the chrM ChIP-seq profile for the NFE2L1 TF in the K562 cell line. Three strong peaks are observed in this case – over the 16S rRNA gene, over *MT-ATP6*, and over the tRNA cluster between *MT-ND4* and *MT-ND5*. BPNet predicts the latter two but not the one over the 16S rRNA (Supplementary Figure 20B). A HepG2 CETCH-seq experiment does not exhibit the same pattern (Supplementary Figure 20C).

**Figure 6:**
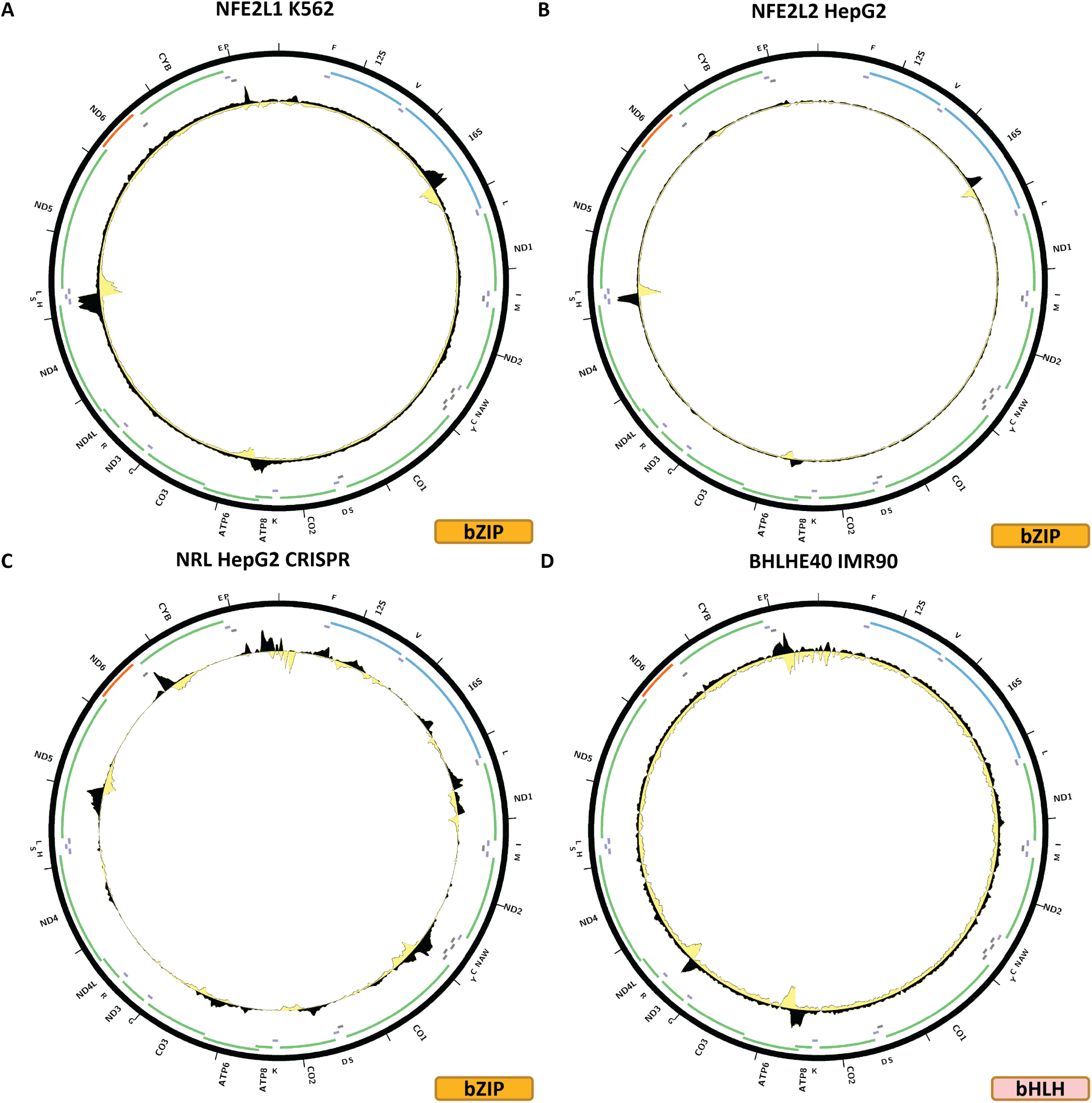
Evidence for mitochondrial genome occupancy by nuclear transcription factors. Black and yellow tracks show the forward- and reverse-strand ChIP-seq coverage over chrM. (A) NFE2L1 (bZIP); (B) NFE2L2 (bZIP); (C) NRL (bZIP); (D) BHLHE40 (bHLH).

Figure 6B shows the chrM ChIP-seq profile for the NFE2L2 TF in the K562 cell line. It is very similar to what is observed for NFE2L1, and in this case too BPNet does not predict a 16S rRNA peak (Supplementary Figure 21B). These peaks are also seen in IMR-90 cells (Supplementary Figure 21C) and weakly in A549 and HeLaS3 cells (Figure 21D–E), using the same antibody for all experiments.

Figure 6C shows the chrM ChIP-seq profile for the NRL TF in a HepG2 CETCH-seq experiment. In this case, multiple, and often potentially complex multisummit peaks are observed all over the genome. They generally match BPNet predictions (Supplementary Figure 22B).

### bHLH TFs

The second major group of TFs exhibiting evidence for mtDNA occupancy are the basic helix–loop–helix (bHLH) transcription factors. Of 122 bHLH factors annotated in the genome, ChIP-seq data is available for 36, of which peaks over chrM are observed for four – BHLHE40, MAX, MITF, and SREBF1.

Figure 6D shows the chrM ChIP-seq profile for the BHLHE40 TF in the K562 cell line. Two peaks are observed – over the *MT-ATP6* gene and over *NT-ND3* /*MT-DN4L*. However, the observed profile does not match the BPNet predicted one (Supplementary Figure 23B), and is also not seen in any other cell line (Supplementary Figure 23C–G). In the GM12878 cell line a different peak is observed over the *MT-ND5* gene (Supplementary Figure 23D); the same antibody was used for both the IMR90 and GM12878 experiments, but a different antibody was used in A549 and HepG2 cells.

Figure 7A shows the chrM ChIP-seq profile for the MAX TF in the K562 cell line. Strong peaks are observed over the 16S rRNA gene and over *MT-CO3*, which are also predicted by BPNet (Supplementary Figure 24B) together with a number of other peaks not seen in the data. Many different additional experiments are available for MAX (Supplementary Figures 24C–J and 25) – all generated with the same antibody, but including experiments from different production groups. These peaks are also seen in endothelial cells of umbilical vein (Supplementary Figure 24F) and H1-hESC (Supplementary Figure 24J), generated by two different productions groups, but not in the rest of the experiments – A549, HepG2 (ChIP and CETCH-seq), GM12878, HCT116, HeLaS3, Ishikawa, liver, MCF-7 and SK-N-SH.

**Figure 7:**
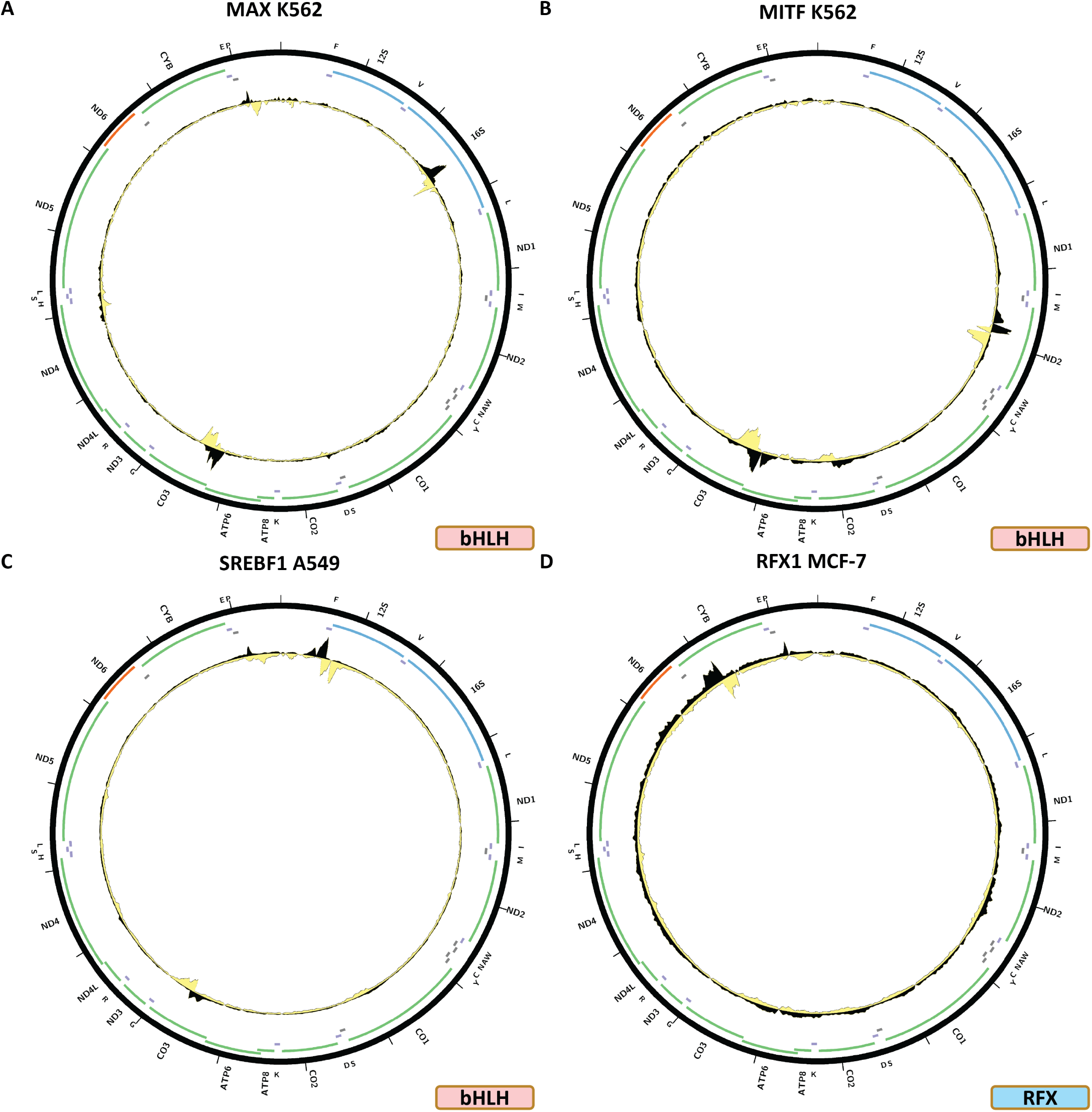
Evidence for mitochondrial genome occupancy by nuclear transcription factors. Black and yellow tracks show the forward- and reverse-strand ChIP-seq coverage over chrM. (A) MAX (bHLH); (B) MITF (bHLH); (C) SREBF1 (bHLH); (D) RFX1 (RFX).

Figure 7B shows the chrM ChIP-seq profile for the MITF TF in the K562 cell line. Peaks are observed over the *MT-ND2* gene and on the boundary between *MT-ATP6* and *MT-CO3*. However, these do not match the predicted BPNet profile (Supplementary Figure 21B).

Figure 7C shows the chrM ChIP-seq profile for the SREBF1 TF in the A549 cell line. In this case, two potential peaks are seen at the very beginning of the 12S rRNA gene and another one over the *MT-C03* gene. These are predicted by BPNet (Supplementary Figure 27B), together with many other peaks not observed in the data.

### RFX TFs

The human genome encodes eight RFX TFs, of which ChIP-seq data now exists for four. Two of these shows evidence for mtDNA occupancy – RFX1 and RFX5.

Figure 7D shows the chrM ChIP-seq profile for the RFX1 TF in the MCF-7 cell line. One peak is observed – over the *MT-CYB* gene – where multiple peaks summits are also predicted by BPNet (Supplementary Figure 28B). This profile is also observed in K562 cells (Supplementary Figure 28C), but not in HepG2 (Supplementary Figure 28D).

Figure 8A shows the chrM ChIP-seq profile for the RFX5 TF in the K562 cell line. Three areas of elevated signal are observed – over *MT-CO2*, in the beginning of *ND5*, and in the *MD-ND3* /*MD-ND4L* region. BPNet predicts multiple strong peaks (Supplementary Figure 29B), two of which match the *MT-CO2* and *MD-ND3* /*MD-ND4L* peaks, but not the *ND5* one. This pattern is also seen in IMR90 cells (Supplementary Figure 29C). In HepG2 cells, a different profile is observed – a peak over the *MT-ND1* gene (Supplementary Figure 29D). No peaks are seen in the available other experiments – SK-N-SH, GM12878, H1-hESC, A549, HeLa-S3, MCF-7 (Supplementary Figure 29E–J), all of which were generated using the same antibody.

**Figure 8:**
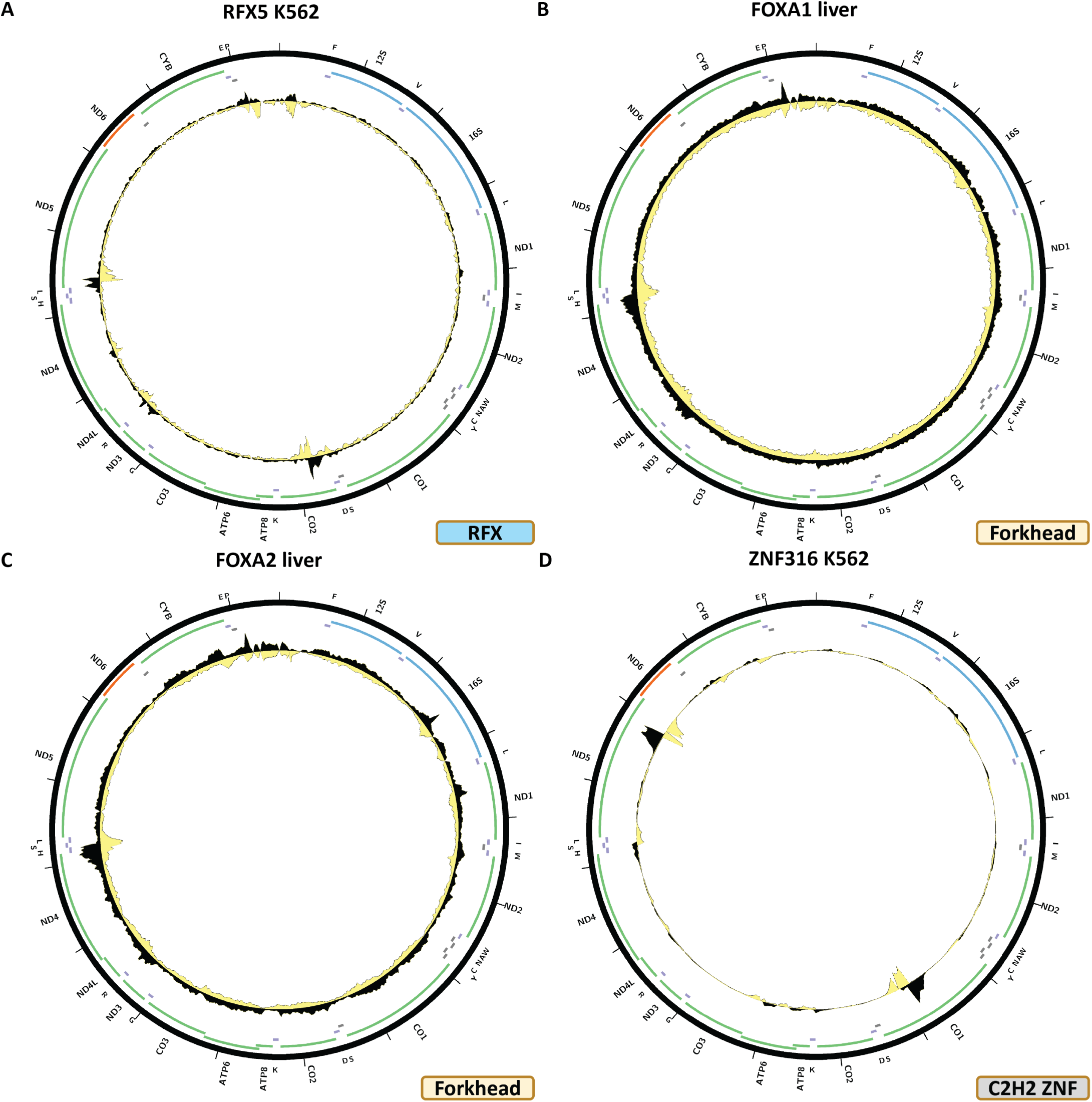
Evidence for mitochondrial genome occupancy by nuclear transcription factors. Black and yellow tracks show the forward- and reverse-strand ChIP-seq coverage over chrM. (A) RFX5 (RFX); (B) FOXA1 (Forkhead); (C) FOXA2 (Forkhead); (D) ZNF316 (C2H2 ZNF).

### Forkhead TFs

A total of 50 forkhead TFs are encoded in the human genome, for which 15 have been assayed by ENCODE. Two of them – FOXA1 and FOXA2 – show putative evidence for mtDNA occupancy.

Figure 8B shows the chrM ChIP-seq profile for the FOXA1 TF in liver. A peak is observed over the tRNA cluster between *MT-ND4* and *MT-ND5*, which is also predicted by BPNet (Supplementary Figure 30B), together with many other peaks not observed in the data. However, HepG2 ChIP-seq carried out with the same antibody does not show the same peak (Supplementary Figure 30D), and neither does HepG2 CETCH-seq (Figure 30E) nor do ChIP-seq datasets for A549, MCF-7 and K562 cells generated using two other different antibodies (Supplementary Figure 30C-D and F-H).

Figure 8C shows the chrM ChIP-seq profile for the FOXA2 TF in liver. The same peak is observed as for FOXA1, but it is not strongly predicted by BPNet (Supplementary Figure 31B). As with FOXA1, it is not seen in cell line data – A549 ChIP-seq, HepG2 CETCH-seq and HepG2 ChIP-seq (Supplementary Figure 31C–E).

### C2H2 zinc finger TFs

The largest TF family in mammals is the C2H2 zinc finger factors. The human genome encodes 774 of these, of which 419 have been now assayed by ENCODE, and 12 show some evidence for mtDNA occupancy – DZIP1, HIVEP1, ZNF225, ZNF274, ZNF350, ZNF598, ZNF768, ZNF839, ZNF891, ZNF263, ZNF280B, ZNF316. Most of these experiments have been carried out using endogenous epitope tagging as specific antibodies for most ZNFs are not available.

Figure 8D shows the chrM ChIP-seq profile for the ZNF316 TF in the K562 cell line. Two strong peaks are observed – over the *MT-CO1* and *MT-ND5* genes, and several weaker ones elsewhere around chrM. These are predicted by BPNet models (Figure 32B). Howevr, a second K562 experiment, carried out by the same production group but with a different antibody does not exhibit any peaks over chrM.

The DZIP1, HIVEP1, ZNF225, ZNF263, ZNF274, ZNF280B, ZNF350, ZNF598, ZNF768, ZNF839, ZNF891, all display a similar pattern over chrM, in, respectively, HepG2 CETCH-seq (Figure 9A), HepG2 CETCH-seq (Figure 9B), HepG2 CETCH-seq (Figure 9C), K562-ChIP-seq (Figure 9D), HepG2 CETCH-seq (Figure 10A), HepG2 CETCH-seq (Figure 10B), HepG2 CETCH-seq (Figure 10C), HepG2 ChIP-seq (Figure 10D), HepG2 CETCH-seq (Figure 11A), HepG2 CETCH-seq (Figure 11B), and HepG2 CETCH-seq Figure 11C). Nine out of eleven of these datasets are the result of endogenous epitope tagging experiments in HepG2 cells, but two are conventional ChIP-seq using TF-specific antibodies. They all display multiple peaks all over the length of the mitochondrial genome, and they are all generally matching the predicted BPNet profiles (Supplementary Figure 33B; Supplementary Figure 34B; Supplementary Figure 35B; Supplementary Figure 36B; Supplementary Figure 37B; Supplementary Figure 38B; Supplementary Figure 39B; Supplementary Figure 40B; Supplementary Figure 41B; Supplementary Figure 42B; Supplementary Figure 43B).

**Figure 9:**
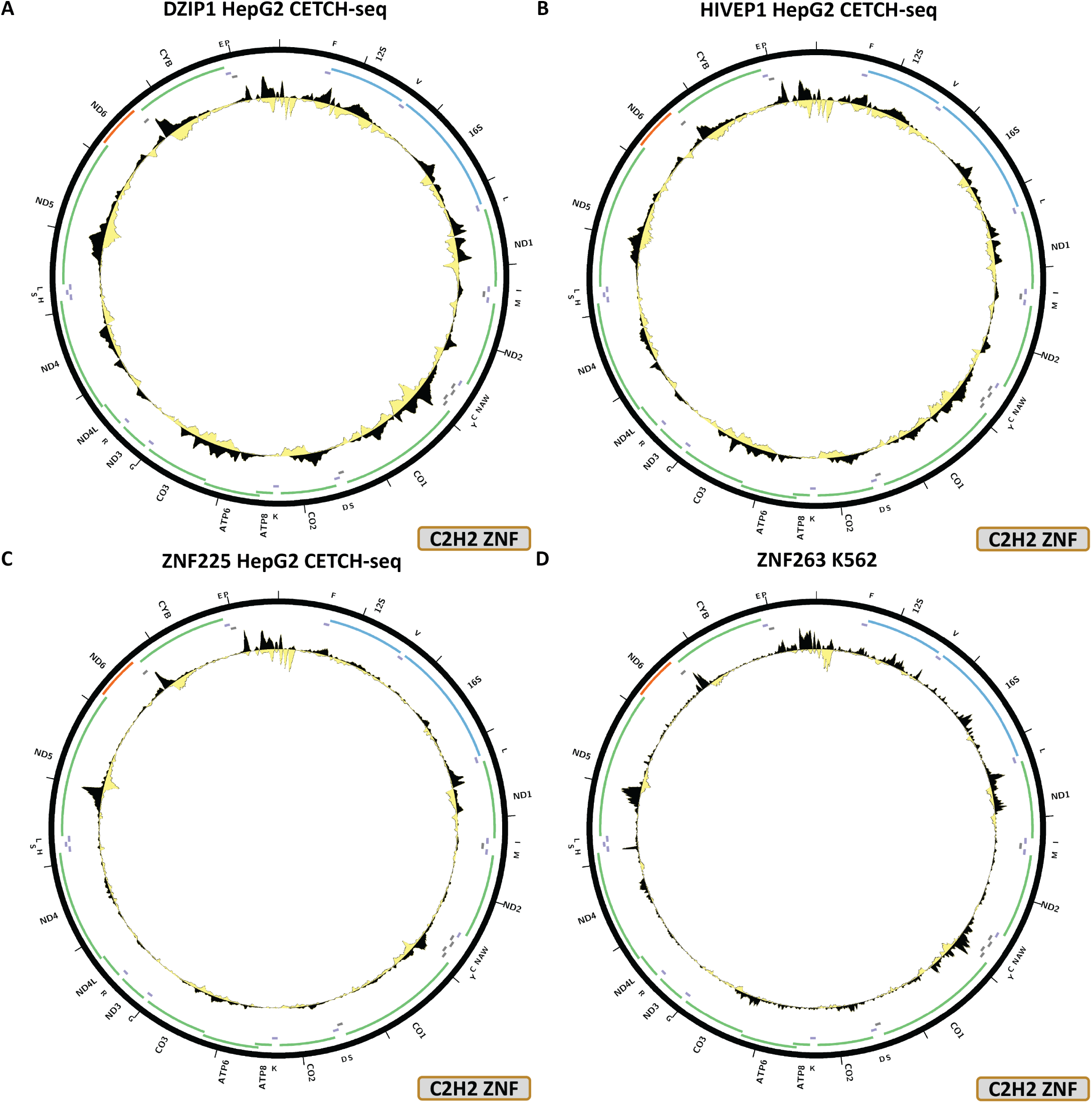
Evidence for mitochondrial genome occupancy by nuclear transcription factors. Black and yellow tracks show the forward- and reverse-strand ChIP-seq coverage over chrM. (A) DZIP1 (C2H2 ZNF); (B) HIVEP1 (C2H2 ZNF); (C) ZNF225 (C2H2 ZNF); (D) ZNF263 (C2H2 ZNF).

**Figure 10:**
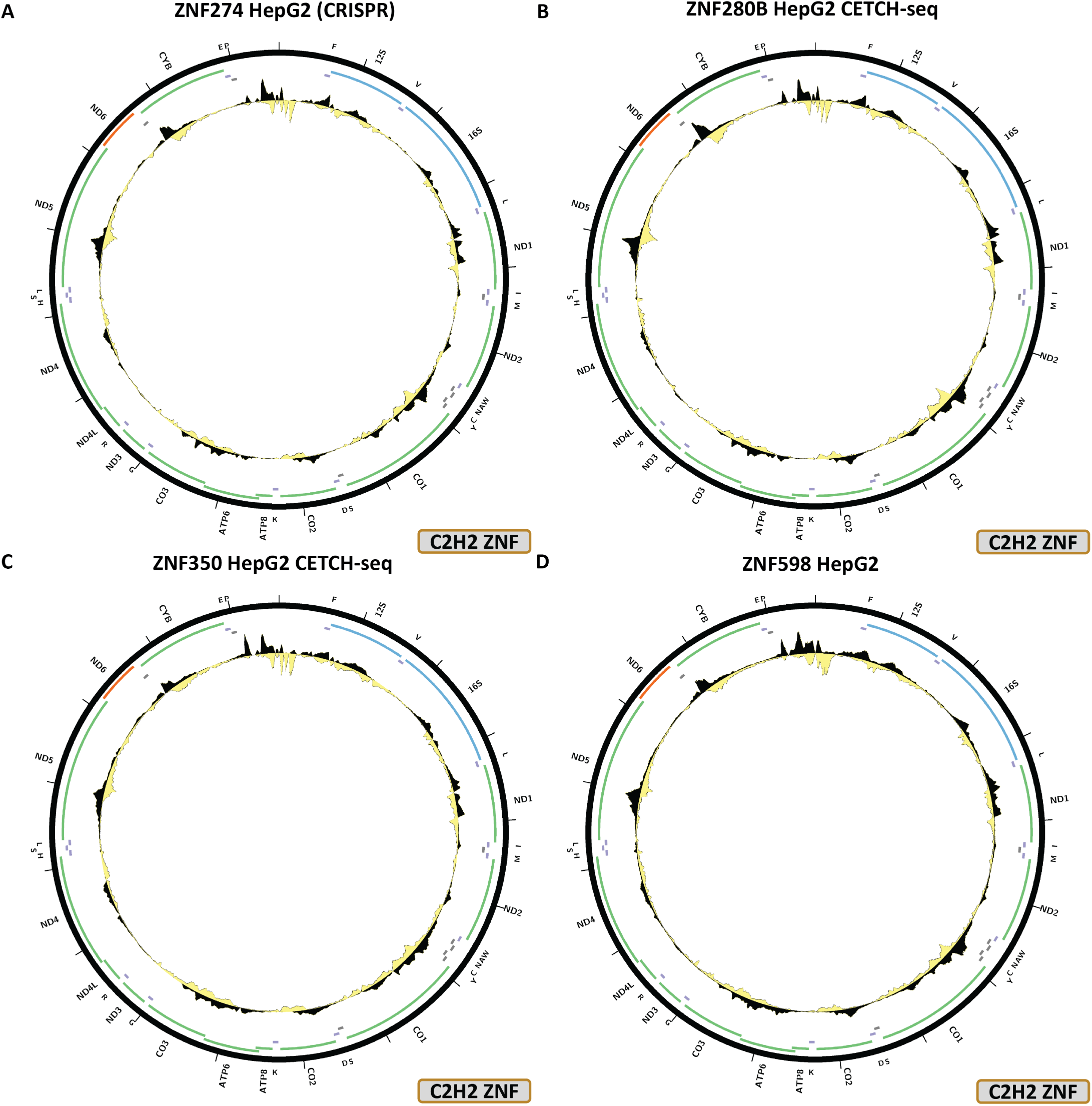
Evidence for mitochondrial genome occupancy by nuclear transcription factors. Black and yellow tracks show the forward- and reverse-strand ChIP-seq coverage over chrM. (A) ZNF274 (C2H2 ZNF); (B) ZNF280B (C2H2 ZNF); (C) ZNF350 (C2H2 ZNF); (D) ZNF598 (C2H2 ZNF).

**Figure 11:**
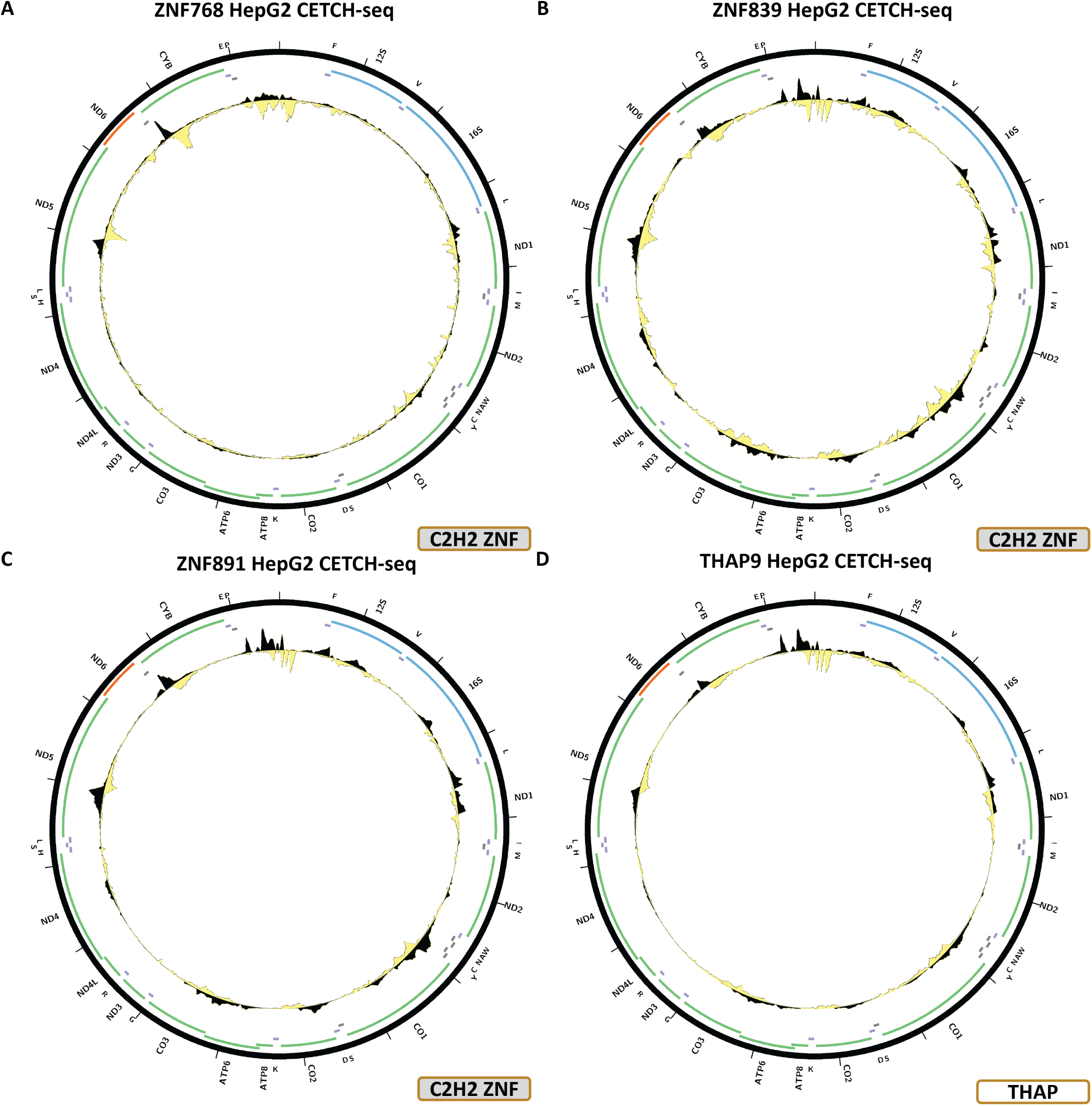
Evidence for mitochondrial genome occupancy by nuclear transcription factors. Black and yellow tracks show the forward- and reverse-strand ChIP-seq coverage over chrM. (A) ZNF768 (C2H2 ZNF); (B) ZNF839 (C2H2 ZNF); (C) ZNF891 (C2H2 ZNF); (D) THAP9 (THAP finger).

For most of these TFs, these are the only datasets available, but in the cases where additional experiments exist, the observed peaks are not replicated – K562 CETCH-seq for HIVEP1 (Supplementary Figure 34C); HEK293 ChIP-seq (Supplementary Figure 35C) and HepG2 CETCH-seq (Supplementary Figure 35D) for ZNF263; HepG2, K562, H1-hESC, HeLa-S3, HCT116, HEK293, and GM12878 ChIP-seq and HEK293 CETCH-seq for ZNF274 (Supplementary Figure 35C–J); HEK293 ChIP-seq for ZNF350 (Supplementary Figure 39C); HEK293 ChIP-seq for ZNF768 (Supplementary Figure 41C).

### THAP finger TFs

The human genome encodes 12 THAP finger TFs, for five of which datasets exist in the ENCODE collection.

Figure 11A shows the chrM CETCH-seq profile for the THAP9 TF in the HepG2 cell line. This factor displays largely the same profile as most of the C2H2 zinc finger TFs discussed above, and it too matches BPNet predictions (Supplementary Figure 44B).

### Rel TFs

Of the ten Rel TFs in the human genome, six have been assayed by ENCODE.

Figure 12A shows the chrM CETCH-seq profile for the NFKB2 TF in the HepG2 cell line. This dataset too exhibits a similar pattern as THAP9 and most of the C2H2 zinc finger TFs, matched by BPNet predictions (Supplementary Figure 45B).

**Figure 12:**
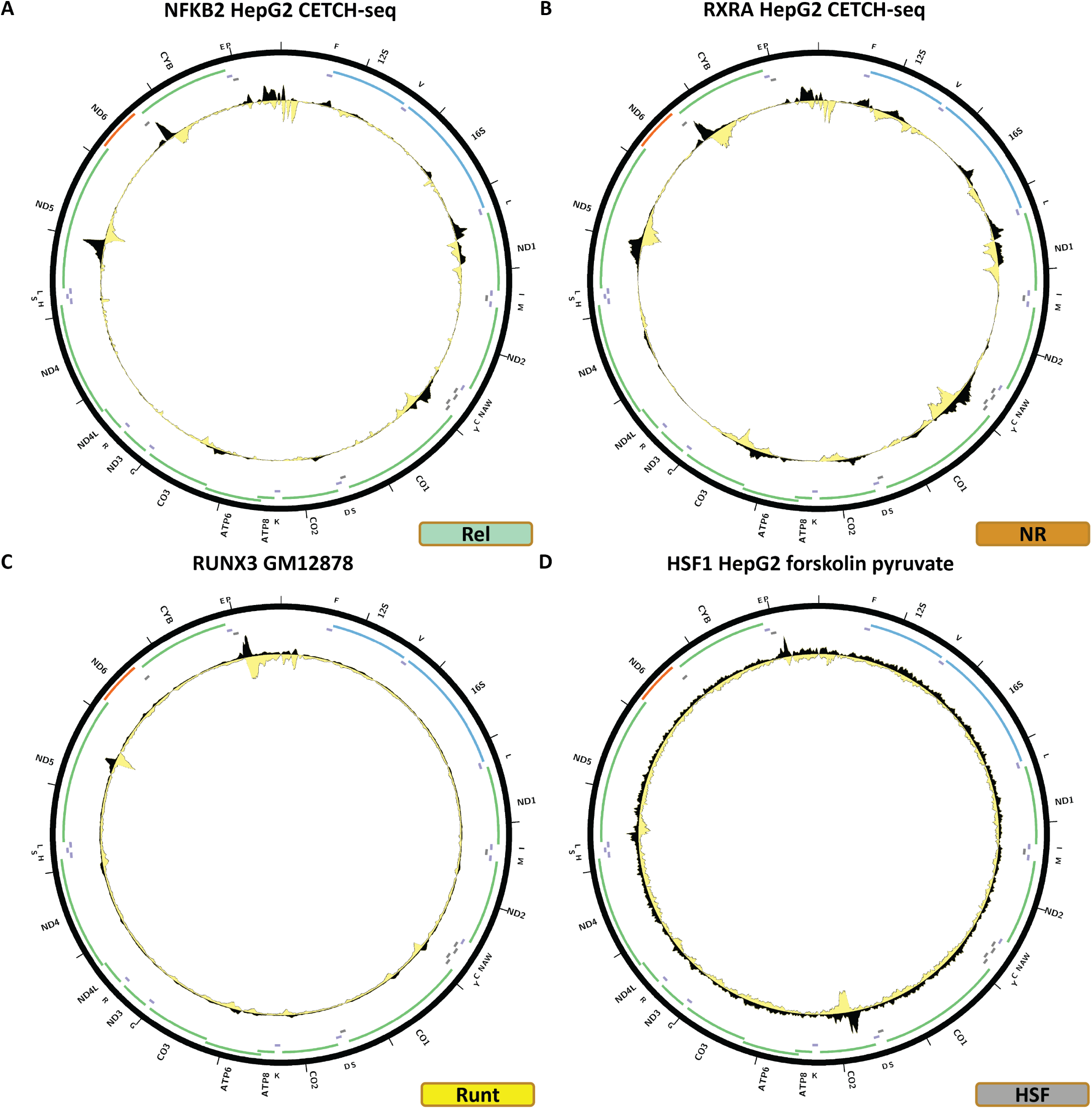
Evidence for mitochondrial genome occupancy by nuclear transcription factors. Black and yellow tracks show the forward- and reverse-strand ChIP-seq coverage over chrM. (A) NFKB2 (Rel); (B) RXRA (nuclear receptor); (C) RUNX3 (Runt); (D) HSF1 (HSF).

### Nuclear receptors

Of the 46 human nuclear receptors, 26 have been assayed by ENCODE.

Figure 12B shows the chrM CETCH-seq profile for the RXRA TF in the HepG2 cell line. This is another case of the same pattern observed for most of the C2H2 zinc finger TFs, THAP9 and NFKB2, and it too largely matches BP-Net predictions (Supplementary Figure 46B). This pattern is not replicated in any of the ChIP-seq datasets available for RXRA in HepG2, GM12878, H1-hESC, liver and SK-N-SH, generated using *α*-RXRA antibodies.

### Runt TFs

Three RUNT TFs are encoded in the human genome. Two of these have been assayed by ENCODE.

Figure 12C shows the chrM ChIP-seq profile for the RUNX3 TF in the GM12878 cell line. One strong peak is observed over the *MT-ND5* gene, two weaker ones over *MT-CO1*, as well as several other loci of slight enrichment. We were not able to train a high quality BPNet model for this TF (Supplementary Figure 47B).

### HSF TFs

ENCODE has assayed three out of eight HSF TFs.

Figure 12D shows the chrM ChIP-seq profile for the HSF1 TF in HepG2 cells treated with forskolin + 1mM pyruvate. One peak is observed over the *MT-CO2* gene, but it is not corroborated by BPNet predictions (Supplementary Figure 48B) and is not replicated in HSF1 ChIP-seq datasets in GM12878 and MCF-7 cells (Figure 48C-D).

### Homeodomain TFs

Homeodomain transcription factors are the second largest class of TFs in mammalian genomes. The human genome encodes 202 of them, plus seven CUT Homeodomain TFs, 16 POU Homeodomain TFs, and nine Paired-box Homeodomain TFs (treated separately in the classification followed here ^1^). We observe evidence for mtDNA occupancy for two Homeodomain TFs and one CUT Homeodomain TF.

Figure 13A shows the chrM ChIP-seq profile for the MEIS2 TF in the K562 cell line. One moderate peak is observed over the *MT-ND1* gene. BPNet predicts elevated signal over that region, but also over many other sites in the mitochondrial genome (Supplementary Figure 49B).

**Figure 13:**
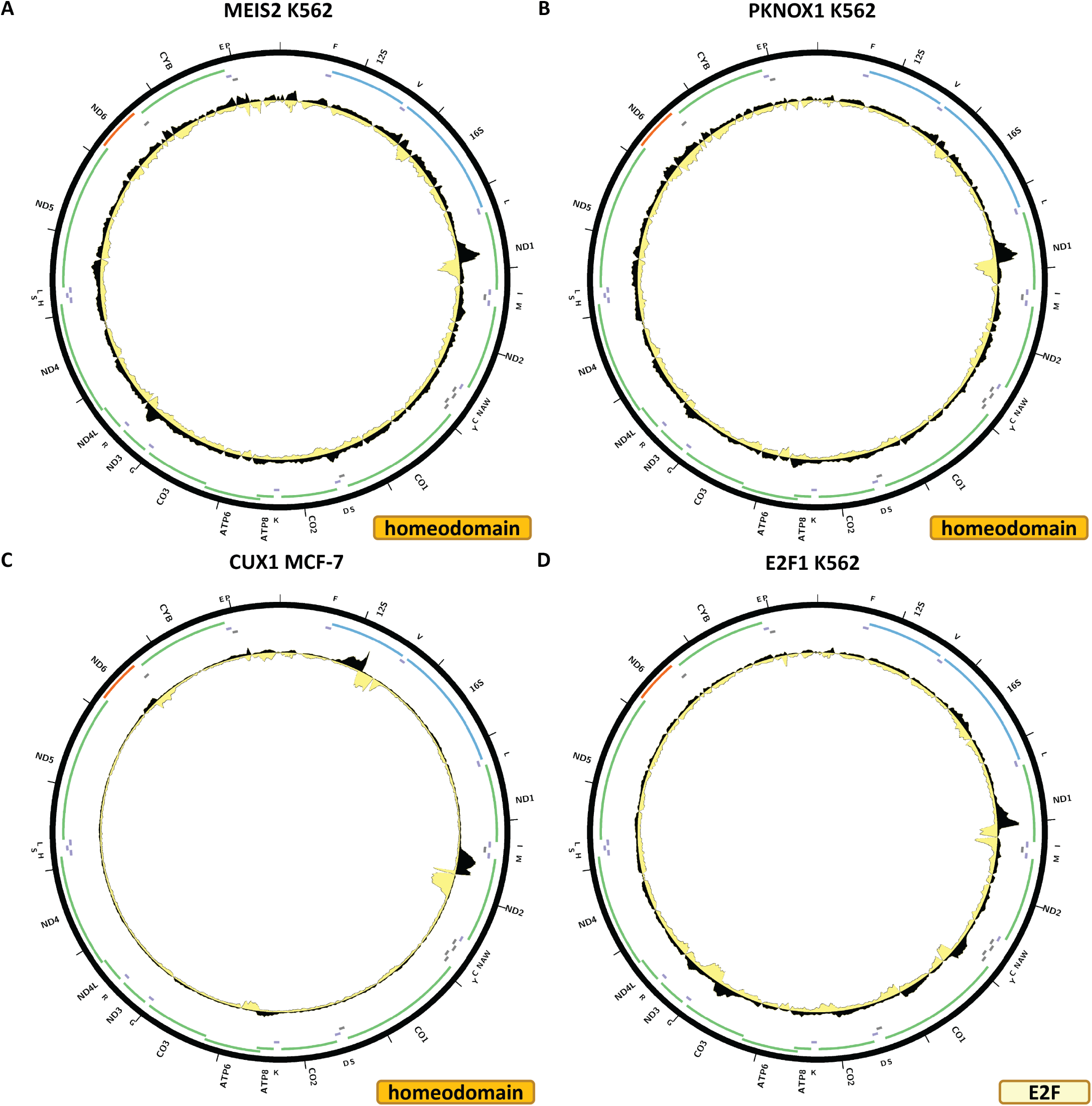
Evidence for mitochondrial genome occupancy by nuclear transcription factors. Black and yellow tracks show the forward- and reverse-strand ChIP-seq coverage over chrM. (A) MEIS2 (Homeodomain); (B) PKNOX1 (Homeodomain); (C) CUX1 (Homeodomain); (D) E2F1 (E2F).

Figure 13B shows the chrM ChIP-seq profile for the PKNOX1 TF in the K562 cell line. The same peak as for MEIS2 is seen, and in this case too BPNet predictions do not match well the observed profile (Figure 50B). The peak is replicated in HEK293T cells (Supplementary Figure 50D), but not in GM12878 (Supplementary Figure 50C) or MCF-7 (Supplementary Figure 50E).

Figure 13C shows the chrM ChIP-seq profile for the CUX1 TF in the MCF-7 cell line. Two strong peaks are observed – over the 12S rRNA gene and over *MT-ND2*, as well as a weaker one over *MT-ATP6*. BPNet predictions include these peaks but also many others, and their relative predicted and observed strengths do not match well (Supplementary Figure 51B). These patterns are replicated in K562 CETCH-seq experiment (Supplementary Figure 51D), but not in K562 ChIP-seq and GM12878 experiments carried out with the same *α*-CUX1 antibody (Supplementary Figure 51C,E).

### E2F TFs

The E2F family consists of 11 TFs in the human genome, for which data is available for ten.

Figure 13D shows the chrM ChIP-seq profile for the E2F1 TF in the K562 cell line. A strong peak is observed over the *MT-ND1* gene, as well as elevated signal around several other loci in the mitochondrial genome. However, this does not match the BPNet-predicted profile (Supplementary Figure 52B), and it is also not replicated in any of the other available datasets – ChIP-seq in K562 cells generated using a different *α*-E2F1 antibody (Supplementary Figure 52C), ChIP-seq in MCF-7 cells carried out using HA-tagged E2F1 (Supplementary Figure 52D), ChIP-seq in HeLaS3 cells carried out using a third *α*-E2F1 antibody (Supplementary Figure 52E), and CETCH-seq experiments in HepG2 and HeLa-S3 cells (Supplementary Figure 52F-G).

### ARID/BRIGHT TFs

Eight out of 15 ARID/BRIGHT TFs in the human genome have been assayed by ENCODE.

Figure 14A shows the chrM ChIP-seq profile for the ARID1B TF in the K562 cell line. A single peak is observed over the *MT-ATP6* gene, but we do not have BPNet support for it (Supplementary Figure 53B), and there are no other datasets available for orthogonal evidence.

**Figure 14:**
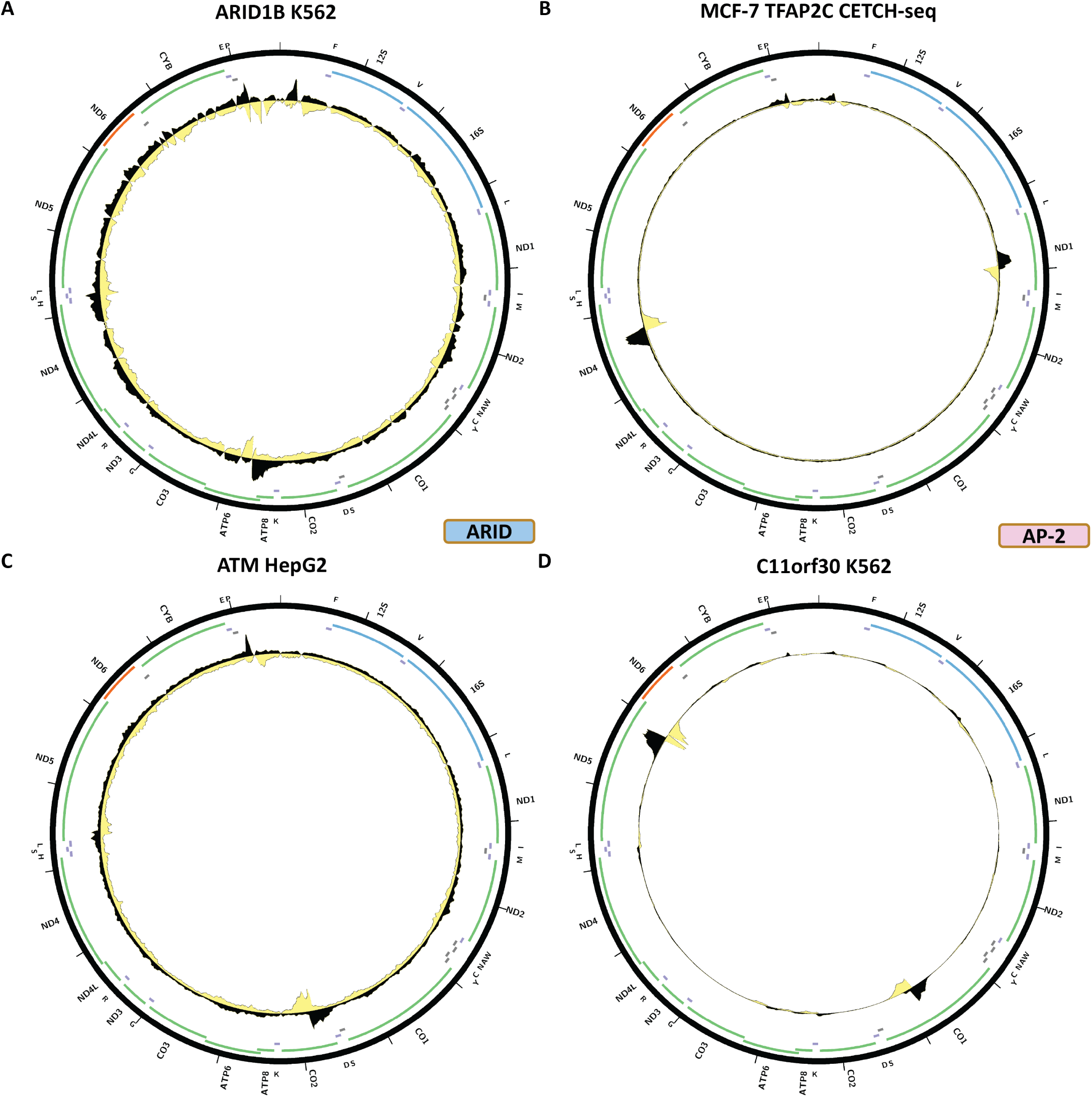
Evidence for mitochondrial genome occupancy by nuclear transcription factors. Black and yellow tracks show the forward- and reverse-strand ChIP-seq coverage over chrM. (A) ARID1B (ARID/BRIGHT) (B) TFAP2C (AP-2); (C) C11orf30; (D) ATM.

### AP-2 TFs

One of the five AP-2 TFs in the human genome has been assayed by ENCODE.

Figure 14B shows the chrM profile for the TFAP2C TF in a CETCH-seq experiment in the MCF-7 cell line. Two very strong peaks are observed – over the *MT-ND1* and *MT-ND4* genes. In this case we were not able to train a good BPNet model, and there are no other experiments available for TFAP2C.

### Other proteins

We did not find any putative chrM peaks for TFs in the following families (Figure 15): AT hook (8/16 assayed), CBF/NF-Y (1/1), CENPB (2/11), CG-1 (1/2), CSD (2/8), CSL (1/2), CxxC (1/11), EBF1 (1/4), Ets (17/28), GATA (7/11), GTF2I-like (1/4), Grainyhead (2/6), HMG/Sox (18/58), IRF (6/9), MADF (1/3), MADS box (1/6), MBD (4/11), Myb/SANT (16/42), NFX (1/2), Ndt80/PhoG (1/2), Pipsqueak (2/2), p53 (1/3), SAND (5/10), SMAD (10/12), STAT (6/7), T-box (5/17), TBP (2/3), TCR/CxC (1/2), TEA (4/5), BED ZF (3/11), CCCH ZF (8/43), and MYM-type ZF (2/16).

**Figure 15:**
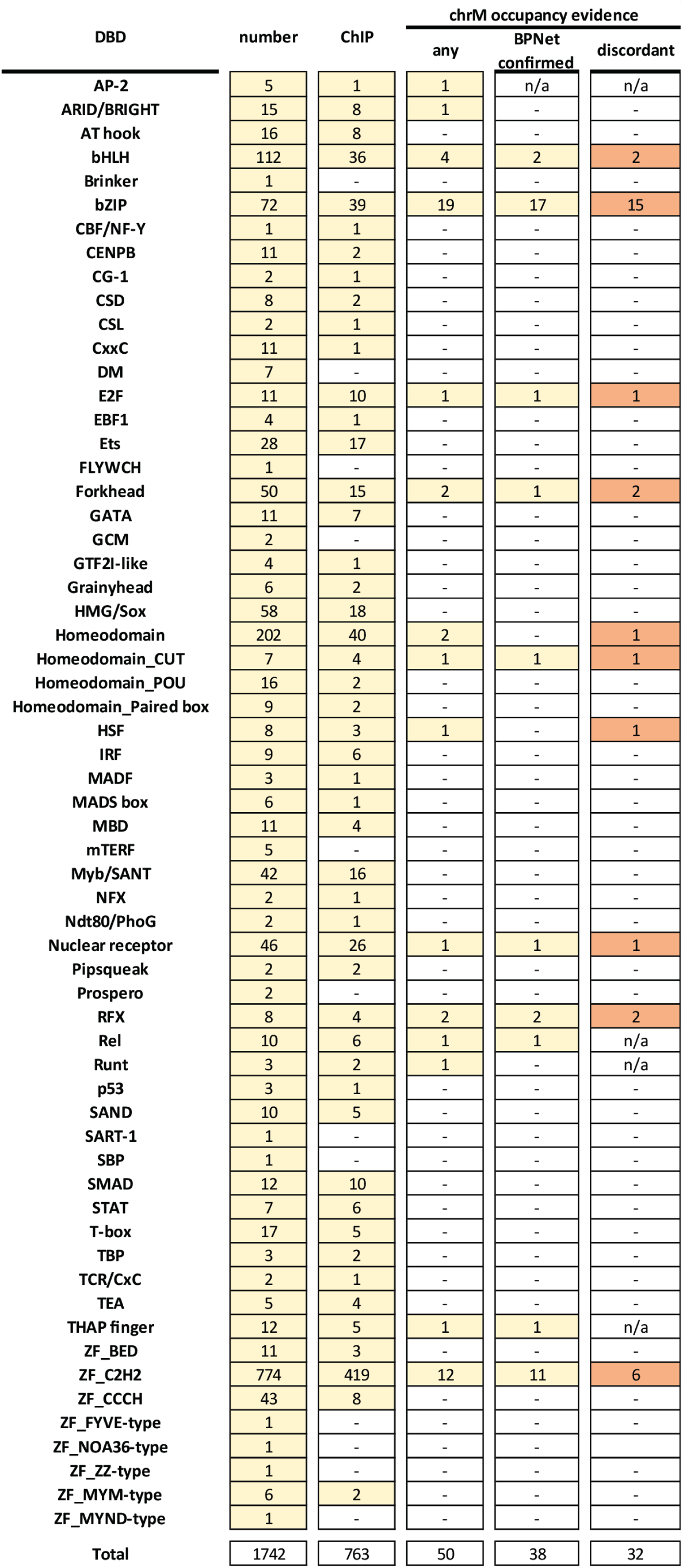
Summary of the available evidence for the physical association of nuclear TFs with the mitochondrial genome. All TFs with ChIP-seq evidence for chrM occupancy are listed in the “any” column. “BPNet confirmed” indicates that the observed ChIP-Seq pattern is corroborated in its key aspects by BP-Net models. The “discordant” TFs are those for which not all available ChIP-seq experiments show chrM peaks.

We observed notable chrM peaks for two other chromatin proteins – C11orf30/EMSY and ATM.

Figure 14C shows the chrM ChIP-seq profile for the C11orf30/EMSY protein in the K562 cell line. Two very strong peaks are observed – over the *MT-CO1* and *MT-ND5* genes. We trained a BPNet model which predicts peaks over a large set of loci, but does include these two peaks (Supplementary Figure 48B). No other datasets are available as orthogonal evidence.

Figure 14D shows the chrM ChIP-seq profile for the ATM TF in the HepG2 cell line. A peak is observed over the *MT-CO2* gene, but the existing BPNet model does not make high-quality predictions over chrM (Supplementary Figure 56B)

## Discussion

In this work we review the current evidence for mtDNA occupancy by nuclear transcription factors using the vastly expanded collection of ChIP-seq datasets generated by the most recent phases of the ENCODE Project Consortium together with interpretable deep neural network modeling of TF occupancy, continuing from our previous work on the subject a decade ago ^26,27^. Some evidence for physical association with mtDNA is found for 50 sequence-specific TFs and two other proteins. However, the interpretation of these observations is less straightforward than it was in the past.

The updated ENCODE collection is qualitatively distinct relative to the much smaller second-phase ENCODE set in that now many TFs have been assayed multiple times, in many different cell lines, and using different combinations of distinct antibodies and/or endogenous epitope tagging. This potentially provides stronger evidence than the more limited data previously available as it can mitigate against the several major concerns that have always existed about observed ChIP-seq peaks over chrM. These are:

1. Whether the experimental protocol used, specifically the fixation step might have involved some kind of permeabilization that allows nuclear TFs to “leak” into the mitochondrial compartment and occupy mtDNA. This should in principle be unlikely if fixation is carried out directly on intact cells, and it would also be expected to result in ChIP-seq profiles showing elevated signal over most of the cognate sequence motifs present in the mitochondrial genome (rather than the one or just a handful of peaks observed in most cases). Still, concerns about such experimental variation are alleviated of mtDNA occupancy is replicated widely across a large number of datasets generated by different productions groups and in different cell lines. On the other hand, the absence of chrM peaks in all cell lines assayed might represent true biological variation.
2. Whether non-specific binding by antibodies, especially polyclonal ones, may be the source of the observed ChIP-seq peaks, i.e. the peaks are real and mediated by some unknown protein that localizes to mtDNA, but it is not the TF that is being assayed that is occupying those sites. Observing the same peaks with multiple different antibodies or a combination of an antibody and endogenous epitope tagging greatly boosts confidence in the physical occupancy of mtDNA by a given TF.
3. Whether epitope tagging affects subcellular localization and/or expression levels. Most of the epitope tagging datasets examined here were generated by C-terminal tagging of the endogenous TF gene. This means that subcellular localization and expression levels should not be affected, but nevertheless such concerns cannot be completely dismissed.

The expanded ENCODE collection provides numerous examples of lack of complete concordance between the different available experiments for each TF. It is thus reasonable to provisionally consider TFs to be likely *in vivo* mtDNA binders if there are at least two orthogonal lines of evidence for their chrM occupancy, i.e. ChIP-seq peaks in datasets generated with at least two different antibodies or an antibody and endogenous tagging, or what is the ideal gold evidential standard – a combination of peaks observed in ChIP-seq datasets and demonstration of localization to mitochondria using immunogold electron microscopy. The latter is however not entirely possible for e.g. many of the C2H2 ZFs, for which epitope tagging was used for ChIP-seq due to the unavailability of immune reagents; in such cases aberrant localization to mitochondria as a result of the tagging cannot be entirely excluded.

With these considerations in mind, we can summarize the available evidence for mtDNA occupancy as follows.

What immediately stands out in the current data is the large number of bZIP factors for which chrM peaks are observed – nearly half (19/39) of the ones that have been assayed. This is unlikely to be an artifact as numerous lines of evidence converge onto bZIP factors playing a role in mitochondria, even though the evidence for each individual TF can be contradictory. For example, ATF2 chrM peaks are seen with multiple antibodies, but not with all or in all epitope tagging experiments; ATF7 peaks are seen in multiple cell lines, but only with one of two antibodies used; FOSL1, FOSL2 and NFE2L1 peaks are not replicated beyond a single dataset, and CEBPG peaks are observed in one epitope-tagged cell line but not in others. On the other hand, CREB1 and MAFK chrM peaks are replicated with multiple different antibodies and NFE peaks in both ChIP-seq and CETCH-seq (although not in all cell lines).

The bZIP factors also include the three TFs for which direct microscopy evidence exists for localization to mitochondria – MAFK, JUN and JUND.

They also exhibit collocalization to a few distinct sites in the mitochondrial genome, which lends mutual support to each other’s mtDNA occupancy because many bZIP factors form heterodimers in the form of the AP-1 transcription factor ^75^. Thus, they would be expected to co-occupy the same sites. AP-1’s nuclear functions also happen to be generally associated with the regulation of growth and proliferation; these are processes in which mitochondria play important roles. While it is currently not clear how AP-1 might be playing a regulatory role in mitochondria mechanistically, this is an obvious functional connection to consider.

On the other extreme of reliability of the available evidence lies the set of C2H2 ZF TFs (DZIP1, HIVEP1, ZNF225, ZNF263, ZNF274, ZNF280B, ZNF350, ZNF598, ZNF768, ZNF839, ZNF891) together with THAP9, NFKB2 and RXRA. All of these datasets exhibit almost the same ChIP-seq profile, are almost all derived from epitope tagging experiments, and they also show elevated signal over nearly all predicted occupancy sites. These peaks are not replicated by any antibody ChIP-seq datasets where available, and thus they are most likely an artifact, although it is not clear why only these CETCH-seq experiments would generate such an artifact and not the hundreds others.

Evidence is also currently weak for mtDNA occupancy by BHLHE40, MITF, FOXA1, FOXA2, HSF1, E2F1, due to lack of replication and/or lack of support from BPNet predictions.

For factors such as MAX, RFX1, RFX5 and PKNOX1 ChIP-seq peaks are seen in multiple cell lines, although not in others. They are provisionally more likely to be truly occupying mtDNA than not.

Yet other factors – SREBF1, RFX1, RUNX3, CUX1, ARID1B, TFAP2C, C11orf30/EMSY and ATM – have only been assayed in a single cell line, and thus the available evidence is simply too limited to say much more about them. Nevertheless, some broad trends emerge. While the bZIP factors appear to be particularly enriched for potential mitochondrial moonlighting, other large and important TF families show very little such evidence. Even if the 12 C2H2 ZFs turn out not to be the result of an experimental artifact, they would represent only a small fraction (12/419) of the huge diversity of such TFs. Other large TF families that, although not exhaustively sampled, don’t seem to bind to chrM include HMG/Sox, nuclear receptors, Homeodomain TFs, and Myb/SANT. A few of the smaller TF families have also been almost exhaustively sampled and they too show no evidence for mitochondrial localization. These include GATA, IRF, SMAD, STAT, and TEA.

In summary, our work represents the most comprehensive catalog of human TFs potentially occupying mitochondrial DNA compiled so far, and provides the foundation for the subsequent direct validation and characterization of the possible functions of these factors in mitochondrial gene regulation.

## Methods

### ChIP-seq data processing

Raw sequencing reads for transcription factor ChIP-seq datasets were downloaded from the ENCODE Consortium Portal ^76^ (https://www.encodeproject.org/; data current as of May 1st 2022). Reads were aligned using Bowtie ^77^ (version 1.1.1) as 1*×*36mers against an index containing the mitochondrial genome, with the following settings ‘‘-v 2 -k 2 -m 1 -t --best --strata’’.

The hg38 version of the *Homo sapiens* genome was used for all analysis.

### Screening for TF occupancy over the mitochondrial genome

We then generated plus- and minus-strand coverage tracks over chrM for all datasets and made Circos ^78^ plots for each such pair. These Circos plots were manually examined to identify likely TF occupancy events and to screen out potentially artifactual high ChIP signal localization events that do not display the expected asymmetric pattern around true occupancy sites.

### Mappability track generation

Mappability was assessed as follows. Sequences of length *N* bases were generated starting at each position in the mitochondrial genome. The resulting set of “reads” was then mapped against the same bowtie index used for mapping real data. Positions covered by *N* reads were considered fully mappable. In this case, *N* = 36 as this is the read length for most of the sequencing data analyzed in this study.

#### BPNet model training and predictions

Uniformly processed ChIP-seq datasets were downloaded from the ENCODE portal. For experiments utilizing paired-end sequencing, PCR duplicates were eliminated. However, for experiments with single-end sequencing, all reads were retained due to the absence of a dependable method for removing duplicates without sacrificing significant genuine signal. Then we generated base resolution signal tracks from the 5’ end of the mapped reads.

The BPNet model architecture and the training approach were adapted from ^70^. The model was designed to accept a one-hot-encoded input DNA sequence of 2,114 base pairs, predicting signals at a 1,000-basepair output window. Additionally, a control input DNA track was also provided to predict the residuals of the ChIP signal from the control track using input sequences. The model outputs comprised of a profile output and a total *log* counts output for 1,000-bp windows.

The model architecture consisted of nine consecutive convolutional layers with ReLU activation. The initial convolutional layer featured a filter size of 21 with no dilation, while subsequent layers utilized filter sizes of 3 with a stride of 1 and increasing dilation rates (power of two) in subsequent layers. Each layer had 64 filters, with residual connections between the convolution layers and zero padding across all layers. Profile prediction was calculated by passing the last dilated convolution output to another convolutional layer with a kernel size of 75 (stride 1) and no padding and no activation, followed by stacking with the control profile tracks and feeding to a final convolutional filter of size 1, operating on one base of the logits and control profiles at a time to generate the predicted profile logits. The *log* of total read counts was computed by performing global average pooling on last dilated convolution output layer. The pooled output was then fed to a dense layer and then concatenated with the *log* of the total counts across both strands from the control experiment, and then further processed through a final dense layer predicting the *log* of the total counts across both strands. The predicted profile logits were converted into probabilities using a single softmax function, generating the profile predictions for both strands, and a negative log-likelihood loss was calculated. Mean squared error of the *log* of total counts across both strands was also computed.

Training epochs comprised of IDR ^79^-thresholded (Irreproducibility Discovery Rate) peaks and non-peak regions (selected to match the GC content of the peak regions) at a 3:1 ratio. Outlier peak regions with signals exceeding 1.2 times the 99th quantile were removed. Additionally, peak sequences were randomly jittered up to 128 bp to augment positive examples. Reverse-complement augmentation was also employed during training. Models were trained and evaluated using 5-fold cross-validation by chromosomes, ensuring no overlap between training, validation, and test sets. Predictions were averaged across both the forward and reverse-complement of the input sequences. All BPNet models and associated outputs will be released as part of the ENCODE Consortium Phase 4 flagship manuscript.

Trained models were then used to generated predicted occupancy profiles for both strands over the mitochondrial genome by splitting the mitochondrial genome into 2,114 bp tiles.

## Data availability

The mitochondrial forward- and reverse-strand tracks analyzed in this study can be found as a UCSC Track Hub at https://mitra.stanford.edu/kundaje/marinovg/oak/public/hubDirectory-chrM/

## Author contributions

G.K.M. conceived the project and carried out data analysis with supervision from A.K. V.R. trained BPNet models and generated BPNet predictions. A.K. and W.J.G. supervised the study. G.K.M. wrote the manuscript with input from all authors.

## Acknowledgments

The authors would like to thank members of the Kundaje and Greenleaf labs for useful comments and discussions. This work was supported by NIH grants 1UM1HG009436 to W.J.G. and A.K., 1DP2OD022870-01 and 1U01HG009431 to A.K, and P50HG007735, RO1 HG008140, U19AI057266 and UM1HG009442 to W.J.G, the Rita Allen Foundation (to W.J.G.), the Baxter Foundation Faculty Scholar Grant, and the Human Frontiers Science Program grant RGY006S (to W.J.G). W.J.G also acknowledges grants 2017-174468 and 2018-182817 from the Chan Zuckerberg Initiative. Fellowship support also provided by the Stanford School of Medicine Dean’s Fellowship (G.K.M.).

## Supplementary Materials

### Supplementary Figures

**Supplementary Figure 1:**
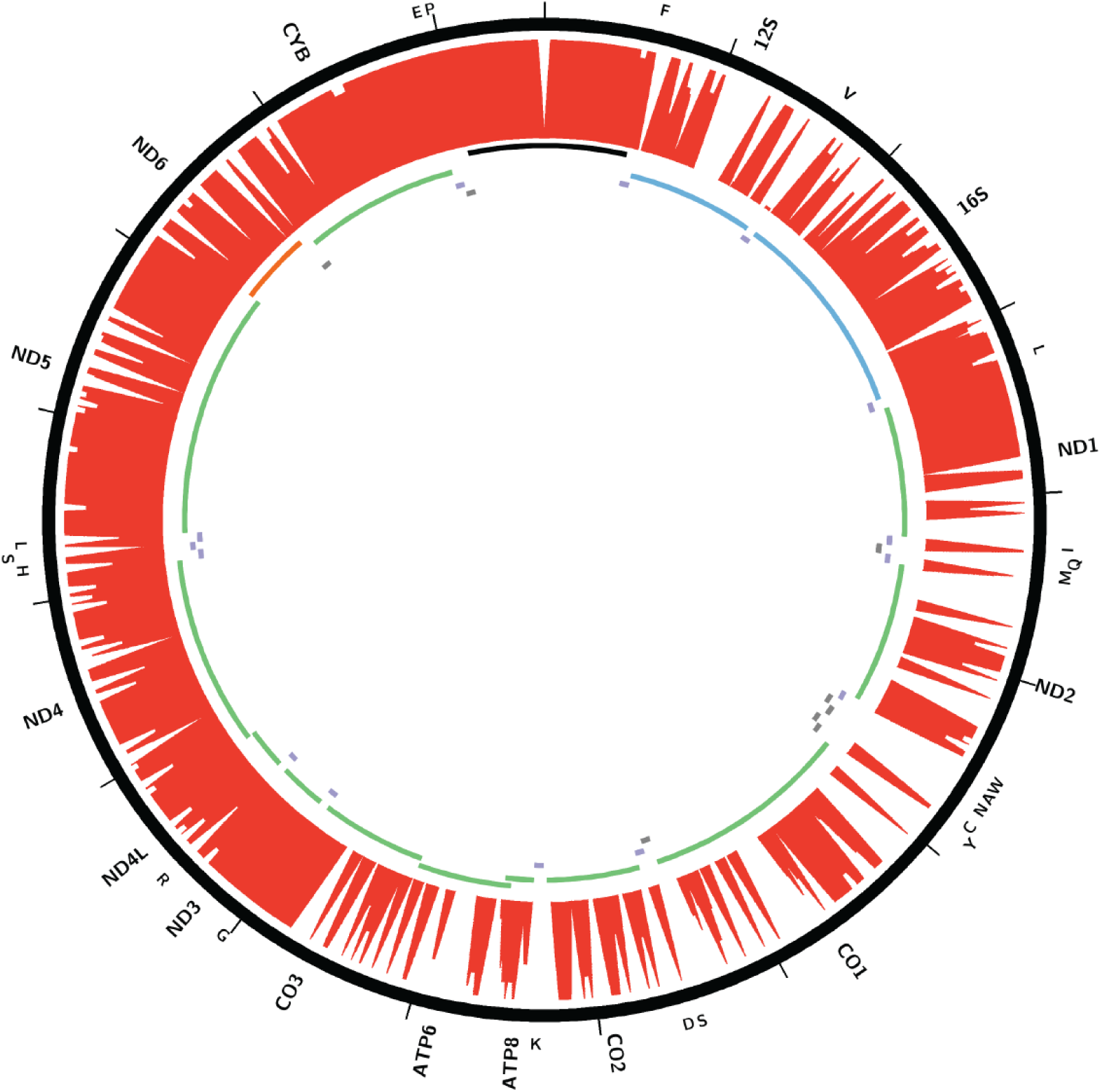
Unique mappability (for 1*×*36mer reads) of the mitochondrial genome in the combined nuclear plus mitochondrial genomic space.

**Supplementary Figure 2:**
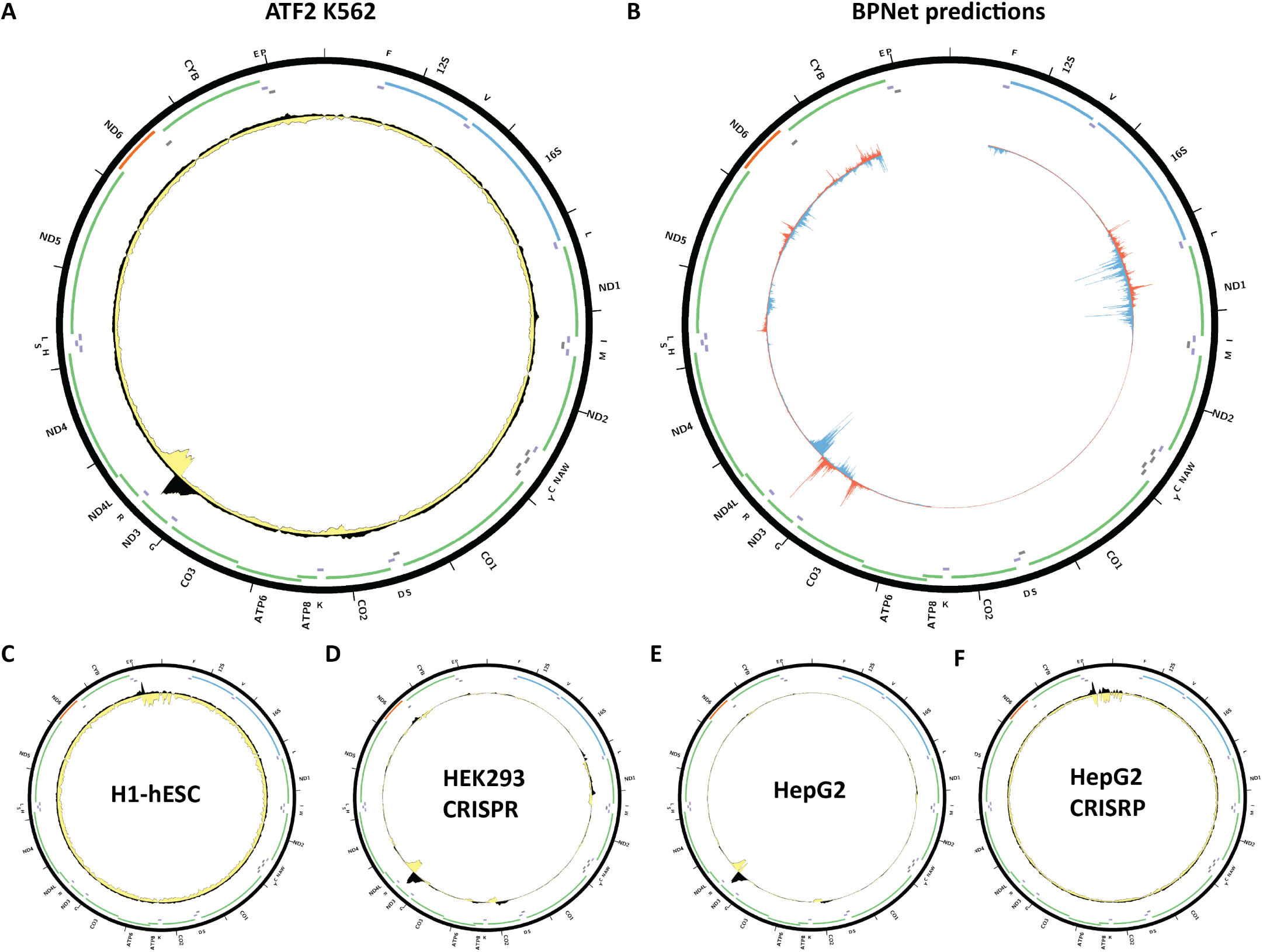
Evidence for mitochondrial genome occupancy by the ATF2 transcription factor. Black and yellow tracks show the forward- and reverse-strand ChIP-seq coverage over chrM. (A) K562 ChIP-seq (ENCODE ID ENCSR869IUD; antibody Bethyl Labs A301-649A). (B) BPNet predictions over chrM (ENCODE ID ENCSR762MDC); (C) H1-hESC ChIP-seq (ENCODE ID ENCSR000BQU; antibody Santa Cruz Biotech sc-81188, Lot ID H0609); (D) HEK293 CETCH-seq (ENCODE ID ENCSR217HTK). (E) HepG2 ChIP-seq (ENCODE ID ENCSR047BUZ; antibody: Bethyl Labs A301-649A); (F) HepG2 CETCH-seq (ENCODE ID ENCSR908HWZ);

**Supplementary Figure 3:**
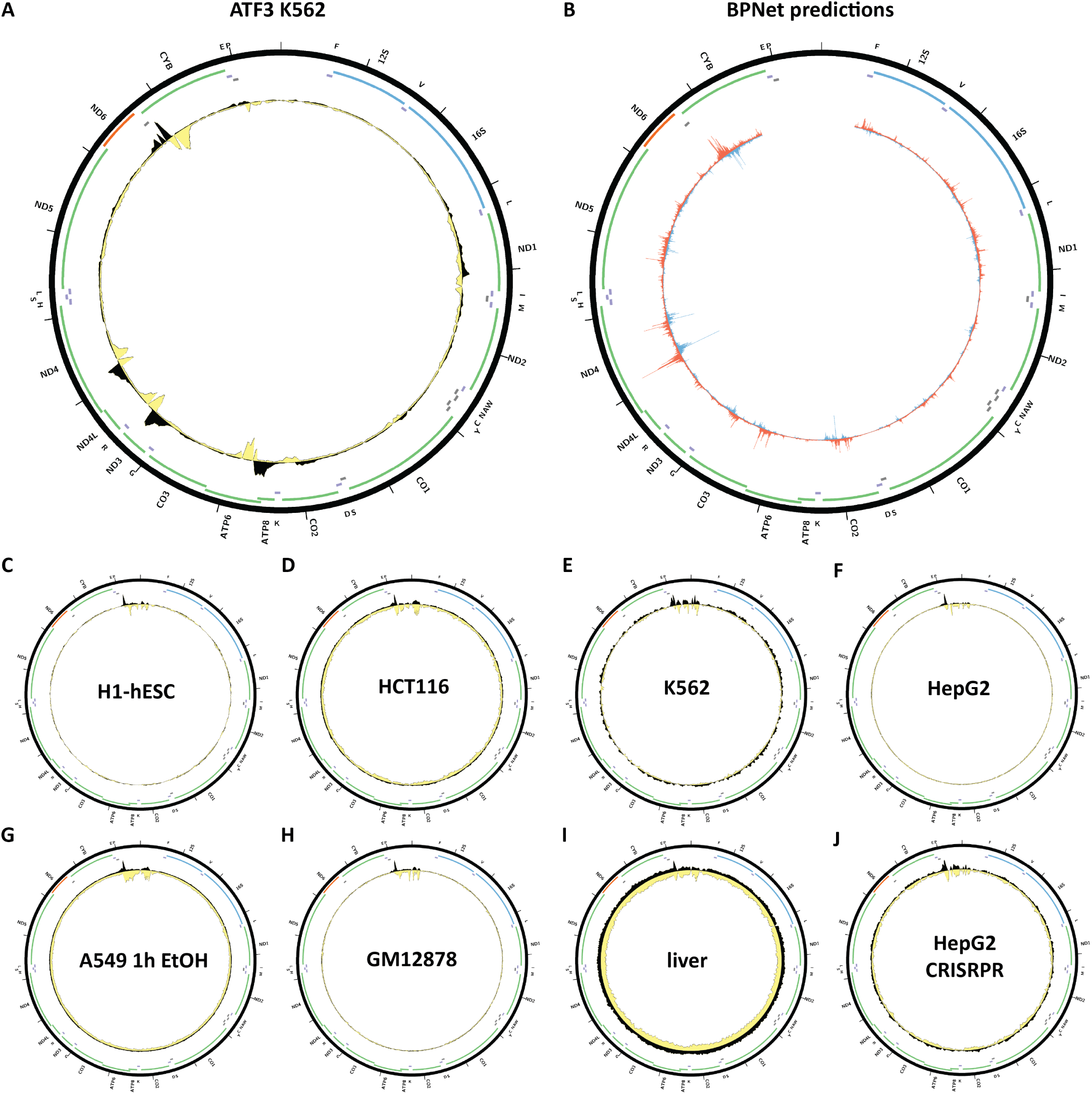
Evidence for mitochondrial genome occupancy by the ATF3transcription factor. Black and yellow tracks show the forward- and reverse-strand ChIP-seq coverage over the mitochondrial genome. (A) K562 ChIP-seq (ENCODE ID ENCSR028UIU; antibody: Active Motif 2/61715); (B) BPNet predictions over chrM (ENCODE ID ENCSR405ECI); (C) H1-hESC ChIP-seq (ENCODE ID ENCSR000BKC; antibody: Santa Cruz Biotech sc-188, Lot ID J2209); (D) HCT116 ChIP-seq (ENCODE ID ENCSR000BUG; antibody: Santa Cruz Biotech sc-188, Lot ID J2209); (E) K562 ChIP-seq (ENCODE ID ENCSR000BNU; antibody: Santa Cruz Biotech sc-188, Lot ID J2209); (F) HepG2 ChIP-seq (ENCODE ID ENCSR000BKE; antibody: Santa Cruz Biotech sc-188, Lot ID J2209); (G) A549 0.02% EtOH 1hChIP-seq (ENCODE ID ENCSR000BPS; antibody: Santa Cruz Biotech sc-188, Lot ID J2209); (H) GM18278 ChIP-seq (ENCODE ID ENCSR000BJY; antibody: Santa Cruz Biotech sc-188, Lot ID J2209); (I) liver ChIP-seq (ENCODE ID ENCSR480LIS; antibody: Santa Cruz Biotech sc-188, Lot ID J2209); (J) HepG2 CETCH-seq (ENCODE ID ENCSR402ZCY).

**Supplementary Figure 4:**
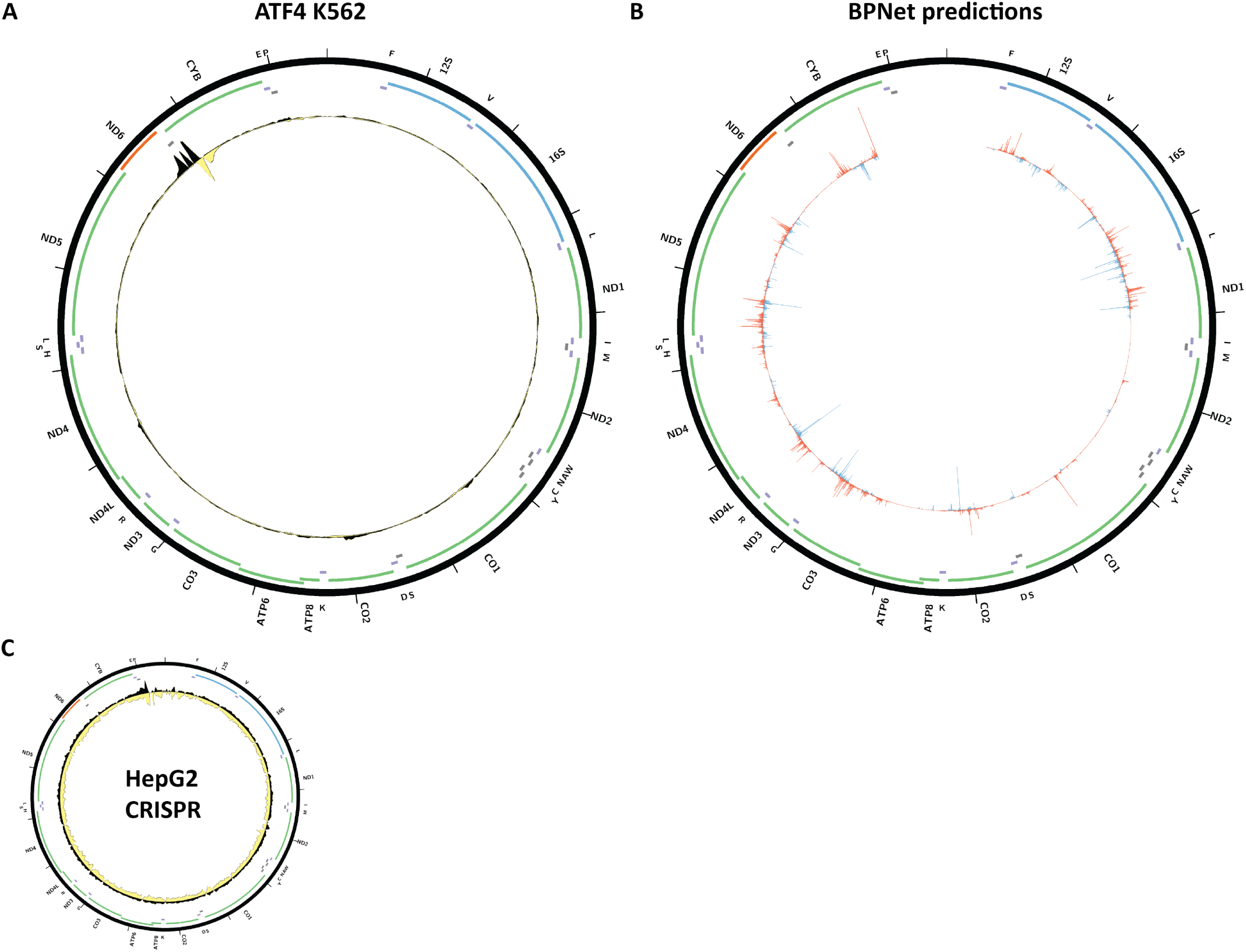
Evidence for mitochondrial genome occupancy by the ATF4 transcription factor. Black and yellow tracks show the forward- and reverse-strand ChIP-seq coverage over chrM. (A) K562 ChIP-seq (ENCODE ID ENCSR145TSJ; antibody: Cell Signaling 11815S); (B) BPNet predictions over chrM (ENCODE ID ENCSR254DFG); (C) HepG2 CETCH-seq (ENCODE ID ENCSR288ZFV).

**Supplementary Figure 5:**
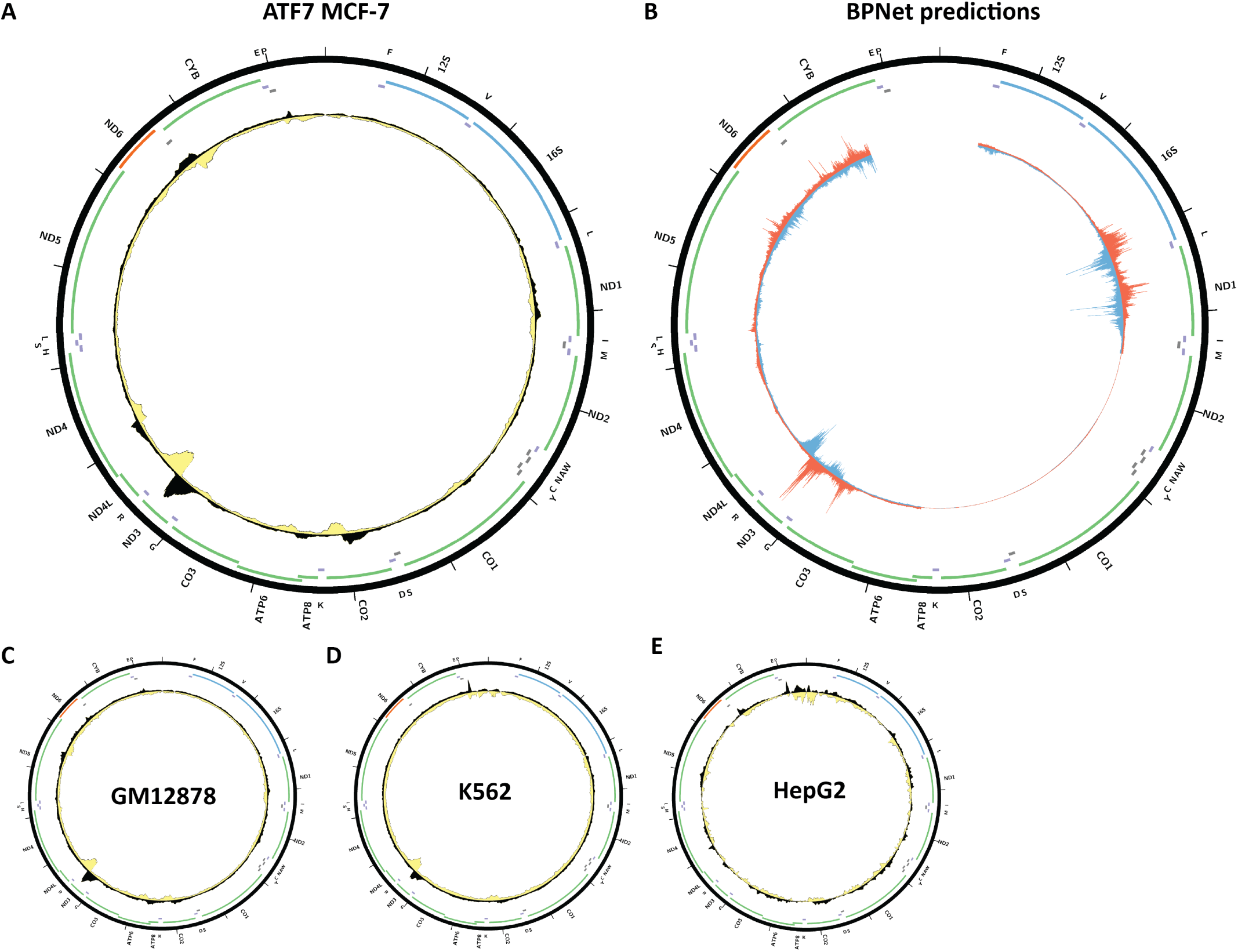
Evidence for mitochondrial genome occupancy by the ATF7 transcription factor. Black and yellow tracks show the forward- and reverse-strand ChIP-seq coverage over chrM. (A) MCF-7 ChIP-Seq (ENCODE ID ENCSR866QPZ; antibody Sigma HPA003384, Lot ID R04563); (B) BPNet predictions over chrM; (C) GM12878 ChIP-seq (ENCODE ID ENCSR014YCR; antibody Sigma HPA003384, Lot ID R04563); (D) K562 ChIP-seq (ENCODE ID ENCSR972ZBV; antibody Sigma HPA003384, Lot ID R04563); (E) HepG2 ChIP-seq (ENCODE ID ENCSR516DDO; antibody: Sigma F1804, Lot ID SLBK1346V).

**Supplementary Figure 6:**
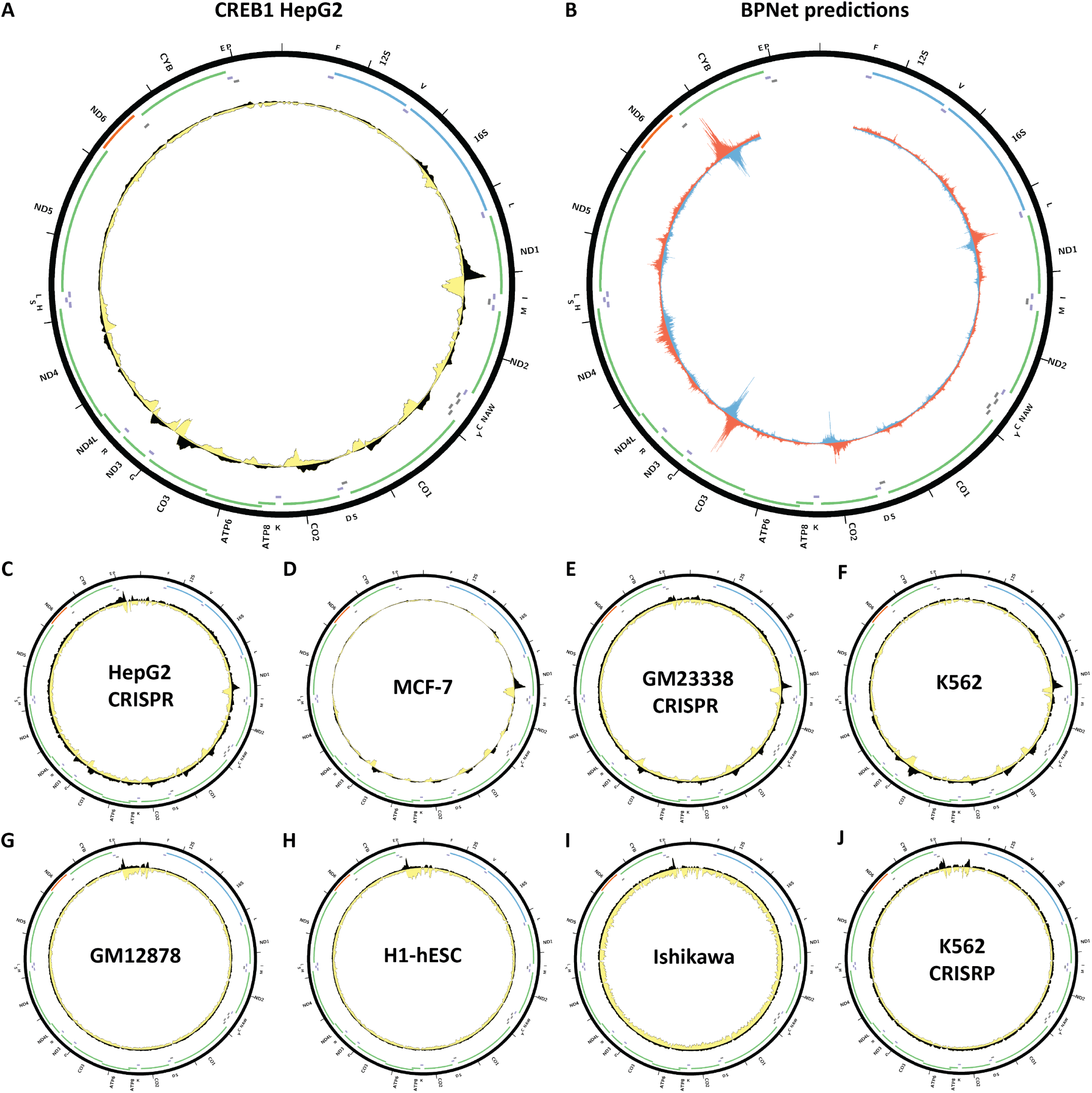
Evidence for mitochondrial genome occupancy by the CREB1 transcription factor. Black and yellow tracks show the forward- and reverse-strand ChIP-seq coverage over chrM. (A) HepG2 ChIP-seq (ENCODE ID ENCSR112ALD; antibody: Cell Signaling 9197S, Lot ID 16); (B) BPNet predictions over chrM (ENCODE ID ENCSR590FMJ); (C) HepG2 CETCH-seq (ENCODE ID ENCSR027JQN); (D) MCF-7 ChIP-seq (ENCODE ID ENCSR897JAS; antibody: Cell Signaling 9197S, Lot ID 16); (E) GM23338 CETCH-seq (ENCODE ID ENCSR214ZAV); (F) K562 ChIP-seq (ENCODE ID ENCSR000BSO; antibody: Santa Cruz Biotech sc-240, Lot ID C2306); (G) GM12878 ChIP-seq (ENCODE ID ENCSR000BUF; antibody: Santa Cruz Biotech sc-240, Lot ID C2306); (H) H1-hESC ChIP-seq (ENCODE ID ENCSR000BSN; antibody: Santa Cruz Biotech sc-240, Lot ID C2306); (I) Ishikawa ChIP-seq (ENCODE ID ENCSR000BUR; antibody: Santa Cruz Biotech sc-240, Lot ID C2306); (F) K562 CETCH-seq (ENCODE ID ENCSR016RFR).

**Supplementary Figure 7:**
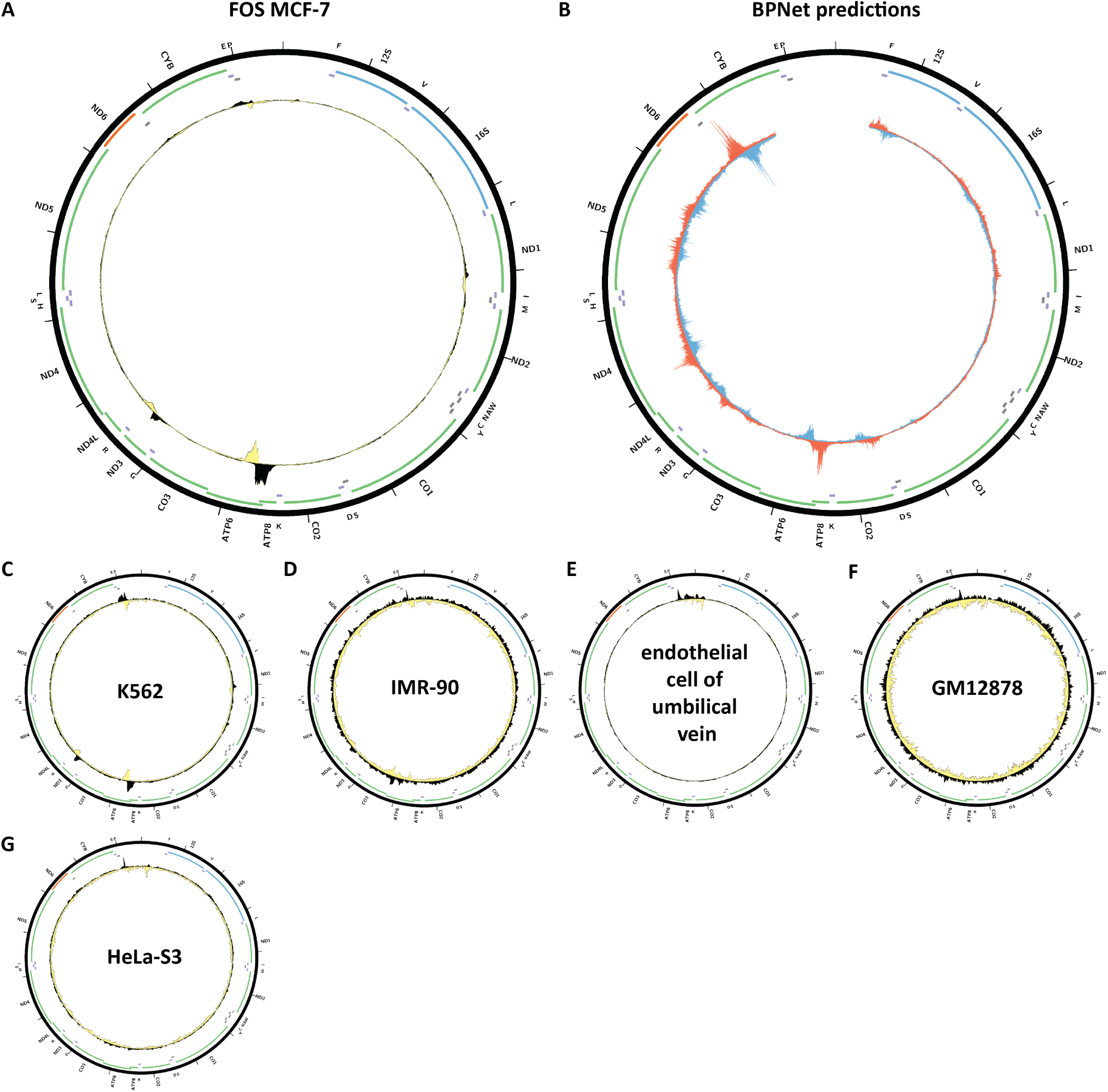
Evidence for mitochondrial genome occupancy by the FOS transcription factor. Black and yellow tracks show the forward- and reverse-strand ChIP-seq coverage over chrM. (A) MCF-7ChIP-seq (ENCODE ID ENCSR569XNP; antibody: Santa Cruz Biotech sc-7202, Lot ID K0810); (B) BPNet predictions over chrM (ENCODE ID ENCSR335UTI); (C) K562 ChIP-seq (ENCODE ID ENCSR000FAI; antibody: Santa Cruz Biotech sc- 7202, Lot ID K0810); (D) IMR-90 ChIP-seq (ENCODE ID ENCSR124AIG; antibody: Santa Cruz Biotech sc-7202, Lot ID K0810); (E) endothelial cell of umbilical vein ChIP-seq (ENCODE ID ENCSR000EVU; antibody: Santa Cruz Biotech sc-7202, Lot ID K0810); (F) GM12878 ChIP-seq (ENCODE ID ENCSR000EYZ; antibody: Santa Cruz Biotech sc-7202, Lot ID K0810); (G) HeLa-S3 ChIP-seq (ENCODE ID ENCSR000EZE; antibody: Santa Cruz Biotech sc-7202, Lot ID K0810).

**Supplementary Figure 8:**
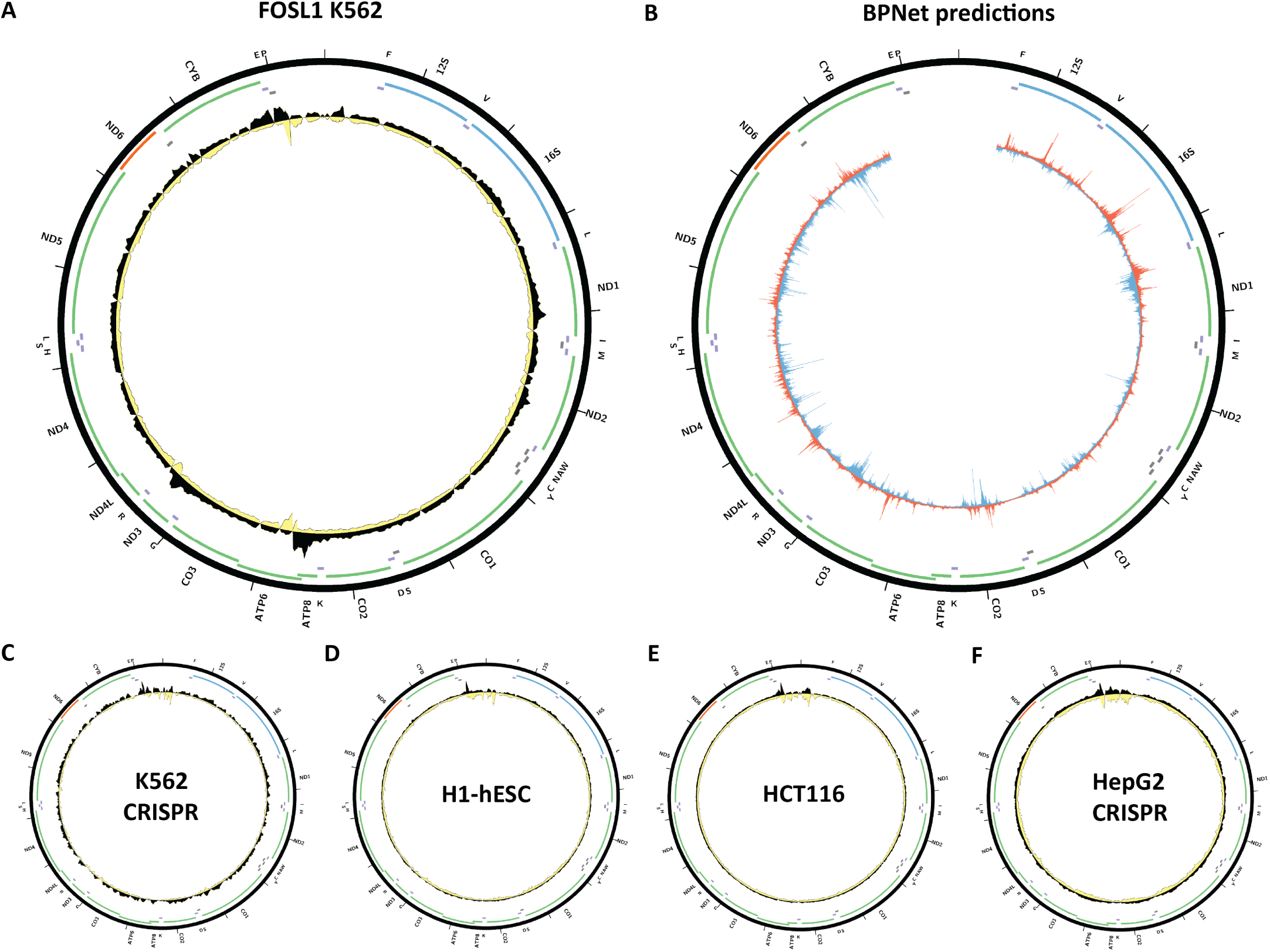
Evidence for mitochondrial genome occupancy by the FOSL1transcription factor. Black and yellow tracks show the forward- and reverse-strand ChIP-seq coverage over chrM. (A) K562 ChIP-seq (ENCODE ID ENCSR239ZLZ; GFP-tagged); (B) BPNet predictions over chrM (ENCODE ID ENCSR285NIO); (C) K562 CETCH-seq (ENCODE ID ENCSR000BMV); (D) H1-hESC ChIP-seq (ENCODE ID ENCSR000BNS; antibody: Santa Cruz Biotech sc-183, Lot ID I0809); (E) HCT116 ChIP-seq (ENCODE ID ENCSR000BTE; antibody: Santa Cruz Biotech sc-183, Lot ID I0809); (F) HepG2 CETCH-seq (ENCODE ID ENCFF660RIA).

**Supplementary Figure 9:**
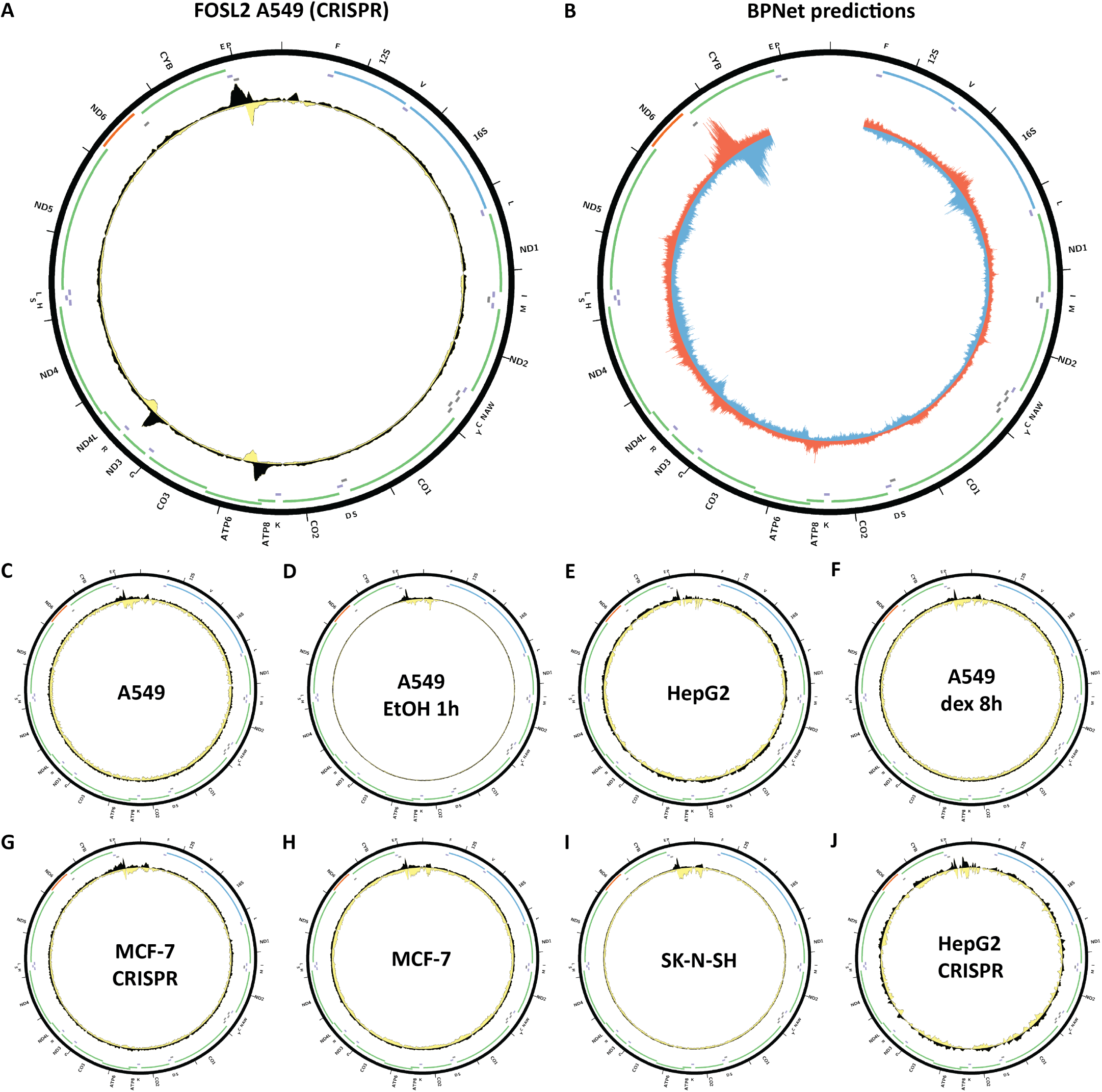
Evidence for mitochondrial genome occupancy by the FOSL2transcription factor. Black and yellow tracks show the forward- and reverse-strand ChIP-seq coverage over chrM. (A) A549 CETCH- seq (ENCODE ID ENCSR448TVS); (B) BPNet predictions over chrM (ENCODE ID ENCSR554ZTS); (C) A549 ChIP-seq (ENCODE ID ENCSR593DGU; antibody: Santa Cruz Biotech sc-160, Lot ID H2713); (D) A549 EtOH 1 hour ChIP-seq (ENCODE ID ENCSR000BQO; antibody: Santa Cruz Biotech sc-604); (E) HepG2 ChIP-seq (ENCODE ID ENCSR000BHP; antibody: Santa Cruz Biotech sc-604); (F) A549 Dex 8 hours ChIP-seq (ENCODE ID ENCSR242EWU; antibody: Santa Cruz Biotech sc-160, Lot ID H2713); (G) MCF-7 CETCH-seq (ENCODE ID ENCSR546KCN); (H) MCF- 7 ChIP-seq (ENCODE ID ENCSR000BUI; antibody: Santa Cruz Biotech sc-604); (I) SK-N-SH ChIP-seq (ENCODE ID ENCSR000BVB; antibody: Santa Cruz Biotech sc-604); (J) HepG2 CETCH-seq (ENCODE ID ENCSR249EYB).

**Supplementary Figure 10:**
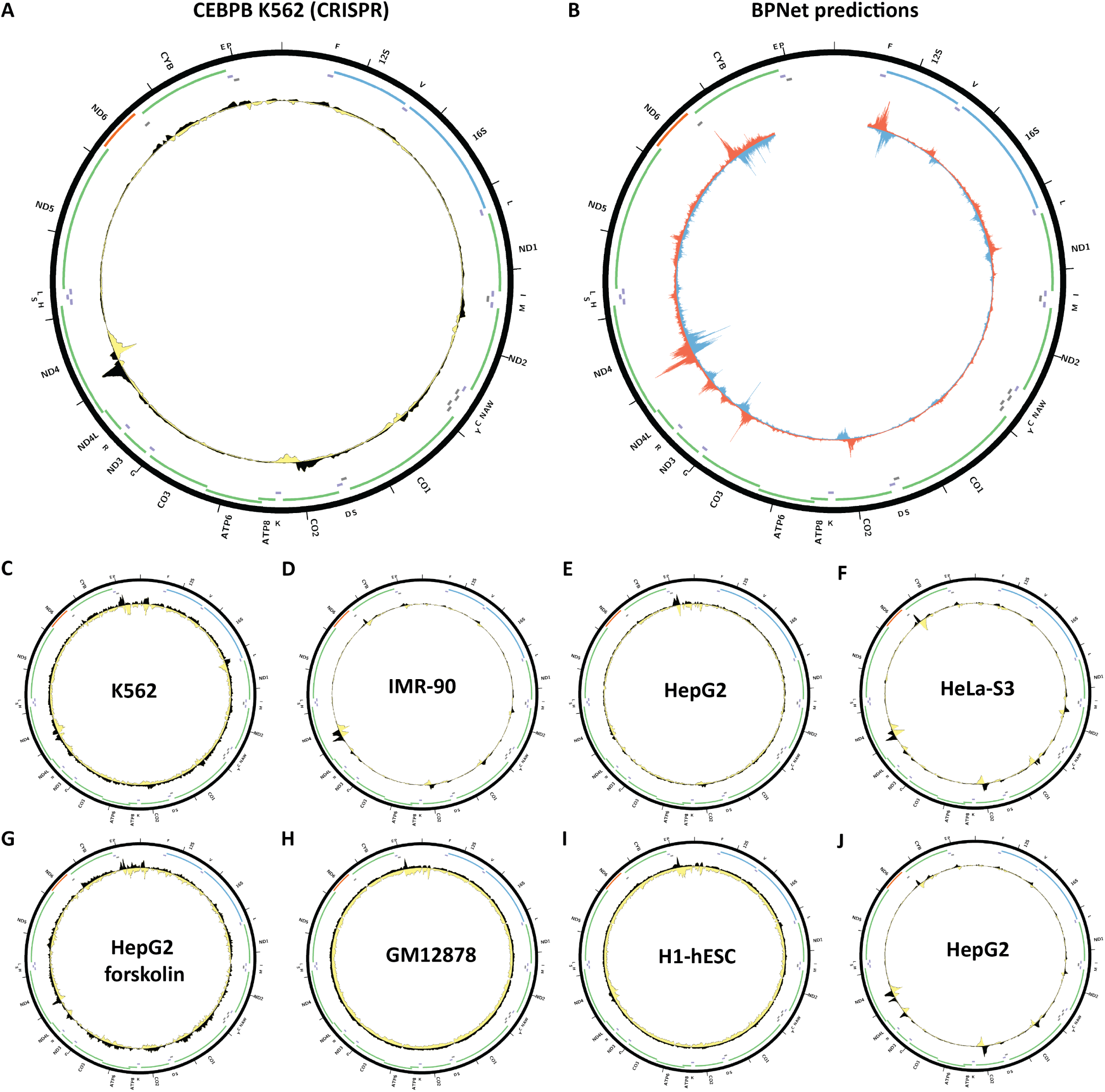
Evidence for mitochondrial genome occupancy by the CEBPB transcription factor. Black and yellow tracks show the forward- and reverse-strand ChIP-seq coverage over chrM. (A) K562CETCH-seq (ENCODE ID ENCSR416QLJ); (B) BPNet predictions over chrM (ENCODE ID ENCSR700UVP); (C) K562 ChIP-seq (ENCODE ID ENCSR000EHE; antibody: Santa Cruz Biotech sc-150, Lot ID I1010); (D) IMR-90 ChIP-seq (ENCODE ID ENCSR000EFM; antibody: Santa Cruz Biotech sc-150, Lot ID I1010); (E) HepG2 ChIP-seq (ENCODE ID ENCSR000EEE; antibody: Santa Cruz Biotech sc-150, Lot ID I1010); (F) HeLa-S3 ChIP-seq (ENCODE ID ENCSR000EDA; antibody: Santa Cruz Biotech sc-150, Lot ID I1010); (G) HepG2 forskolin ChIP-seq (ENCODE ID ENCSR000EEX; antibody: Santa Cruz Biotech sc-150, Lot ID I1010); (H) GM12878 ChIP-seq (ENCODE ID ENCSR000BRX; antibody: Santa Cruz Biotech sc-150, Lot ID I1010); (I) H1-hESC ChIP-seq (ENCODE ID ENCSR000EBV; antibody: Santa Cruz Biotech sc-150, Lot ID I1010); (J) HepG2 ChIP-seq (ENCODE ID ENCSR000BQI; antibody: Santa Cruz Biotech sc-150, Lot ID I1010).

**Supplementary Figure 11:**
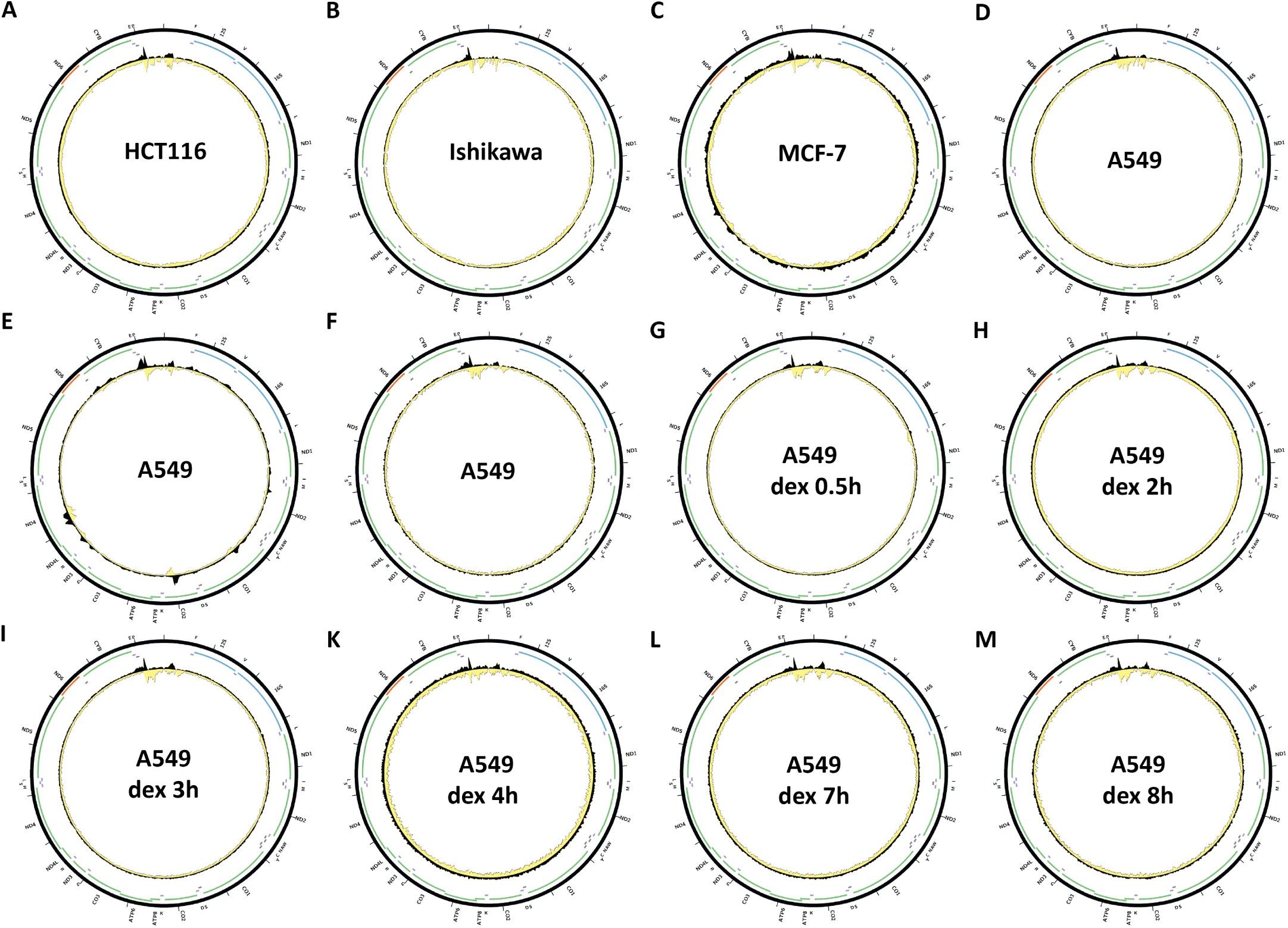
Evidence for mitochondrial genome occupancy by the CEBPB transcription factor. Black and yellow tracks show the forward- and reverse-strand ChIP-seq coverage over chrM. (A) HCT116ChIP-seq (ENCODE ID ENCSR000BSD; antibody: Santa Cruz Biotech sc-150, Lot ID I1010); (B) Ishikawa ChIP-seq (ENCODE ID ENCSR000BTT; antibody: Santa Cruz Biotech sc-150, Lot ID I1010); (C) MCF-7 ChIP-seq (ENCODE ID ENCSR000BSR; antibody: Santa Cruz Biotech sc-150, Lot ID I1010); (D) A549 ChIP-seq (ENCODE ID ENCSR000BUB; antibody: Santa Cruz Biotech sc-150, Lot ID I1010); (E) A549 ChIP-seq (ENCODE ID ENCSR000DYI; antibody: Santa Cruz Biotech sc-150, Lot ID I1010); (F) A549 ChIP-seq (ENCODE ID ENCSR701TCU; antibody: Santa Cruz Biotech sc-150, Lot ID I1010); (G) A549 dexamethasone 0.5hours ChIP-seq (ENCODE ID ENCSR447ZMS; antibody: Santa Cruz Biotech sc-150, Lot ID D2315); (H) A549 dexamethasone 2 hours ChIP-seq (ENCODE ID ENCSR182OZC; antibody: Santa Cruz Biotech sc-150, Lot ID D2315); (I) A549 dexamethasone 3 hours ChIP-seq (ENCODE ID ENCSR216GEB; antibody: Santa Cruz Biotech sc-150, Lot ID D2315); (K) A549 dexamethasone 4 hours ChIP-seq (ENCODE ID ENCSR606ZTC; antibody: Santa Cruz Biotech sc-150, Lot ID D2315); (L) A549 dexamethasone 7 hours ChIP-seq (ENCODE ID ENCFF887KKK; antibody: Santa Cruz Biotech sc-150, Lot ID D2315); (M) A549 dexamethasone 8 hours ChIP-seq (ENCODE ID ENCSR474DCX4; antibody: Santa Cruz Biotech sc-150, Lot ID D2315).

**Supplementary Figure 12:**
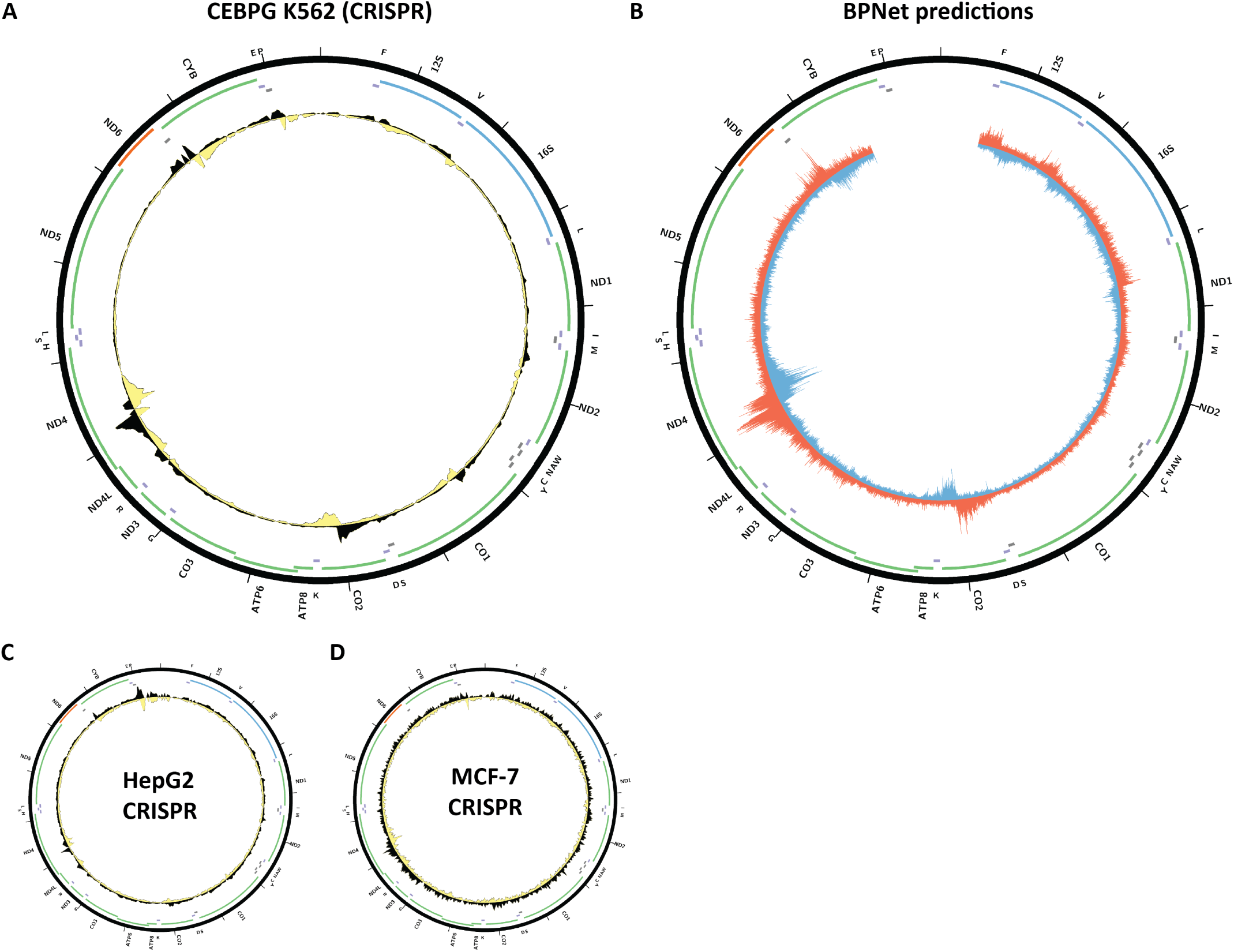
Evidence for mitochondrial genome occupancy by the CEBPG transcription factor. Black and yellow tracks show the forward- and reverse-strand ChIP-seq coverage over chrM. (A) K562 CETCH-seq (ENCODE ID ENCSR620VIC); (B) BPNet predictions over chrM (ENCODE ID ENCSR136AYM); (C) HepG2 ChIP-seq (ENCODE ID ENCSR639IIZ); (D) MCF-7 ChIP-seq (ENCODE ID ENCSR094ZCF).

**Supplementary Figure 13:**
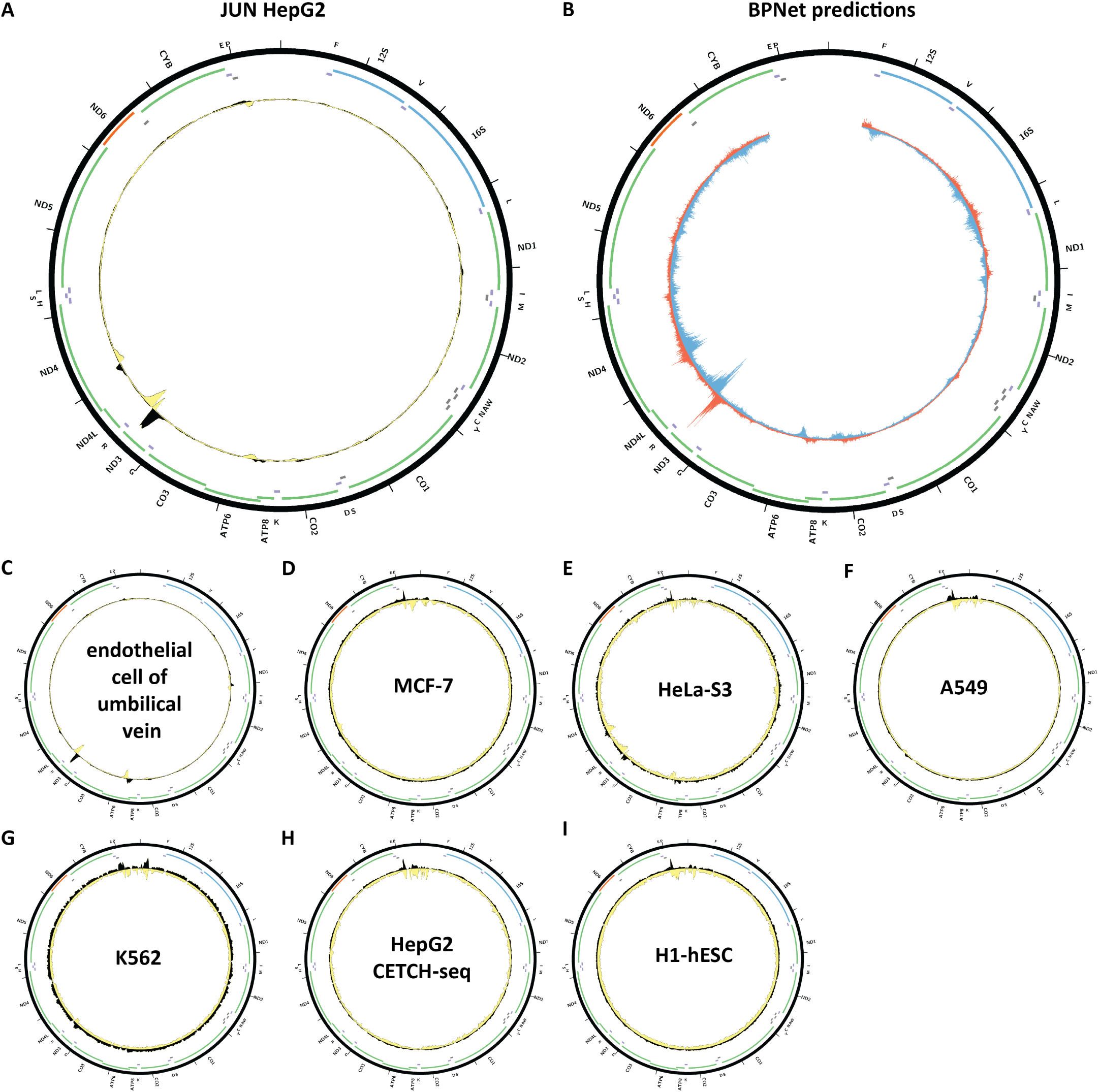
Evidence for mitochondrial genome occupancy by the JUN transcription factor. Black and yellow tracks show the forward- and reverse-strand ChIP-seq coverage over chrM. (A) HepG2ChIP-seq (ENCODE ID ENCSR000EEK; antibody: Santa Cruz Biotech sc-1694, Lot ID C2206); (B) BPNet predictions over chrM (ENCODE ID ENCSR701HFV); (C) endothelial cell of umbilical vein ChIP-seq (ENCODE ID ENCSR000EFA; antibody: Santa Cruz Biotech sc-1694, Lot ID C2206); (D) MCF-7 ChIP-seq (ENCODE ID ENCSR176EXN; antibody: Santa Cruz Biotech sc-1694, Lot ID C2206); (E) HeLa-S3 ChIP-seq (ENCODE ID ENCSR000EDG; antibody: Santa Cruz Biotech sc-1694, Lot ID C2206); (F) A549 ChIP-seq (ENCODE ID ENCSR996DUT; antibody: Santa Cruz Biotech sc-1694, Lot ID C2206); (G) K562 ChIP-seq (ENCODE ID ENCSR000EFS; antibody: Santa Cruz Biotech sc-1694, Lot ID C2206); (H) HepG2 CETCH-seq (ENCODE ID ENCSR747VUU); (I) H1-hESC ChIP-seq (ENCODE ID ENCSR000ECA; antibody: Santa Cruz Biotech sc-1694, Lot ID C2206).

**Supplementary Figure 14:**
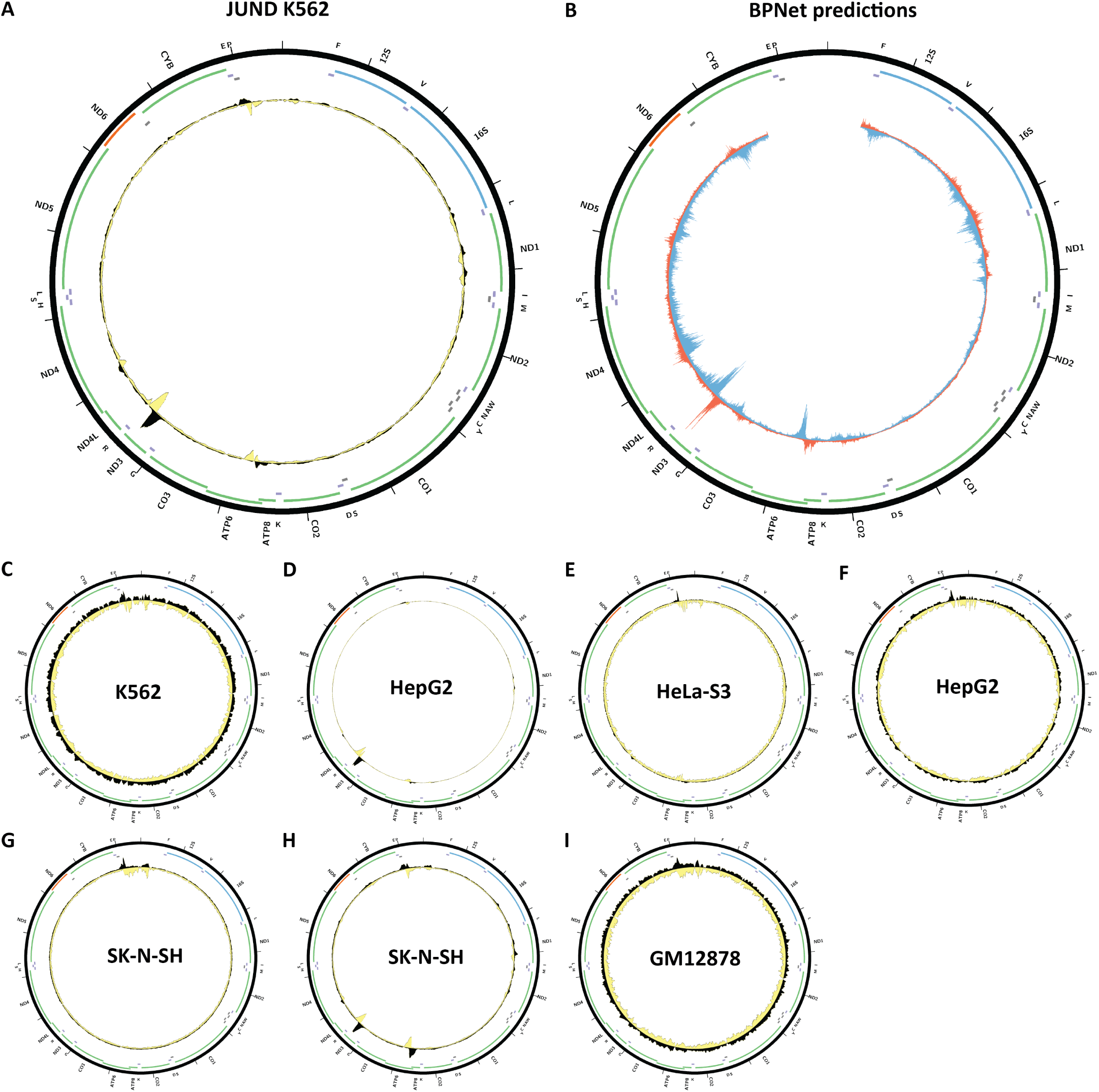
Evidence for mitochondrial genome occupancy by the JUND transcription factor. Black and yellow tracks show the forward- and reverse-strand ChIP-seq coverage over chrM. (A) K562ChIP-seq (ENCODE ID ENCSR000DJX; GFP-tagged); (B) BPNet predictions over chrM (ENCODE ID ENCSR767UUB); (C) K562 ChIP-seq (ENCODE ID ENCSR000DJX; antibody: GFP-tagged); (D) HepG2 ChIP-seq (ENCODE ID ENCSR000EEI; antibody: Santa Cruz Biotech sc-74); (E) HeLa-S3 ChIP-seq (ENCODE ID ENCSR000EDH; antibody: Santa Cruz Biotech sc-74); (F) HepG2 ChIP-seq (ENCODE ID ENCSR000BGK; antibody: Santa Cruz Biotech sc-74); (G) SK-N-SH ChIP-seq (ENCODE ID ENCSR000BSK; antibody: Santa Cruz Biotech sc-74); (H) SK-N-SH ChIP-seq (ENCODE ID ENCSR000EIB; antibody: Santa Cruz Biotech sc-74); (I) GM18278 ChIP-seq (ENCODE ID ENCSR000DYS; antibody: Santa Cruz Biotech sc-74).

**Supplementary Figure 15:**
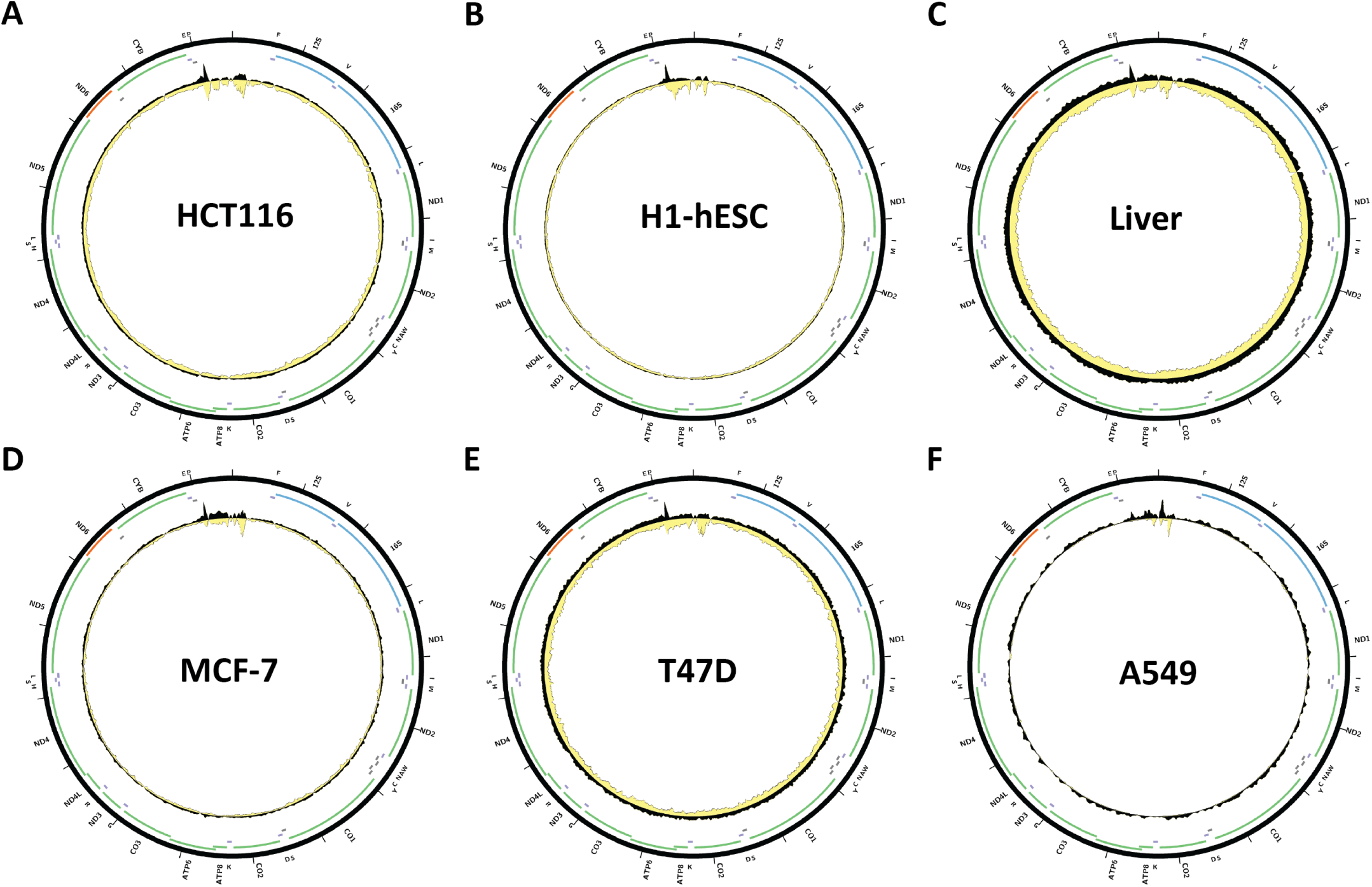
Evidence for mitochondrial genome occupancy by the JUND transcription factor. Black and yellow tracks show the forward- and reverse-strand ChIP-seq coverage over chrM. (A) HCT116ChIP-seq (ENCODE ID ENCSR000BSA; antibody: Santa Cruz Biotech sc-74); (B) H1-hESC ChIP-seq (ENCODE ID ENCSR000BKP; antibody: Santa Cruz Biotech sc-74); (C) Liver ChIP-seq (ENCODE ID ENCSR837GTK; antibody: Santa Cruz Biotech sc-74); (D) MCF-7 ChIP-seq (ENCODE ID ENCSR000BSU; antibody: Santa Cruz Biotech sc-74); (E) T47D ChIP-seq (ENCODE ID ENCSR000BVO; antibody: Santa Cruz Biotech sc-74); (F) A549 ChIP-seq (ENCODE ID ENCSR000BRF; antibody: Santa Cruz Biotech sc-74).

**Supplementary Figure 16:**
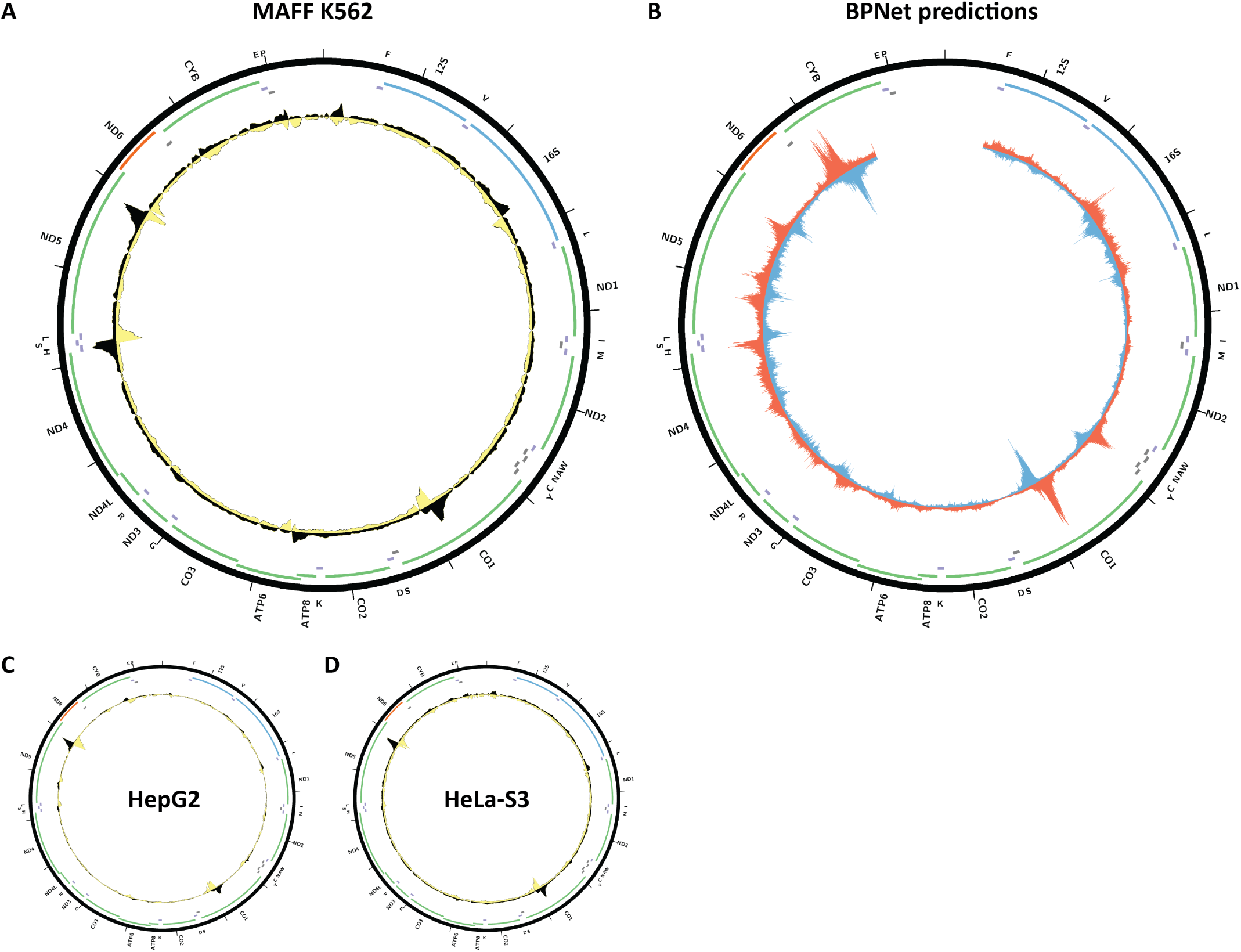
Evidence for mitochondrial genome occupancy by the MAFF transcription factor. Black and yellow tracks show the forward- and reverse-strand ChIP-seq coverage over chrM. (A) K562ChIP-seq (ENCODE ID ENCSR000EGI; antibody: Sigma M8194, Lot ID 125K4837); (B) BPNet predictions over chrM (ENCODE ID ENCSR512XRO); (C) HepG2 ChIP-seq (ENCODE ID ENCSR000EEC; antibody: Sigma M8194, Lot ID 125K4837); (D) HeLa-S3 ChIP-seq (ENCODE ID ENCSR140DSL; antibody: Sigma M8194, Lot ID 125K4837).

**Supplementary Figure 17:**
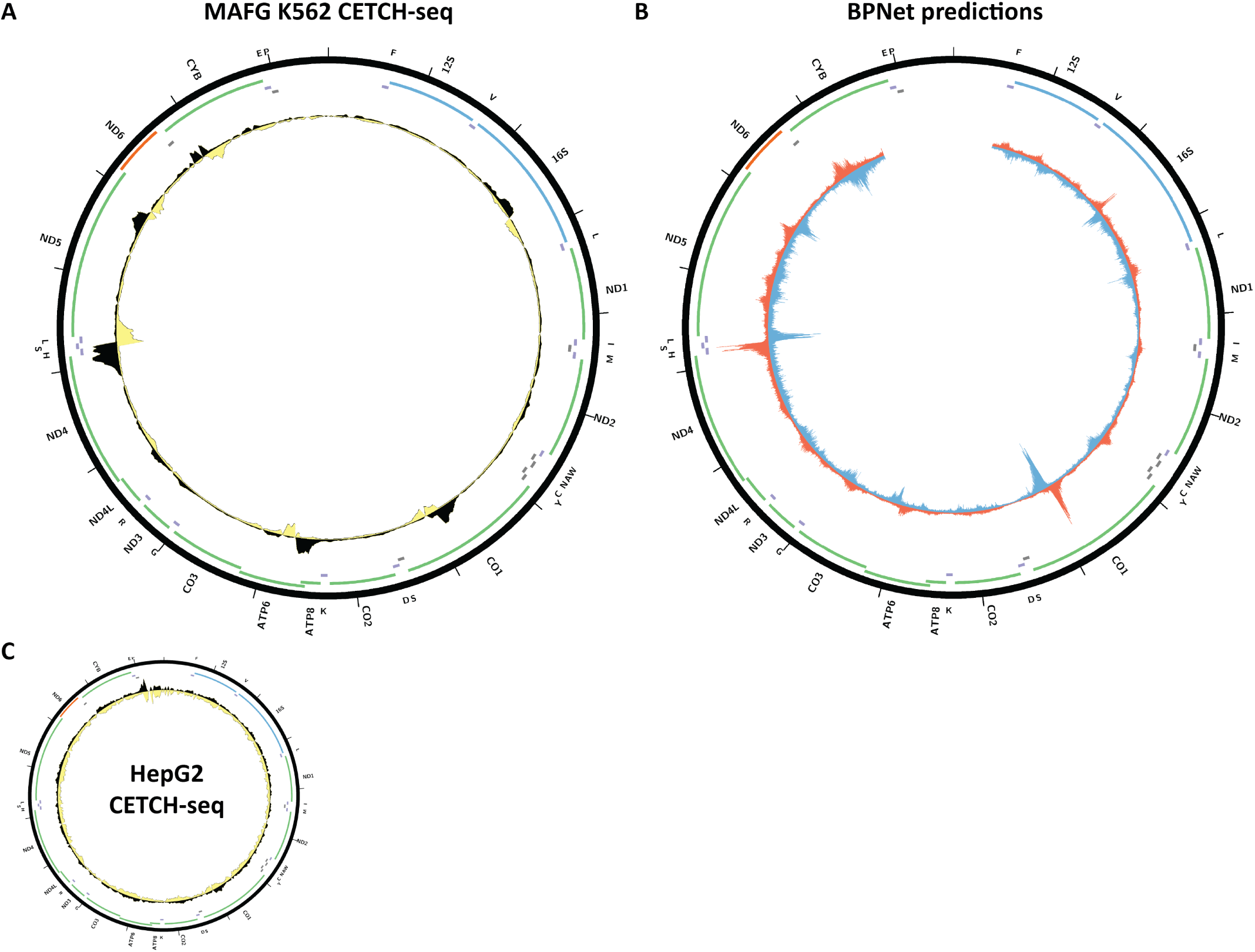
Evidence for mitochondrial genome occupancy by the MAFG transcription factor. Black and yellow tracks show the forward- and reverse-strand ChIP-seq coverage over chrM. (A) K562 CETCH- seq (ENCODE ID ENCSR818DQV); (B) BPNet predictions over chrM (ENCODE ID ENCSR324HYA); (C) HepG2 CETCH-seq (ENCODE ID ENCSR708KAA).

**Supplementary Figure 18:**
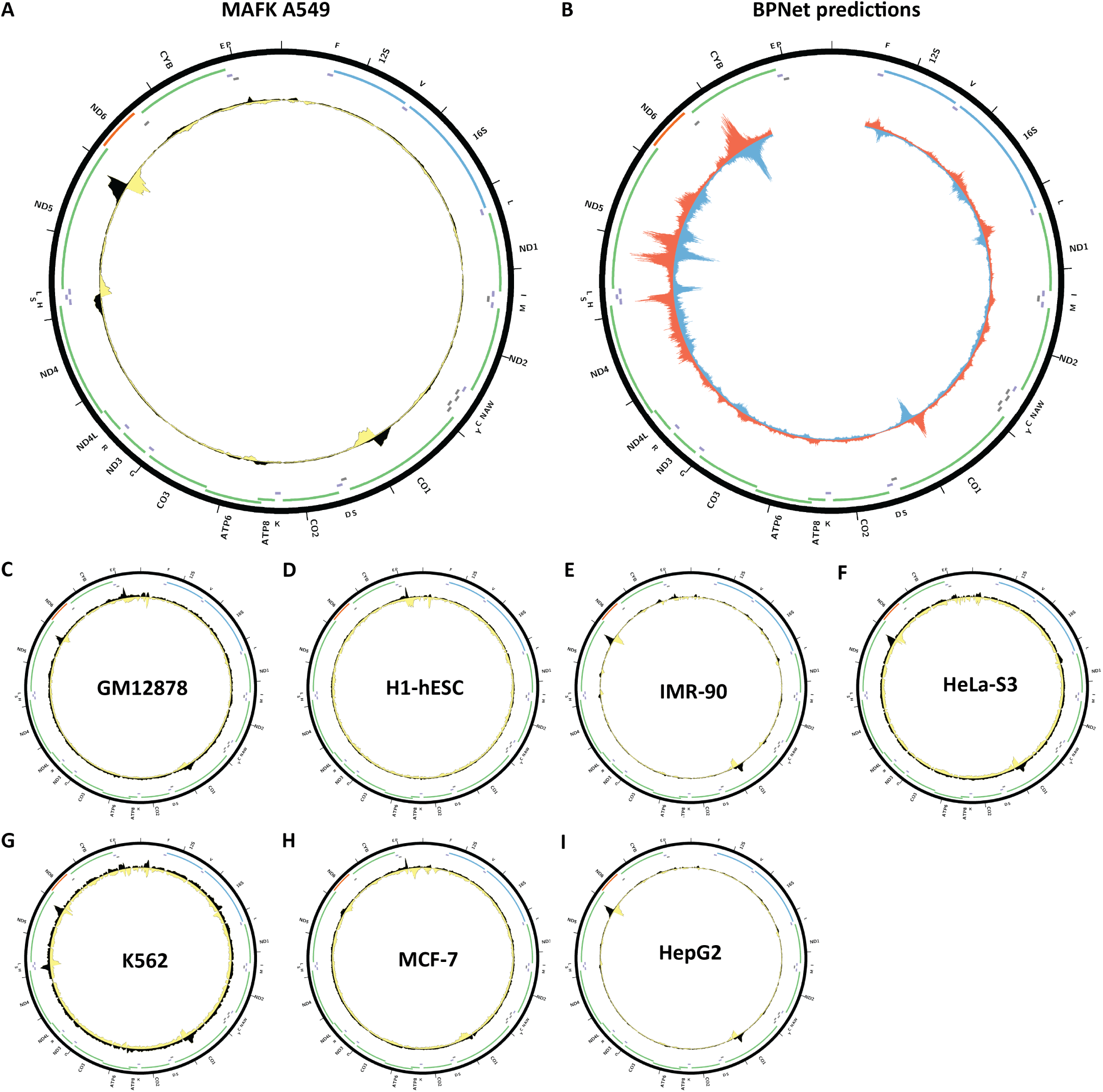
Evidence for mitochondrial genome occupancy by the MAFK transcription factor. Black and yellow tracks show the forward- and reverse-strand ChIP-seq coverage over chrM. (A) A549ChIP-seq (ENCODE ID ENCSR541WQI; antibody: Abcam ab50322, Lot ID 904274); (B) BPNet predictions over chrM (ENCODE ID ENCSR415PWX); (C) GM18278 ChIP-seq (ENCODE ID ENCSR000DYV; antibody: Abcam ab50322, Lot ID 904274); (D) H1-hESC ChIP-seq (ENCODE ID ENCSR000EBS; antibody: Abcam ab50322, Lot ID 904274); (E) IMR-90 ChIP-seq (ENCODE ID ENCSR000EFH; antibody: Abcam ab50322, Lot ID 904274); (F) HeLa-S3 ChIP-seq (ENCODE ID ENCSR000ECK; antibody: Abcam ab50322, Lot ID 904274); (G) K562 ChIP-seq (ENCODE ID ENCSR000EGX; antibody: Abcam ab50322, Lot ID 904274); (H) MCF-7 ChIP-seq (ENCODE ID ENCSR555PBN; antibody: Abcam ab50322, Lot ID 904274); (I) HepG2 ChIP-seq (ENCODE ID ENCSR000EDZ; antibody: Santa Cruz Biotech sc-477, Lot ID K1709).

**Supplementary Figure 19:**
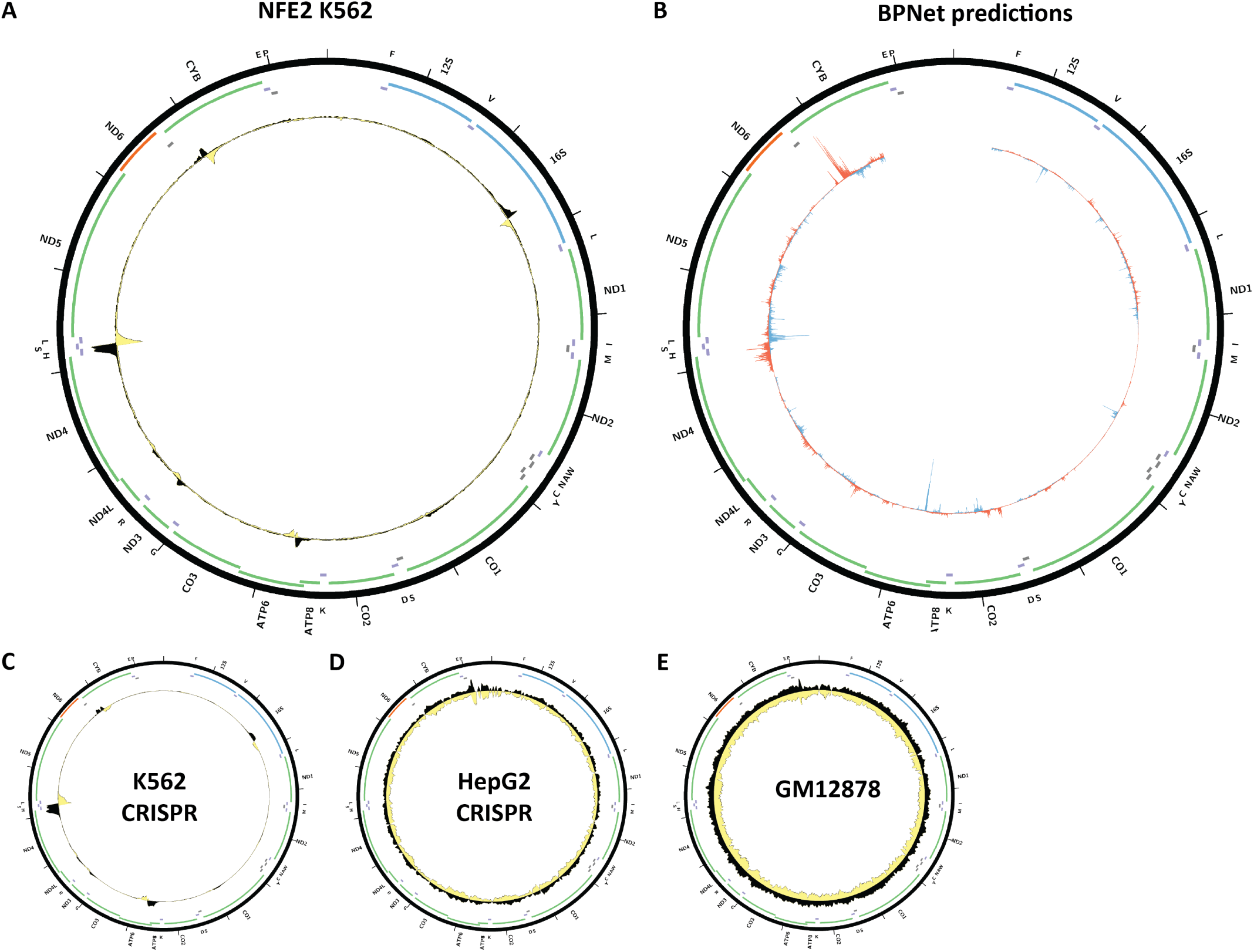
Evidence for mitochondrial genome occupancy by the NFE2 transcription factor. Black and yellow tracks show the forward- and reverse-strand ChIP-seq coverage over chrM. (A) K562 ChIP-seq (ENCODE ID ENCSR000FAF; antibody Santa Cruz Biotech sc-291, Lot ID D1703); (B) BPNet predictions over chrM (ENCODE ID ENCSR833AMA); (C) K562 CETCH-seq (ENCODE ID ENCSR552YGL); (C) HepG2 CETCH-seq (ENCODE ID ENCSR983FBD); (C) GM12878 ChIP-seq (ENCODE ID ENCSR000DZY; antibody Santa Cruz Biotech sc-22827).

**Supplementary Figure 20:**
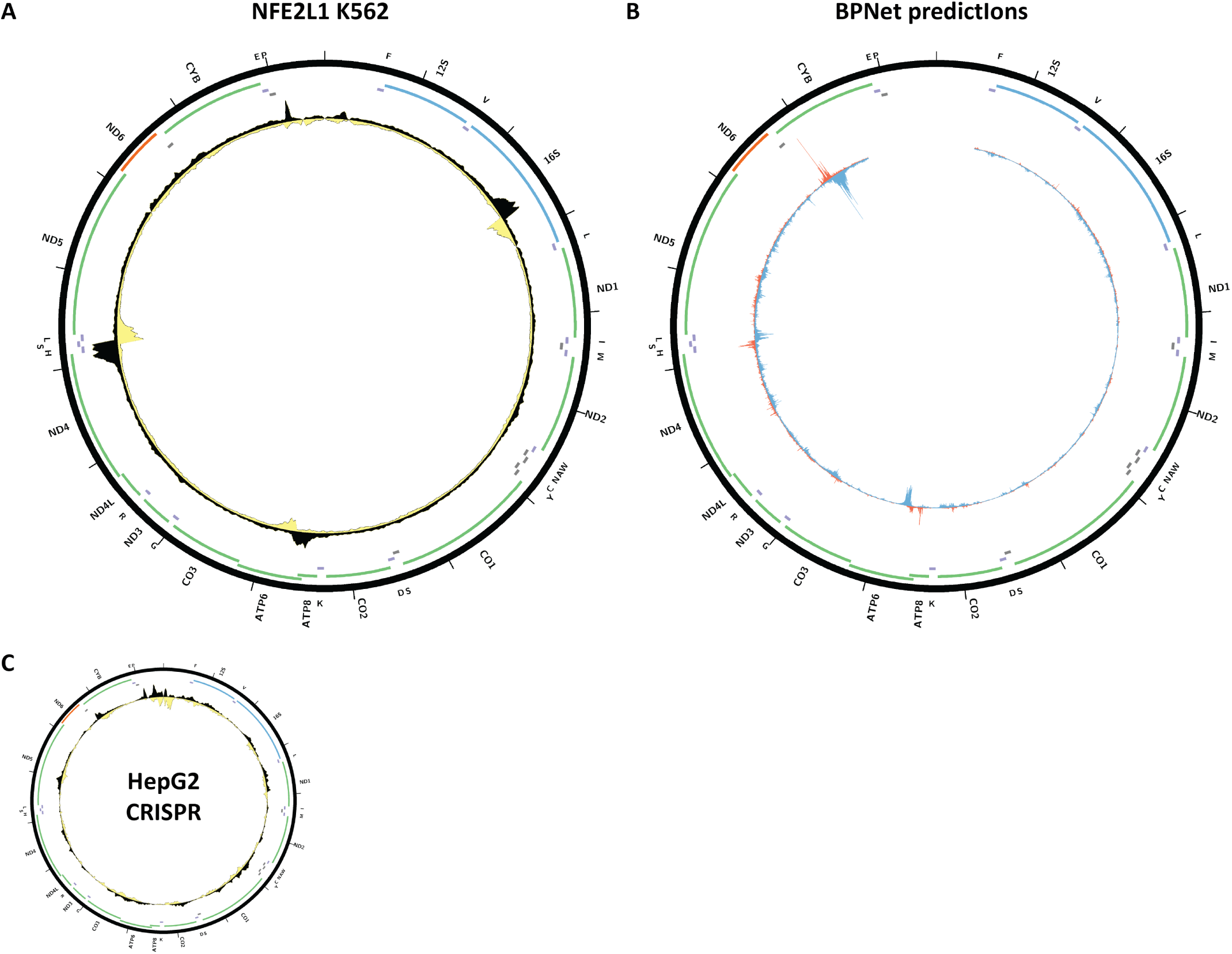
Evidence for mitochondrial genome occupancy by the NFE2L1transcription factor. Black and yellow tracks show the forward- and reverse-strand ChIP-seq coverage over chrM. (A) K562 ChIP-seq (ENCODE ID ENCSR632SHZ; GFP-tagged); (B) BPNet predictions over chrM; (C) HepG2 CETCH-seq (ENCODE ID ENCSR543SBE; antibody:).

**Supplementary Figure 21:**
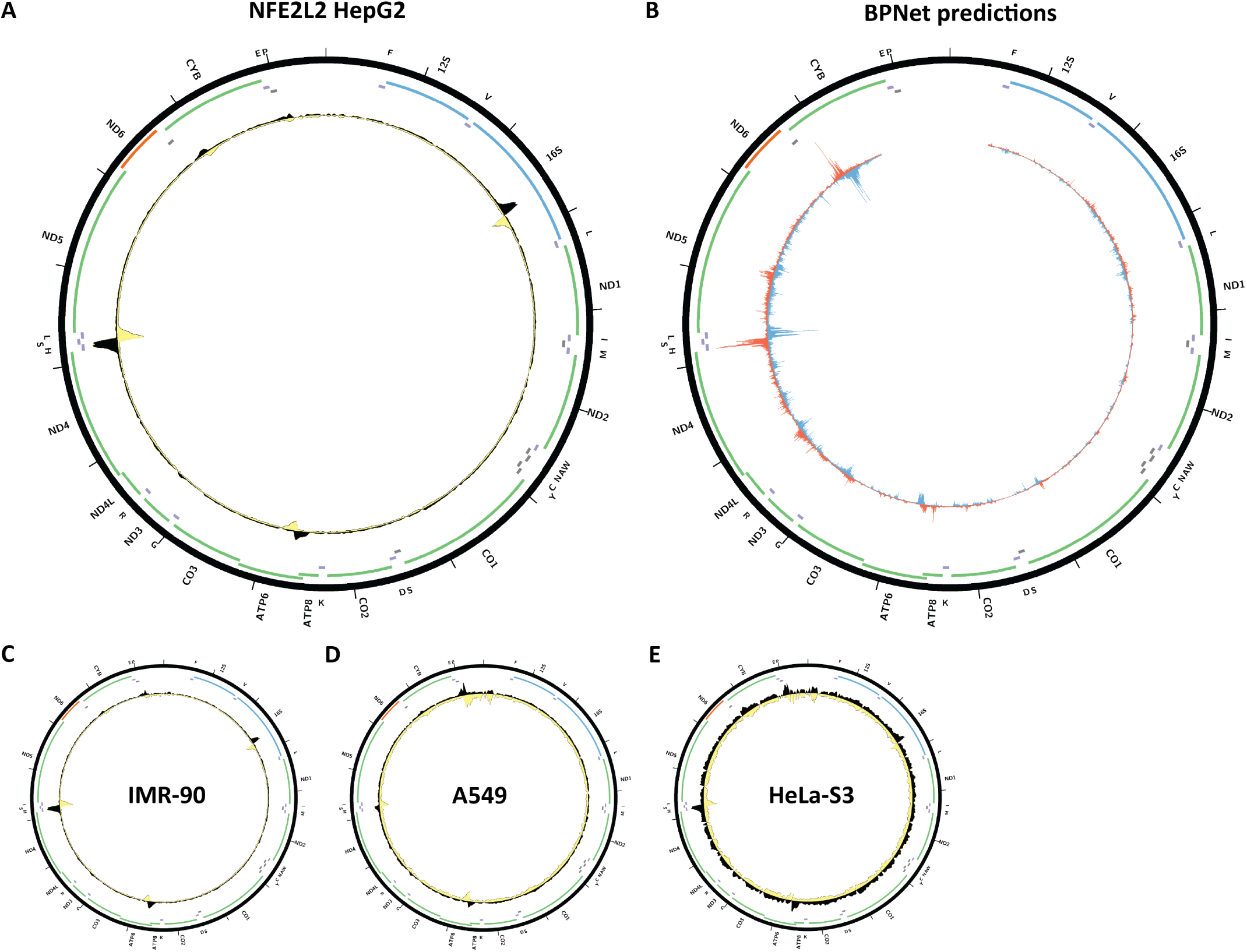
Evidence for mitochondrial genome occupancy by the NFE2L2transcription factor. Black and yellow tracks show the forward- and reverse-strand ChIP-seq coverage over chrM. (A) HepG2 ChIP-seq (ENCODE ID ENCSR488EES; antibody: Santa Cruz Biotech sc-13032, Lot ID A1711); (B) BPNet predictions over chrM (ENCODE ID ENCSR301SDD); (C) IMR-90 ChIP-seq (ENCODE ID ENCSR197WGI; antibody: Santa Cruz Biotech sc-13032, Lot ID A1711); (D) A549 ChIP-seq (ENCODE ID ENCSR584GHV; antibody: Santa Cruz Biotech sc-13032, Lot ID A1711); (E) HeLa-S3 ChIP-seq (ENCODE ID ENCSR707IUN; antibody: Santa Cruz Biotech sc-13032, Lot ID A1711).

**Supplementary Figure 22:**
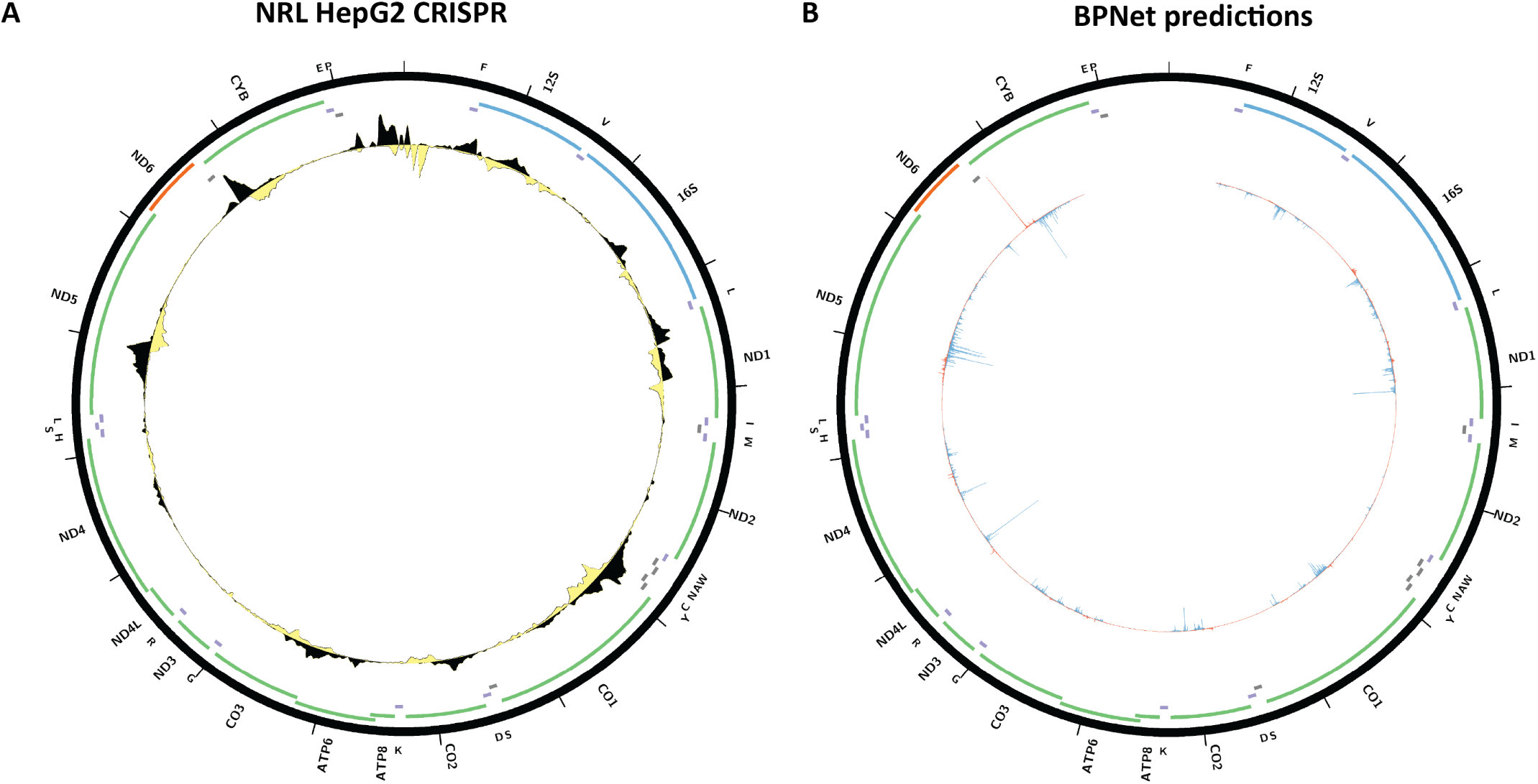
Evidence for mitochondrial genome occupancy by the NRL transcription factor. Black and yellow tracks show the forward- and reverse-strand ChIP-seq coverage over chrM. (A) HepG2 CETCH-seq (ENCODE ID ENCSR518KLO); (B) BPNet predictions over chrM.

**Supplementary Figure 23:**
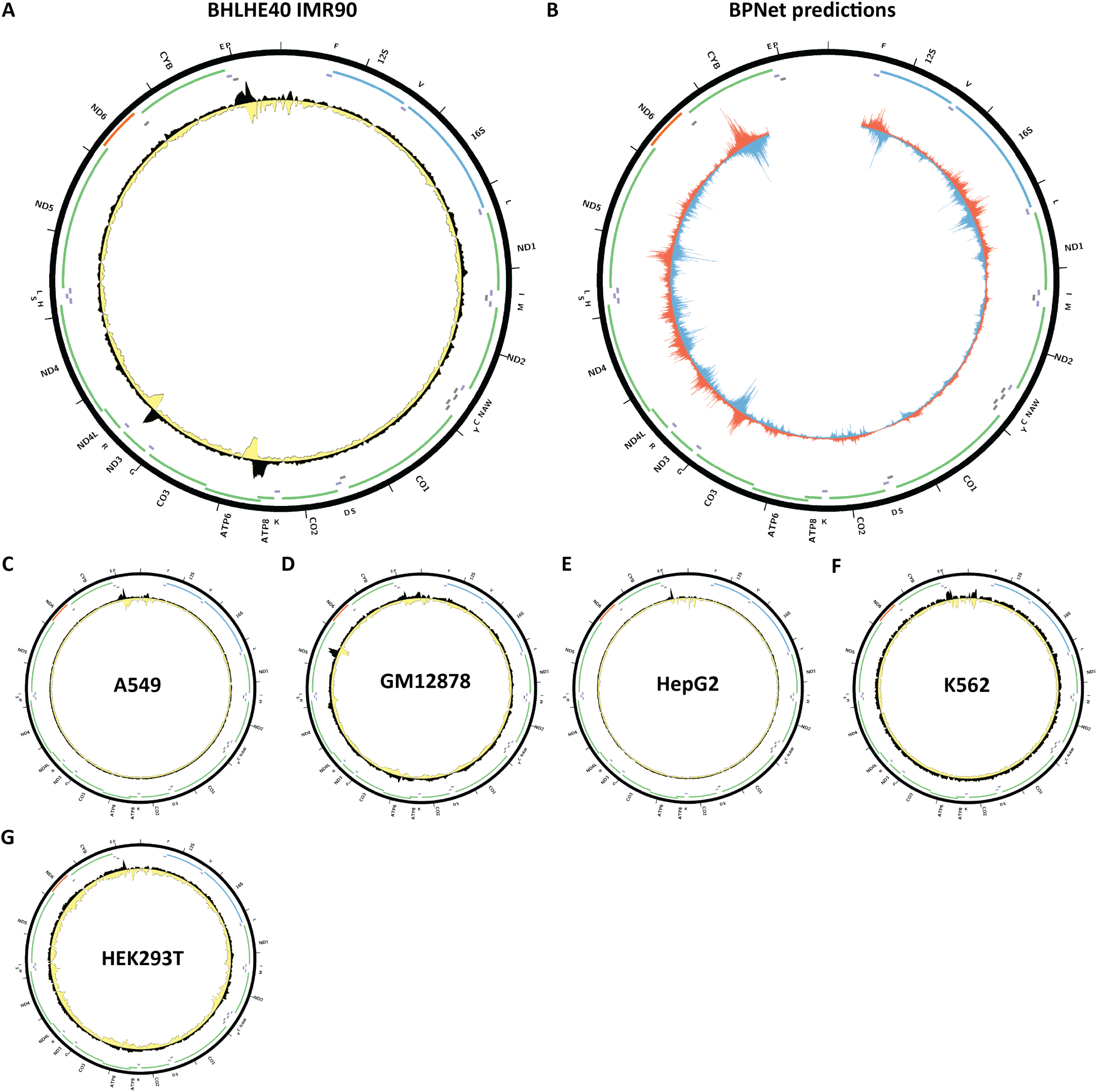
Evidence for mitochondrial genome occupancy by the BHLHE40 transcription factor. Black and yellow tracks show the forward- and reverse-strand ChIP-seq coverage over chrM. (A) IMR90 ChIP-seq (ENCODE ID ENCSR957KYB; antibody: Novus NB100-1800, Lot ID A1); (B) BPNet predictions over chrM (ENCODE ID ENCSR695LTU); (C) A549 ChIP-seq (ENCODE ID ENCSR000DYJ; antibody: Santa Cruz Biotech sc-101023); (D) GM12878 ChIP-seq (ENCODE ID ENCSR987MTA; antibody: Novus NB100-1800, Lot ID A1); (E) HepG2 ChIP-seq (ENCODE ID ENCSR000BID; antibody: Santa Cruz Biotech sc-101023); (F) K562 ChIP-seq (ENCODE ID ENCSR000EGV; antibody: Novus NB100-1800, Lot ID A1); (G) HEK293T ChIP-seq (ENCODE ID ENCSR789GVU; antibody: Novus NB100-1800, Lot ID A1).

**Supplementary Figure 24:**
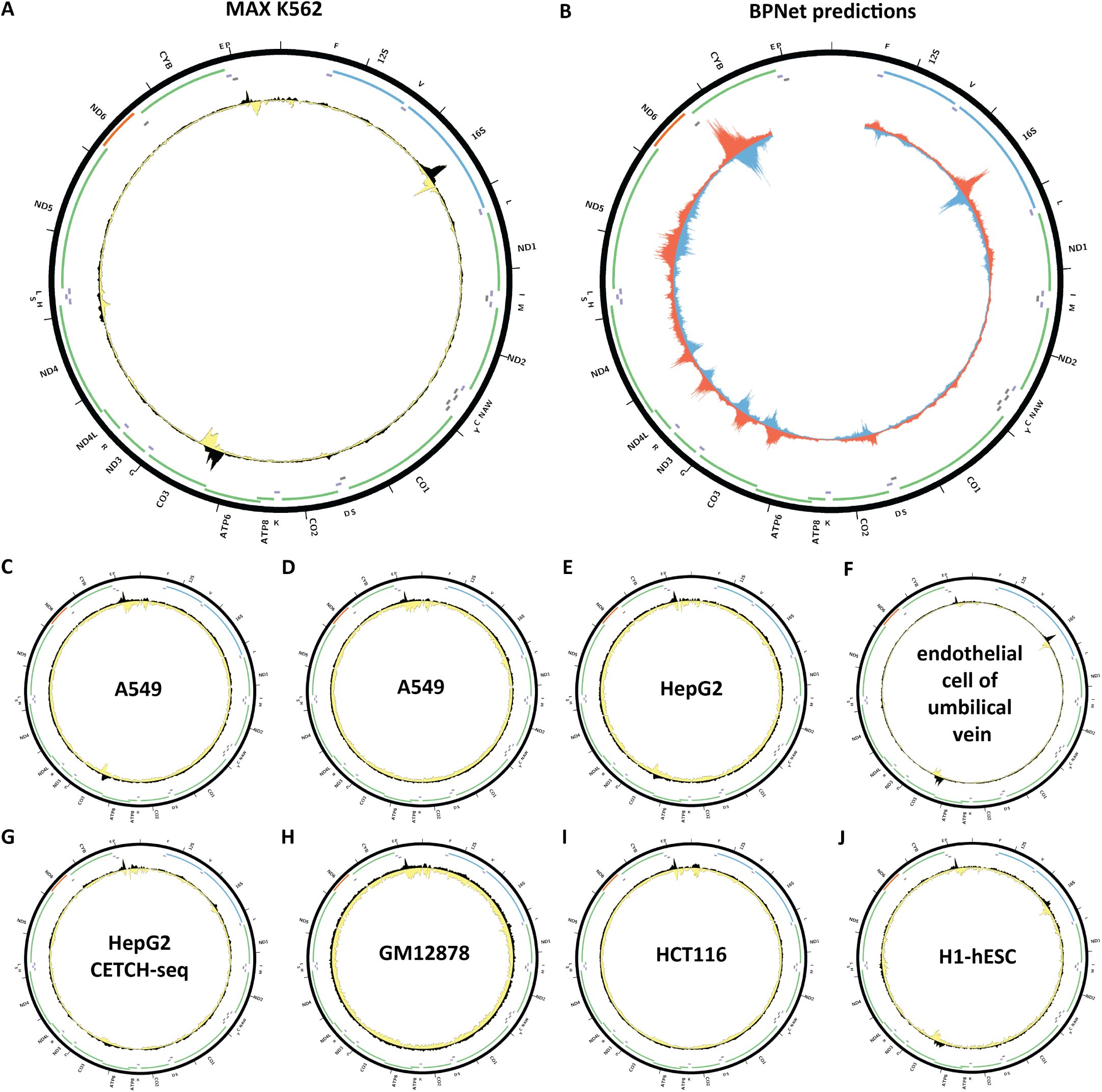
Evidence for mitochondrial genome occupancy by the MAX transcription factor. Black and yellow tracks show the forward- and reverse-strand ChIP-seq coverage over chrM. (A) K562ChIP-seq (ENCODE ID ENCSR000FAE; antibody: Santa Cruz Biotech sc-197, Lot ID J0809); (B) BPNet predictions over chrM (ENCODE ID ENCSR445ZHD); (C) A549 ChIP-seq (ENCODE ID ENCSR000DYG; antibody: Santa Cruz Biotech sc-197, Lot ID J0809); (D) A549 ChIP-seq (ENCODE ID ENCSR000BTJ; antibody: Santa Cruz Biotech sc-197, Lot ID J0809); (E) HepG2 ChIP-seq (ENCODE ID ENCSR000EDS; antibody: Santa Cruz Biotech sc-197, Lot ID J0809); (F) endothelial cell of umbilical vein ChIP-seq (ENCODE ID ENCSR000EEZ; antibody: Santa Cruz Biotech sc-197, Lot ID J0809); (G) HepG2 CETCH-seq (ENCODE ID ENCSR168DYA); (H) GM18278 ChIP-seq (ENCODE ID ENCSR000DZF; antibody: Santa Cruz Biotech sc-197, Lot ID J0809); (I) HCT116 ChIP-seq (ENCODE ID ENCSR000BSH; antibody: Santa Cruz Biotech sc-197, Lot ID J0809); (J) H1-hESC ChIP-seq (ENCODE ID ENCSR000EUP; antibody: Santa Cruz Biotech sc-197, Lot ID J0809).

**Supplementary Figure 25:**
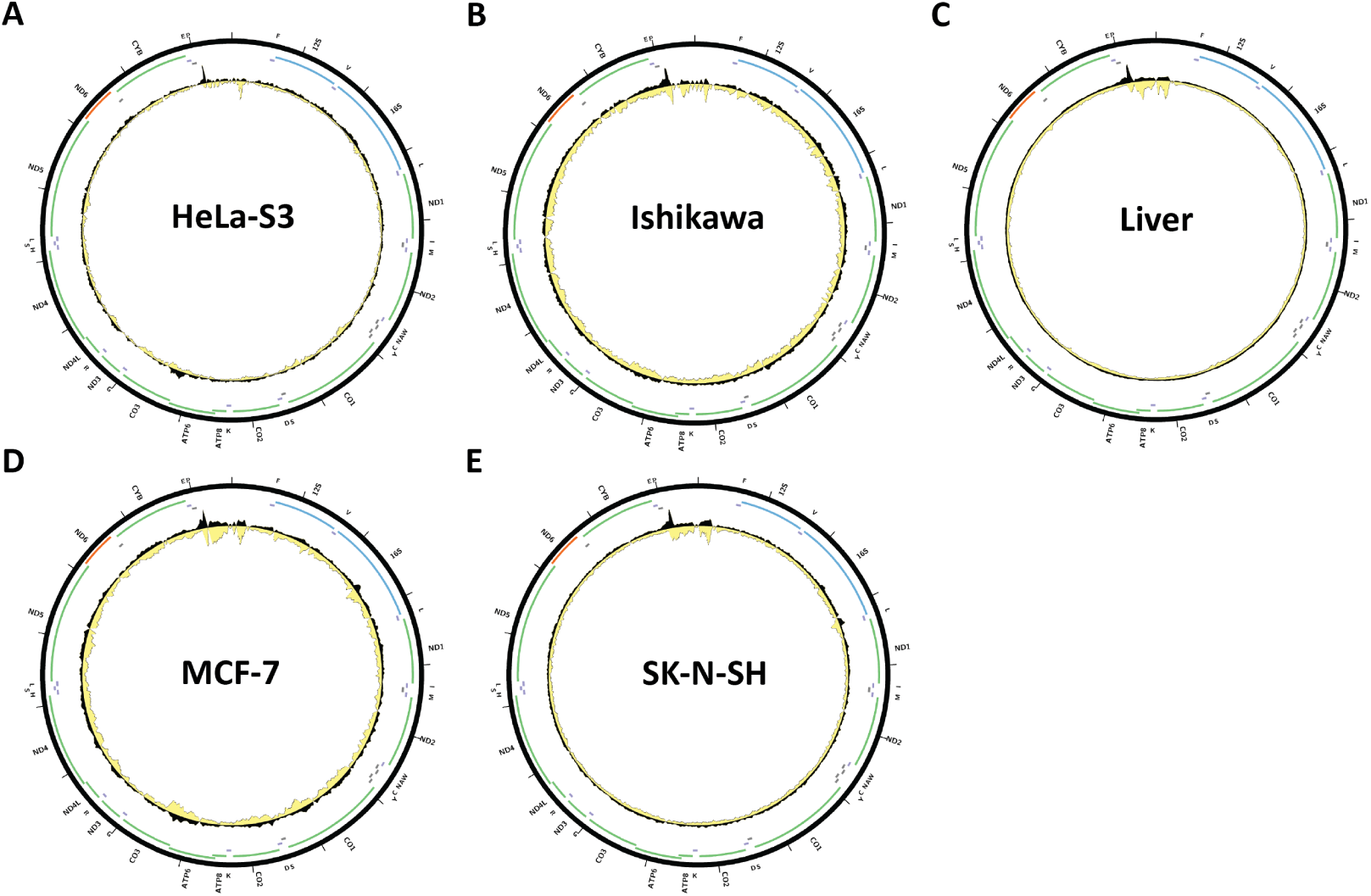
Evidence for mitochondrial genome occupancy by the MAX transcription factor. Black and yellow tracks show the forward- and reverse-strand ChIP-seq coverage over chrM. (A) HeLa-S3ChIP-seq (ENCODE ID ENCSR000EZF; antibody: Santa Cruz Biotech sc-197, Lot ID J0809); (B) Ishikawa ChIP-seq (ENCODE ID ENCSR000BTY; antibody: Santa Cruz Biotech sc-197, Lot ID J0809); (C) Liver ChIP-seq (ENCODE ID ENCSR521IID; antibody: Santa Cruz Biotech sc-197, Lot ID J0809); (D) MCF-7 ChIP-seq (ENCODE ID ENCSR000BUL; antibody: Santa Cruz Biotech sc-197, Lot ID J0809); (E) SK-N-SH ChIP-seq (ENCODE ID ENCSR000BVD; antibody: Santa Cruz Biotech sc-197, Lot ID J0809).

**Supplementary Figure 26:**
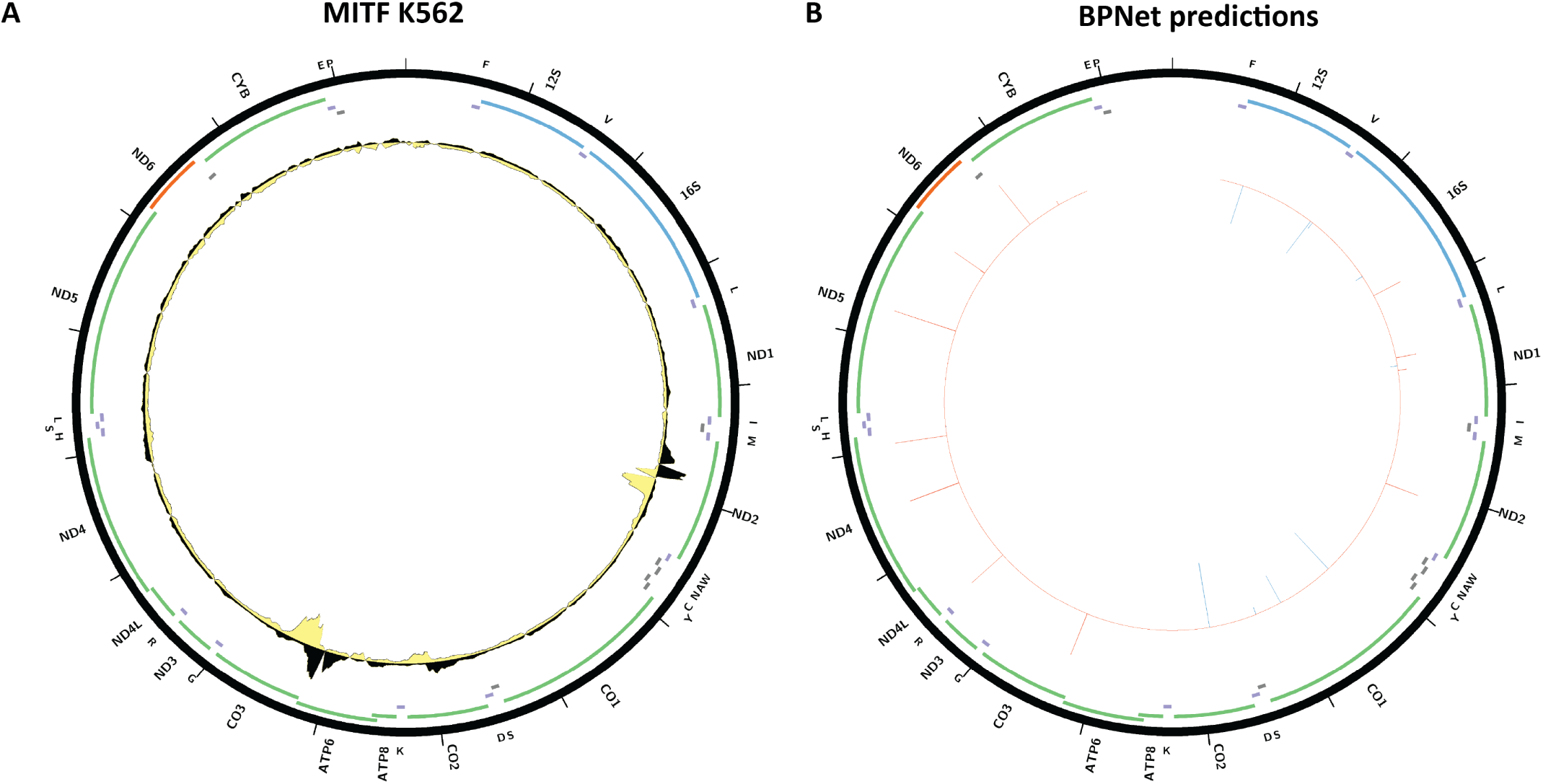
Evidence for mitochondrial genome occupancy by the MITF transcription factor. Black and yellow tracks show the forward- and reverse-strand ChIP-seq coverage over chrM. (A) K562ChIP-seq (ENCODE ID ENCSR797SWM; antibody: Active Motif 39789, Lot ID 11313002); (B) BPNet predictions over chrM (ENCODE ID ENCSR551LHV).

**Supplementary Figure 27:**
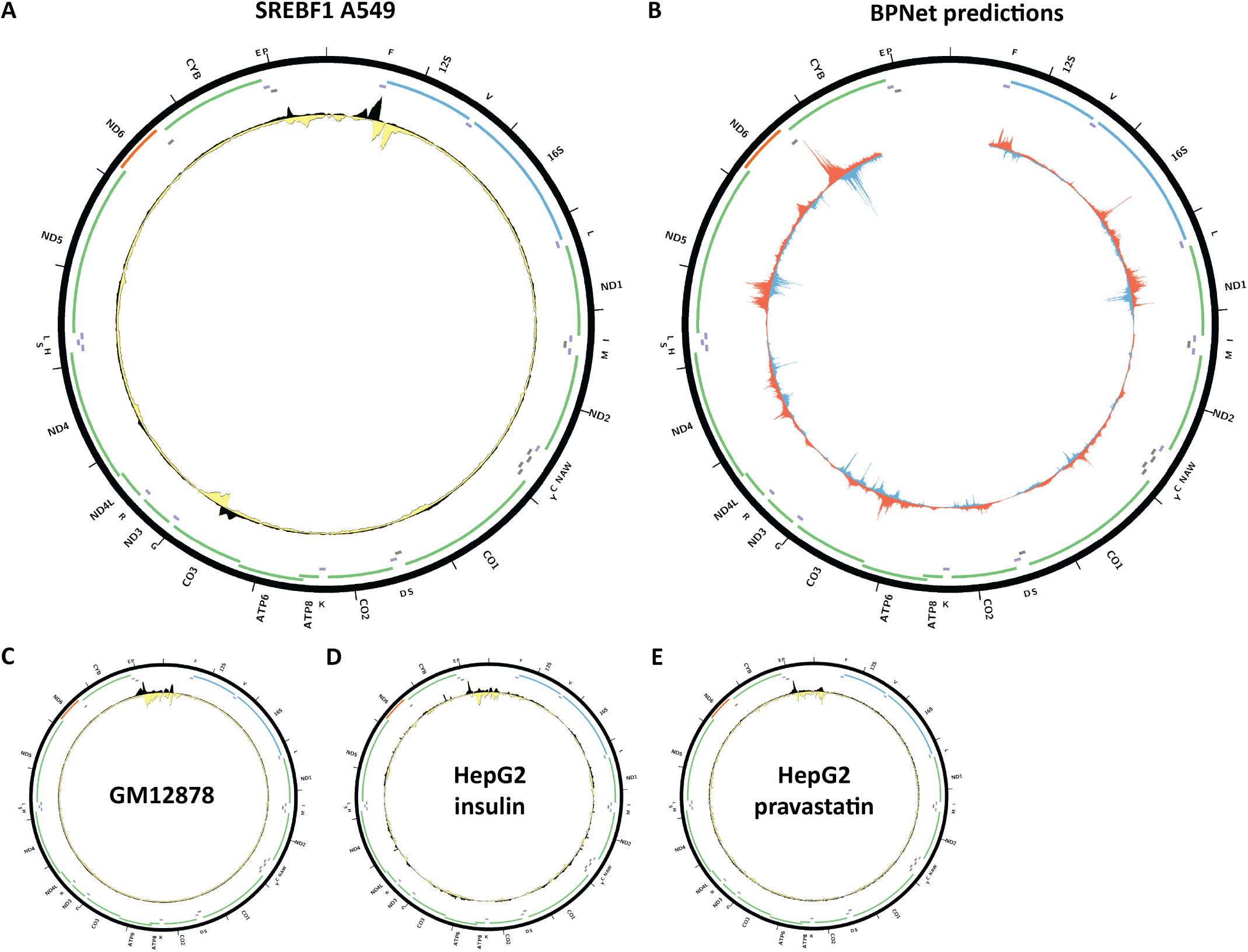
Evidence for mitochondrial genome occupancy by the SREBF1transcription factor. Black and yellow tracks show the forward- and reverse-strand ChIP-seq coverage over chrM. (A) A549 ChIP-seq (ENCODE ID ENCSR897MYK; antibody: Santa Cruz Biotech sc-8984, Lot ID 10211); (B) BPNet predictions over chrM (ENCODE ID ENCSR182NCE); (C) GM12878 ChIP-seq (ENCODE ID ENCSR000DYU; antibody: Santa Cruz Biotech sc-8984, Lot ID 10211); (D) HepG2 (10 *µ*M insulin, 100 *µ*M 22-hydroxycholesterol, 6 hours post-treatment) ChIP-seq (ENCODE ID ENCSR000EEO; antibody: Santa Cruz Biotech sc-8984, Lot ID 10211); (E) HepG2 (2 *µ*M pravastatin, 16 hours post-treatment) ChIP-seq (ENCODE ID ENCSR000EZP; antibody: Santa Cruz Biotech sc-8984, Lot ID 10211).

**Supplementary Figure 28:**
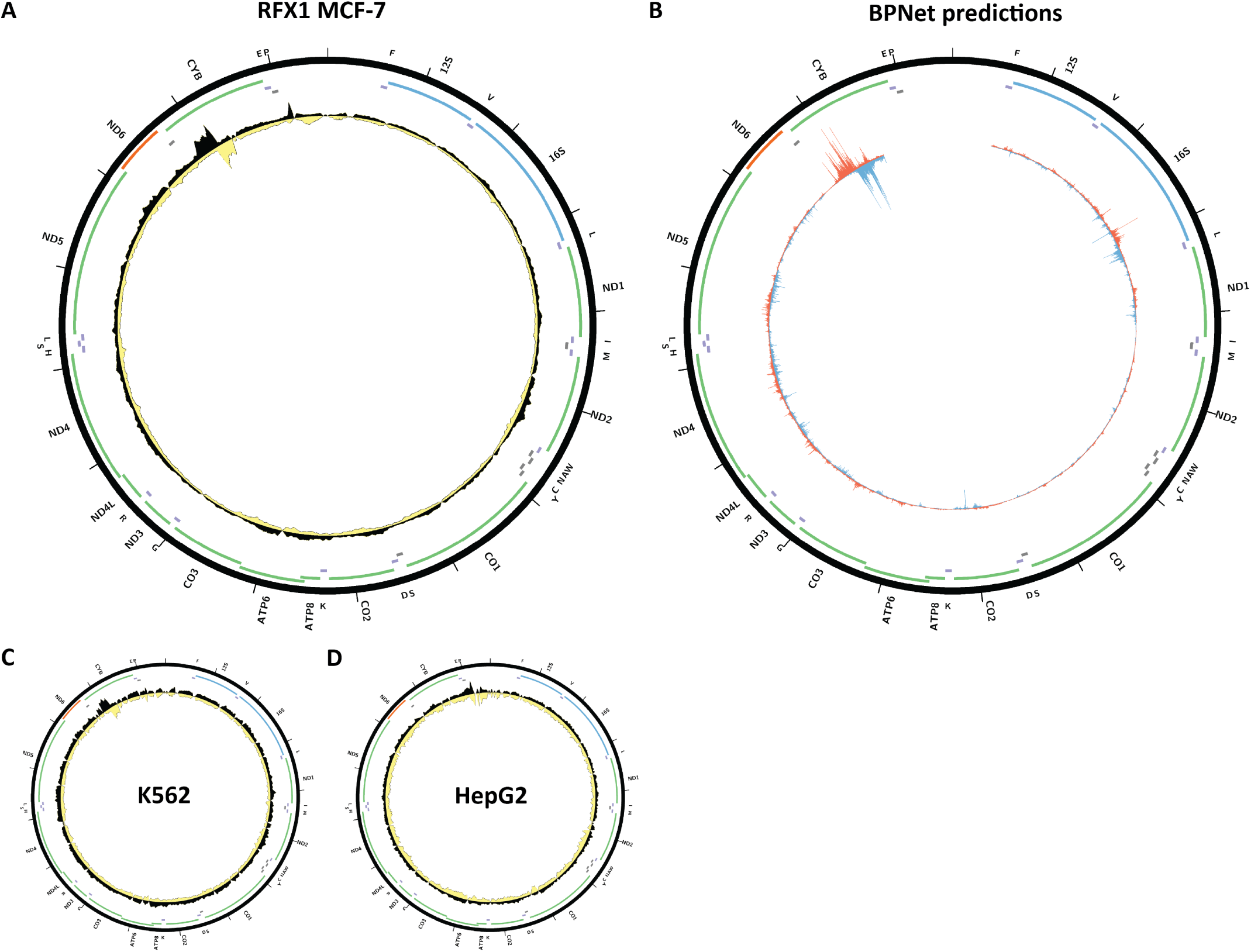
Evidence for mitochondrial genome occupancy by the RFX1transcription factor. Black and yellow tracks show the forward- and reverse-strand ChIP-seq coverage over chrM. (A) MCF-7 ChIP-seq (ENCODE ID ENCSR788XNX; antibody: Santa Cruz Biotech sc-10652, Lot ID E1911); (B) BPNet predictions over chrM (ENCODE ID ENCSR594OAT); (C) K562 ChIP-seq (ENCODE ID ENCSR968GIB; antibody: Santa Cruz Biotech sc-10652, Lot ID E1911); (D) HepG2 ChIP-seq (ENCODE ID ENCSR928API; antibody: Santa Cruz Biotech sc-10652, Lot ID E1911).

**Supplementary Figure 29:**
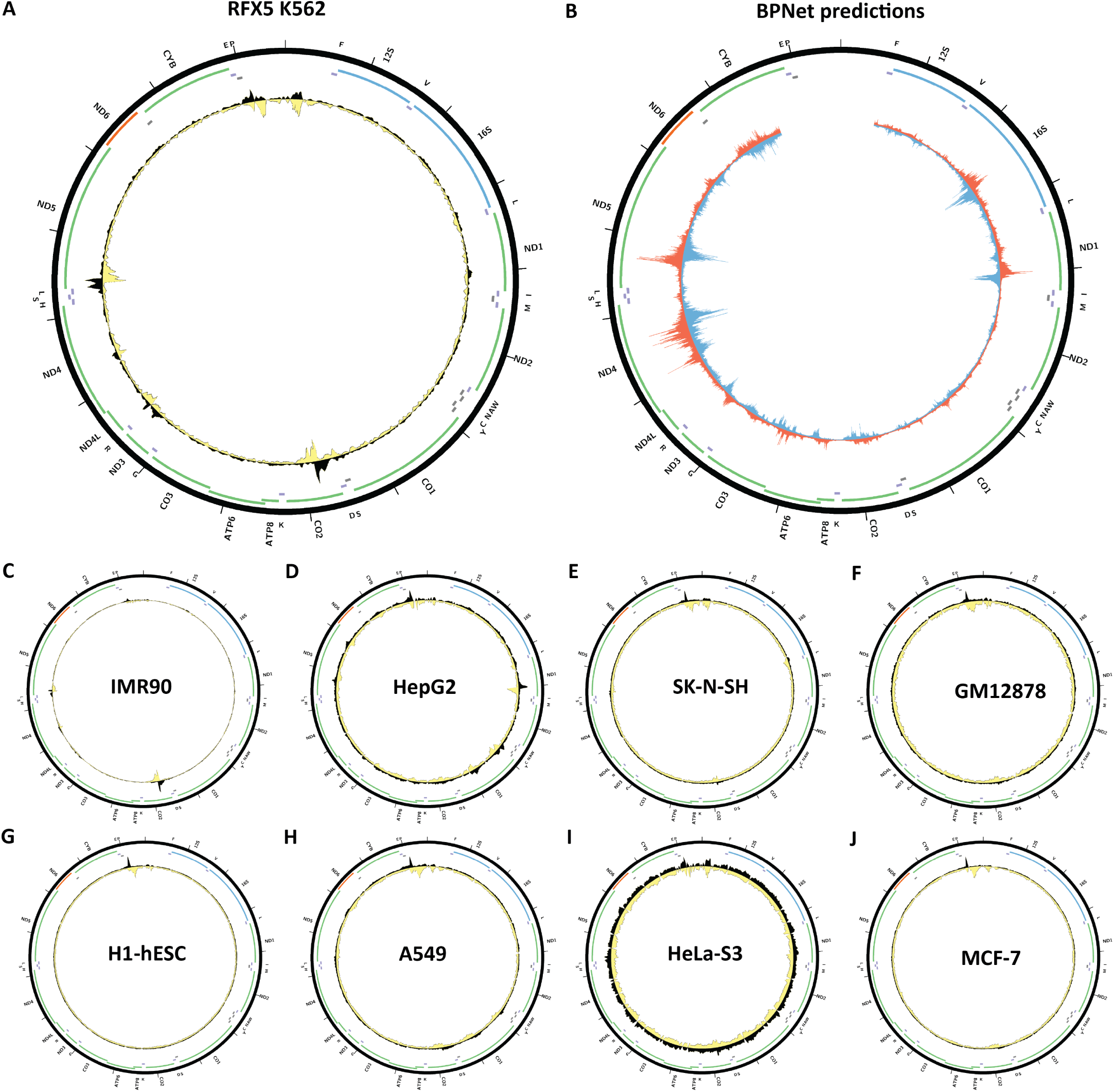
Evidence for mitochondrial genome occupancy by the RFX5 transcription factor. Black and yellow tracks show the forward- and reverse-strand ChIP-seq coverage over chrM. (A) K562 ChIP-seq (ENCODE ID ENCSR000EGO; antibody: Rockland 200-401-194, Lot ID 14562); (B) BPNet predictions over chrM; (C) IMR-90 ChIP-seq (ENCODE ID ENCSR000EFD; antibody: Rockland 200-401-194, Lot ID 14562); (D) HepG2 ChIP-seq (ENCODE ID ENCSR000EEA; antibody: Rockland 200-401-194, Lot ID 14562); (E) SK-N-SH ChIP-seq (ENCODE ID ENCSR000EHY; antibody: Rockland 200-401-194, Lot ID 14562); (F) GM12878 ChIP-seq (ENCODE ID ENCSR000DZW; antibody: Rockland 200-401-194, Lot ID 14562); (G) H1-hESC ChIP-seq (ENCODE ID ENCSR000ECF; antibody: Rockland 200-401-194, Lot ID 14562); (H) A549 ChIP-seq (ENCODE ID ENCSR064LJN; antibody: Rockland 200-401-194, Lot ID 14562); (I) HeLa-S3 ChIP-seq (ENCODE ID ENCSR000ECX; antibody: Rockland 200-401-194, Lot ID 14562); (J) MCF-7 ChIP-seq (ENCODE ID ENCSR924TVL; antibody: Rockland 200-401-194, Lot ID 14562).

**Supplementary Figure 30:**
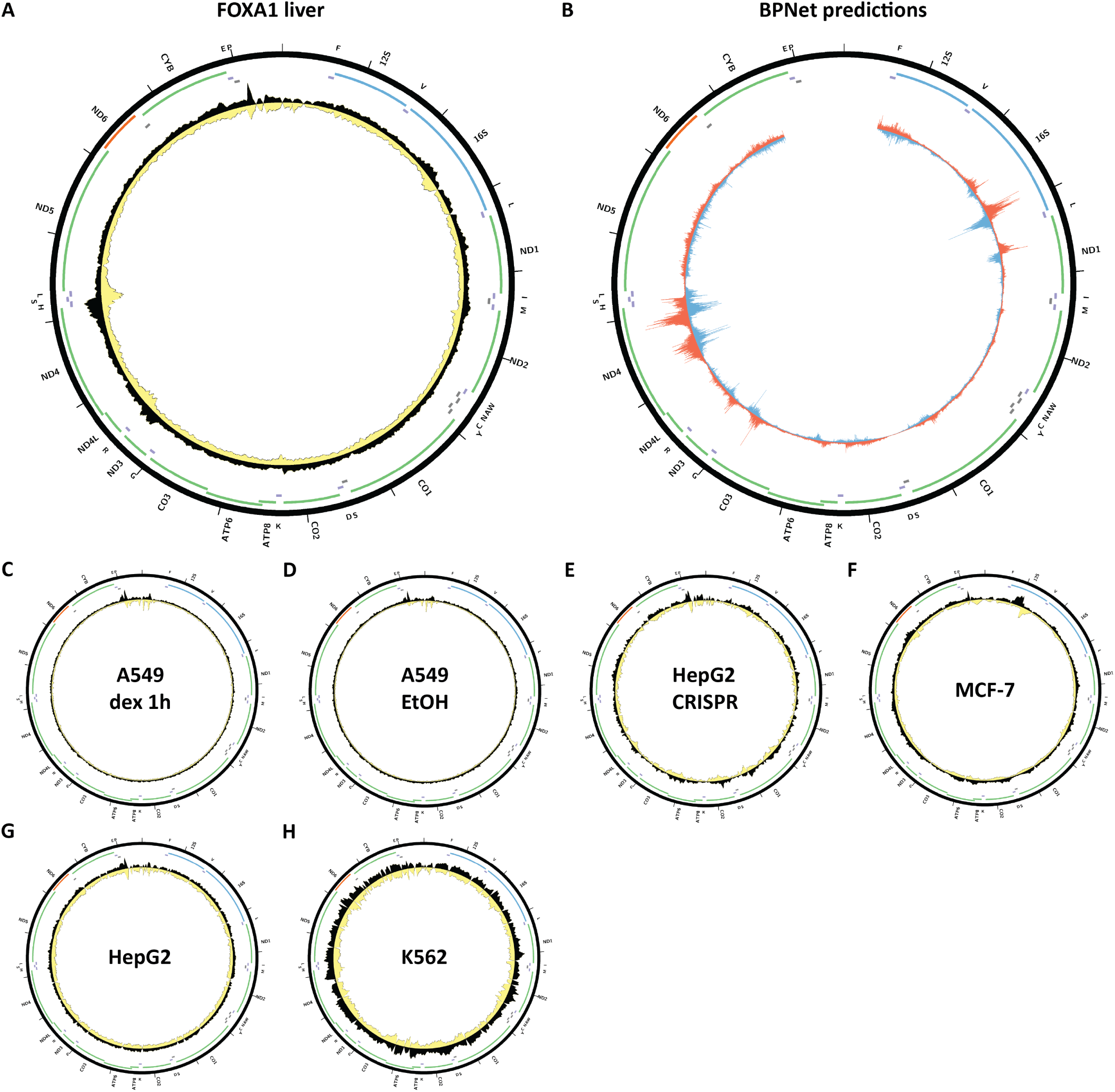
Evidence for mitochondrial genome occupancy by the FOXA1 transcription factor. Black and yellow tracks show the forward- and reverse-strand ChIP-seq coverage over chrM. (A) Liver ChIP-seq (ENCODE ID ENCSR324RCI; antibody: Santa Cruz Biotech sc-6553, Lot ID H1209); (B) BPNet predictions over chrM (ENCODE ID ENCSR578NTH); (C) A549 Dex 1 hours ChIP-seq (ENCODE ID ENCSR000BPX; antibody: Santa Cruz Biotech sc-101058, Lot ID A2706); (D) A549 EtOH 1 hour ChIP-seq (ENCODE ID ENCSR000BRD; antibody: Santa Cruz Biotech sc-101058, Lot ID A2706); (E) HepG2 CETCH-seq (ENCODE ID ENCSR865RXA); (F) MCF-7 ChIP-seq (ENCODE ID ENCSR126YEB; antibody: GeneTex GTX100308, Lot ID 39435); (G) HepG2 ChIP-seq (ENCODE ID ENCSR000BLE; antibody: Santa Cruz Biotech sc-6553, Lot ID H1209); (H) K562 ChIP-seq (ENCODE ID ENCSR819LHG; antibody: GeneTex GTX100308, Lot ID 39435).

**Supplementary Figure 31:**
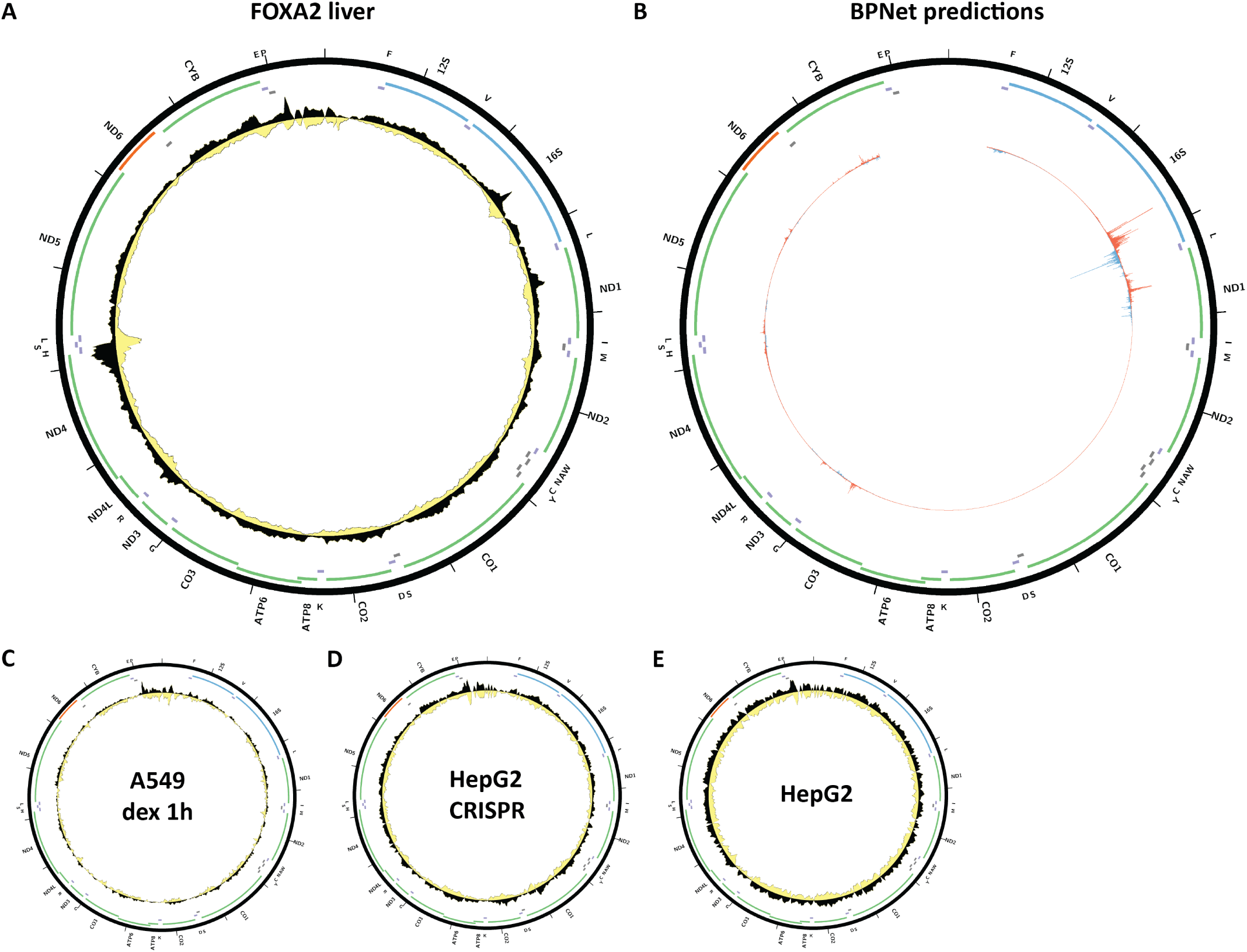
Evidence for mitochondrial genome occupancy by the FOXA2transcription factor. Black and yellow tracks show the forward- and reverse-strand ChIP-seq coverage over chrM. (A) Liver ChIP-seq (ENCODE ID ENCSR080XEY; antibody: Santa Cruz Biotech sc-6554, Lot ID L1409); (B) BPNet predictions over chrM (ENCODE ID ENCSR318CMD); (C) A549 Dex 1 hours ChIP-seq (ENCODE ID ENCSR000BRE; antibody: Santa Cruz Biotech sc-6554, Lot ID L1409); (D) HepG2 CETCH-seq (ENCODE ID ENCSR490AMH); (E) HepG2 ChIP-seq (ENCODE ID ENCSR000BNI; antibody: Santa Cruz Biotech sc-6554, Lot ID L1409).

**Supplementary Figure 32:**
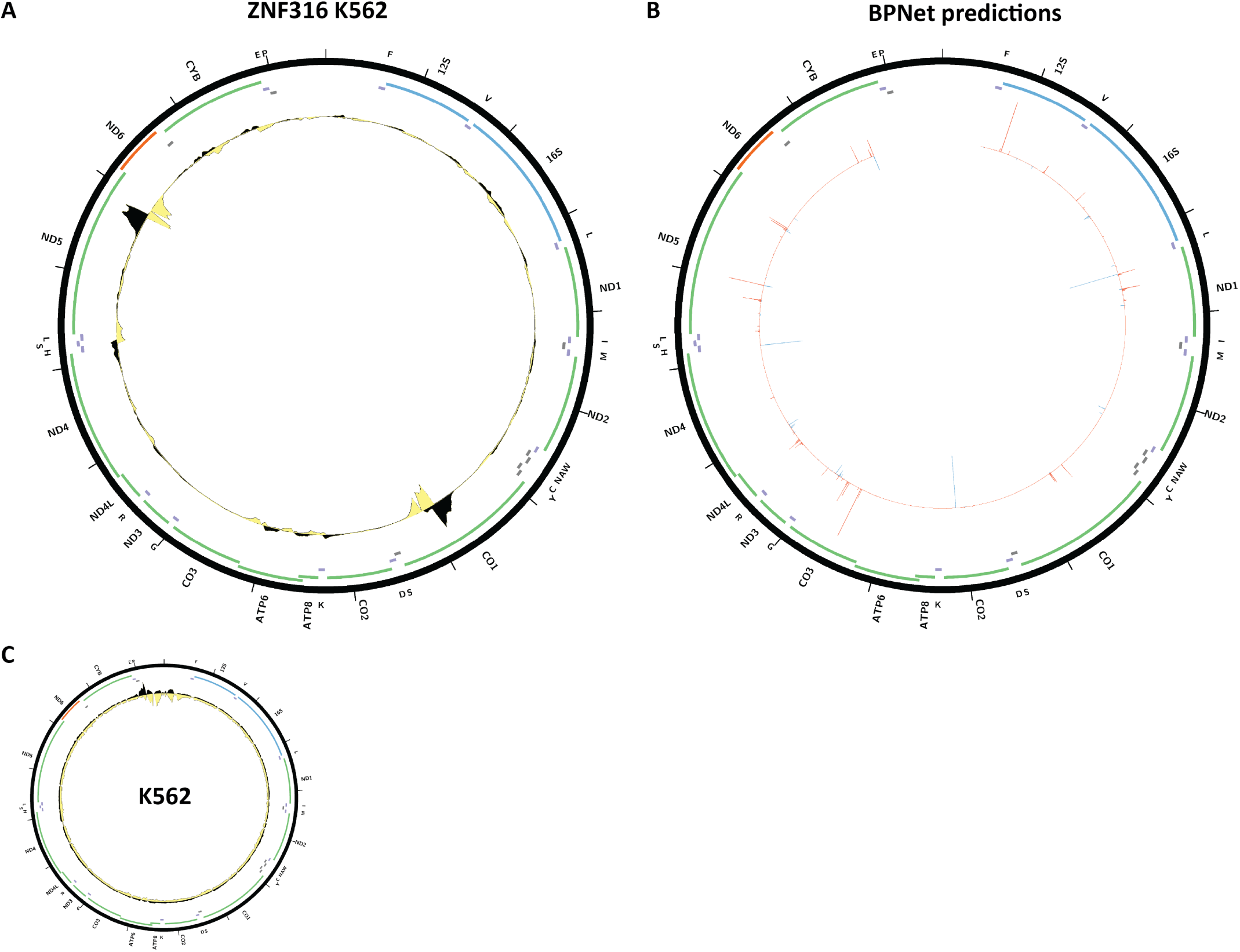
Evidence for mitochondrial genome occupancy by the ZNF316transcription factor. Black and yellow tracks show the forward- and reverse-strand ChIP-seq coverage over chrM. (A) K562 ChIP-seq (ENCODE ID ENCSR167KBO; antibody: Bethyl Labs A303-248A); (B) BPNet predictions over chrM; (C) K562 ChIP-seq (ENCODE ID ENCSR200JYP; antibody: Bethyl Labs A303-249A).

**Supplementary Figure 33:**
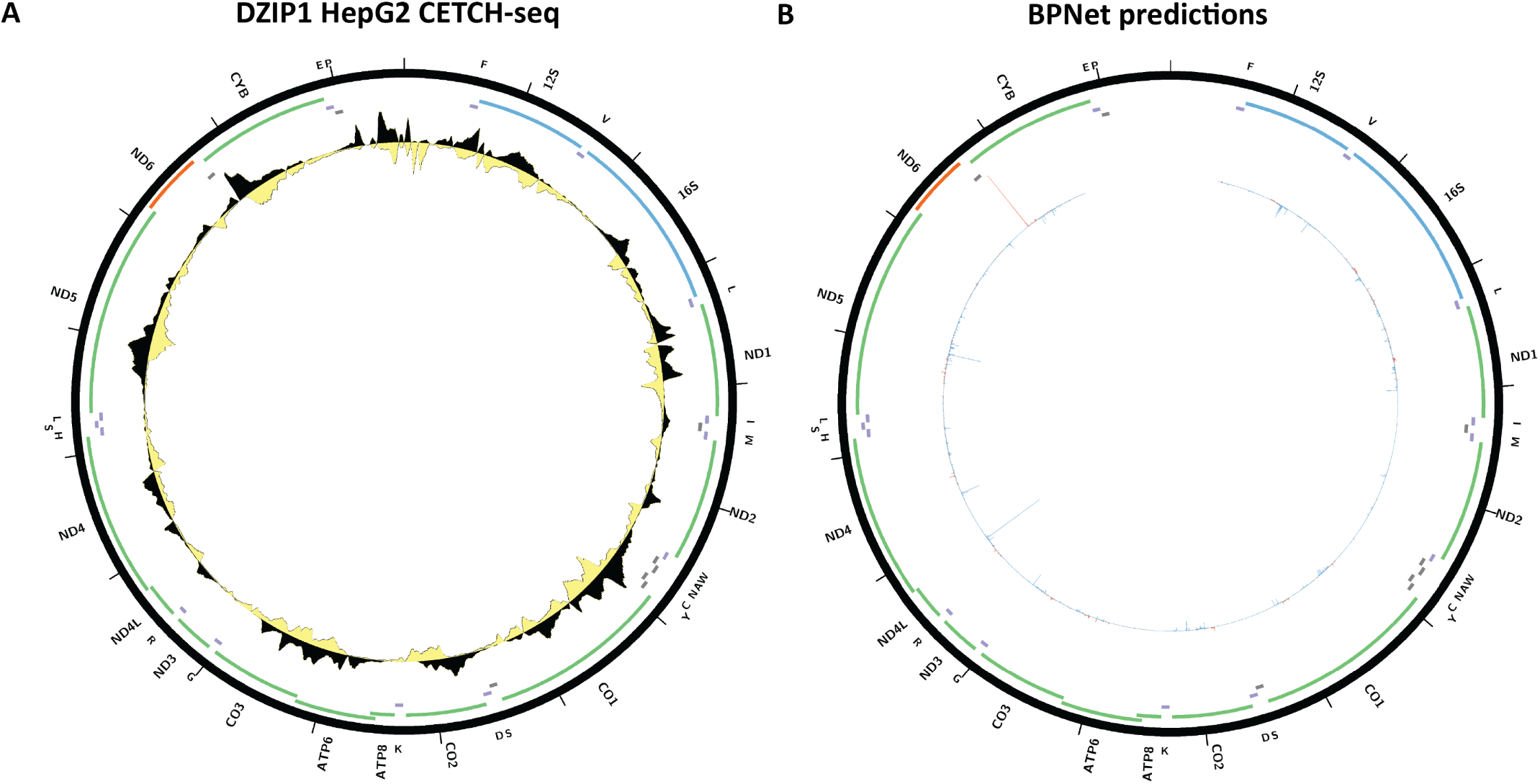
Evidence for mitochondrial genome occupancy by the DZIP1 transcription factor. Black and yellow tracks show the forward- and reverse-strand ChIP-seq coverage over chrM. (A) HepG2 CETCH- seq (ENCODE ID ENCSR895KNN); (B) BPNet predictions over chrM.

**Supplementary Figure 34:**
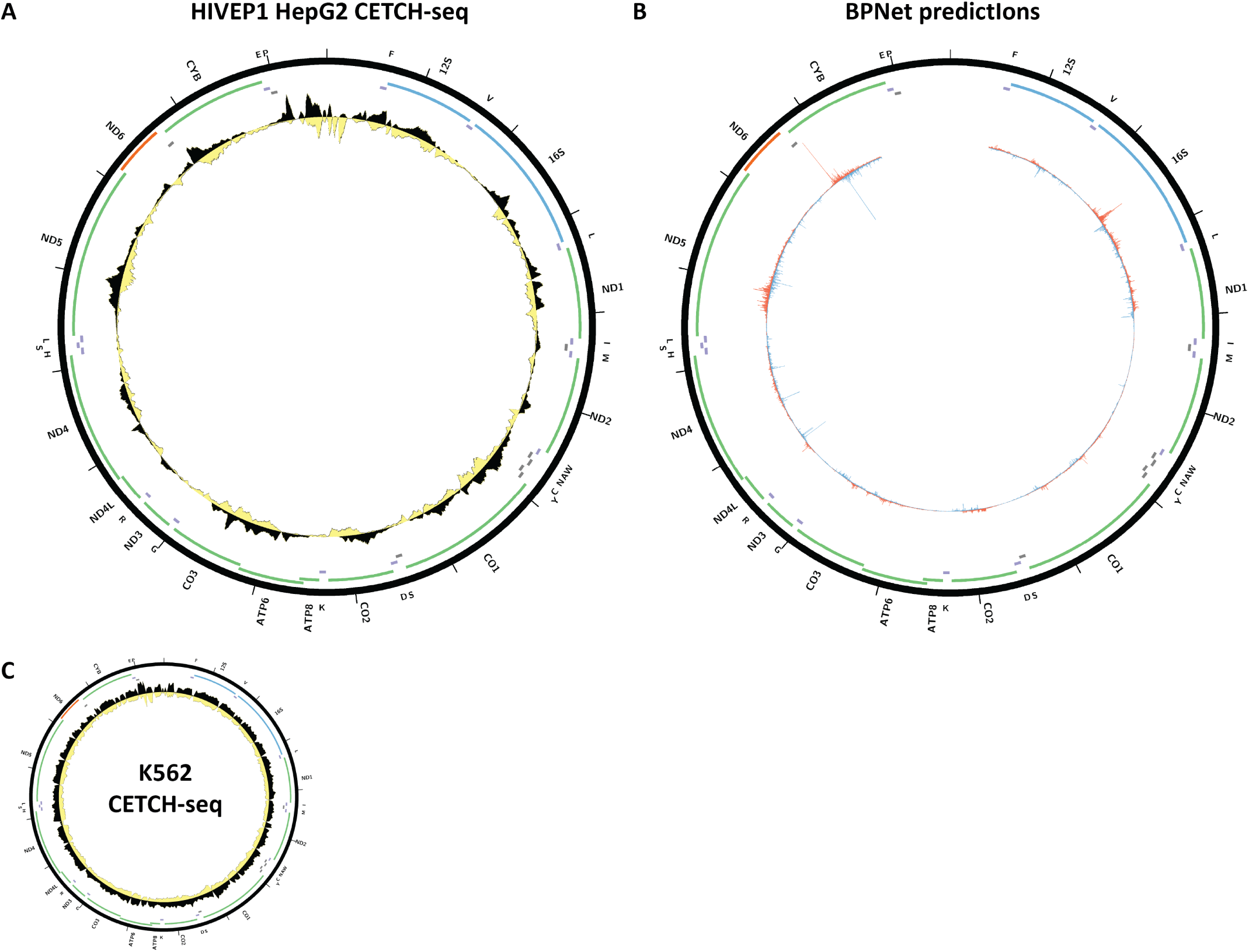
Evidence for mitochondrial genome occupancy by the HIVEP1 transcription factor. Black and yellow tracks show the forward- and reverse-strand ChIP-seq coverage over chrM. (A) HepG2 CETCH-seq (ENCODE ID ENCSR697WMX); (B) BPNet predictions over chrM (ENCODE ID ENCSR272FZX); (C) K562 CETCH-seq (ENCODE ID ENCSR947PJZ).

**Supplementary Figure 35:**
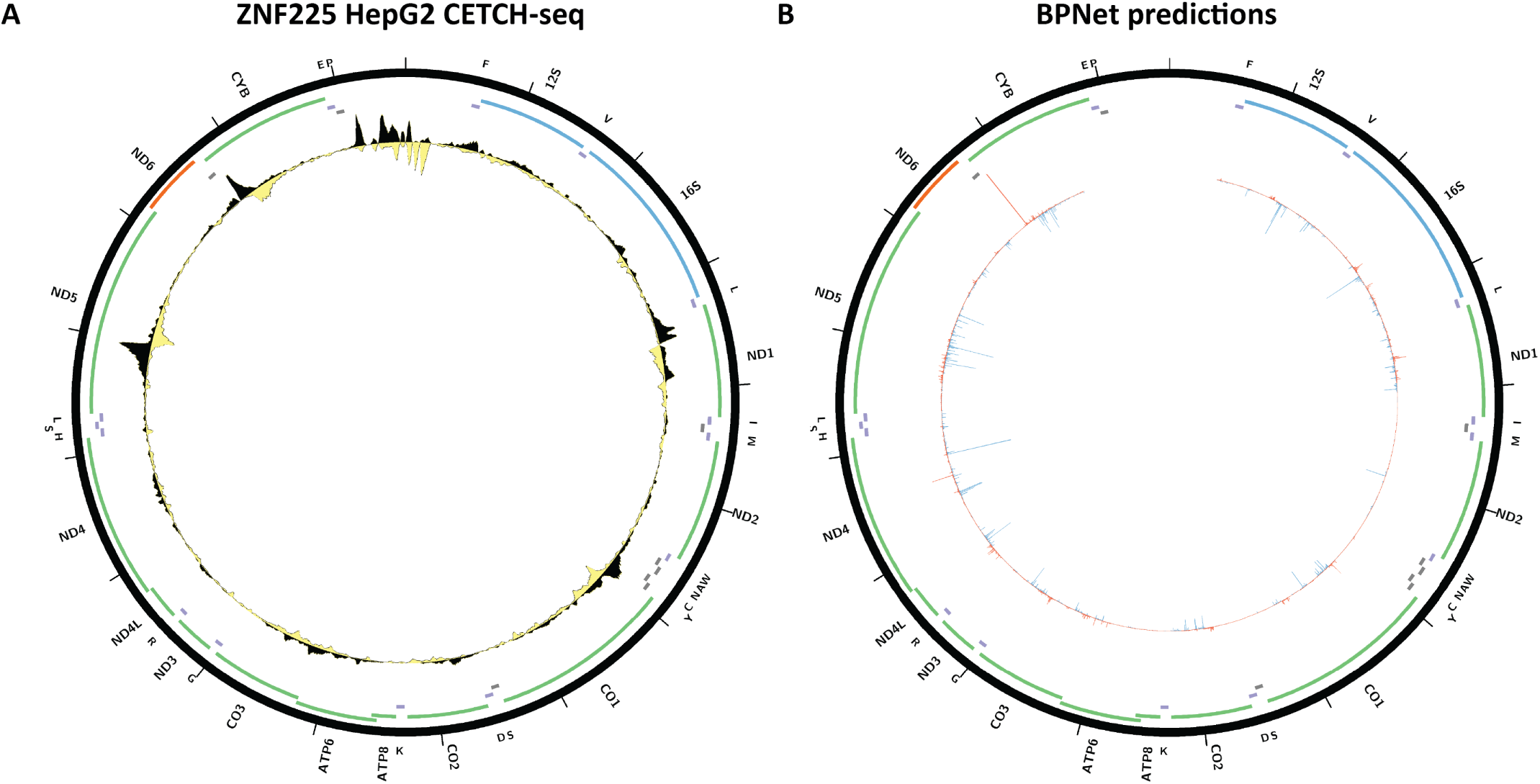
Evidence for mitochondrial genome occupancy by the ZNF225 transcription factor. Black and yellow tracks show the forward- and reverse-strand ChIP-seq coverage over chrM. (A) HepG2 CETCH- seq (ENCODE ID ENCSR075PWK); (B) BPNet predictions over chrM.

**Supplementary Figure 36:**
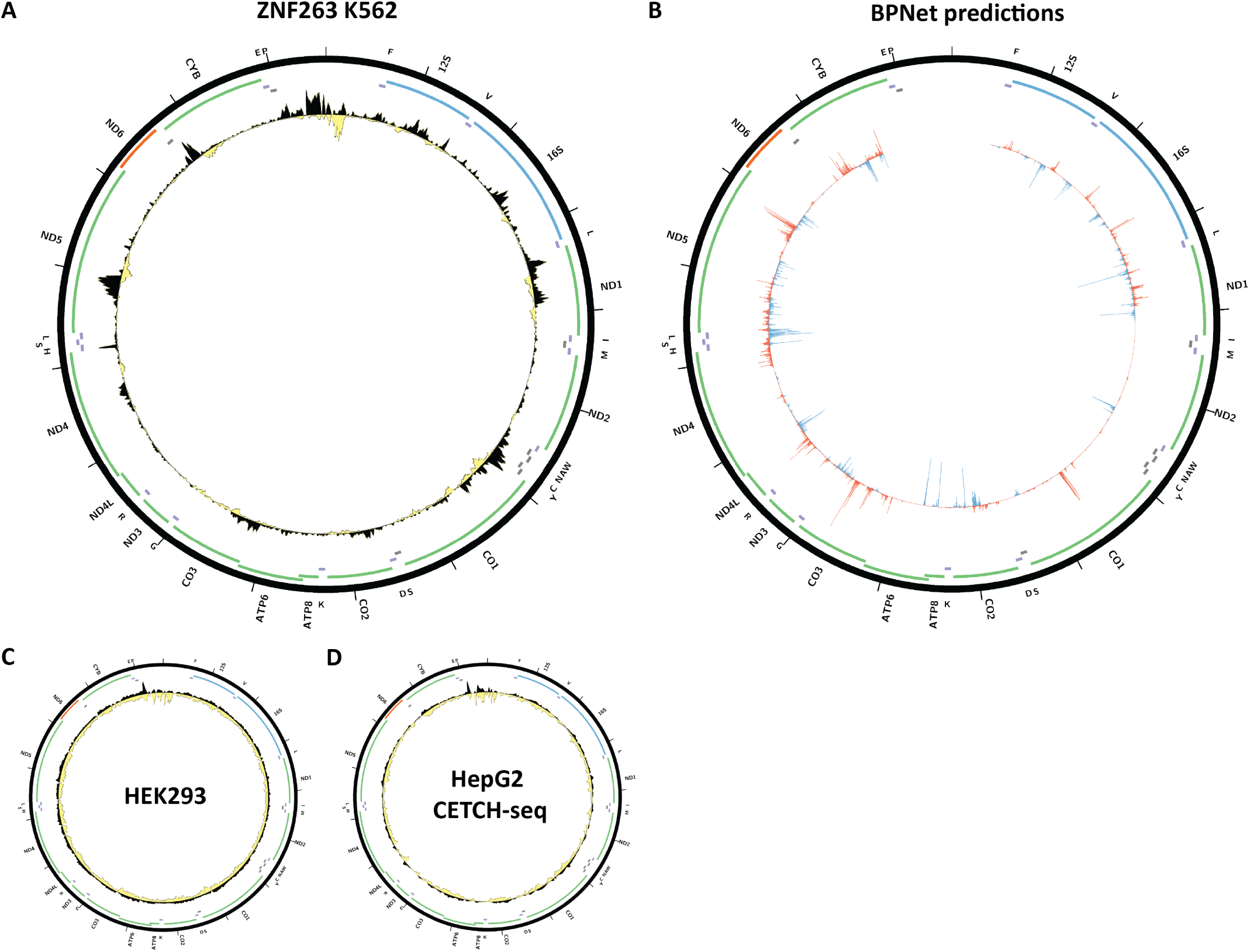
Evidence for mitochondrial genome occupancy by the ZNF263transcription factor. Black and yellow tracks show the forward- and reverse-strand ChIP-seq coverage over chrM. (A) K562 ChIP-seq (ENCODE ID ENCSR000EWN; antibody: Novus H00010127-A01); (B) BPNet predictions over chrM; (C) HEK293 ChIP-seq (ENCODE ID ENCSR000EVD; antibody: Novus H00010127-A01); (D) HepG2 CETCH-seq.

**Supplementary Figure 37:**
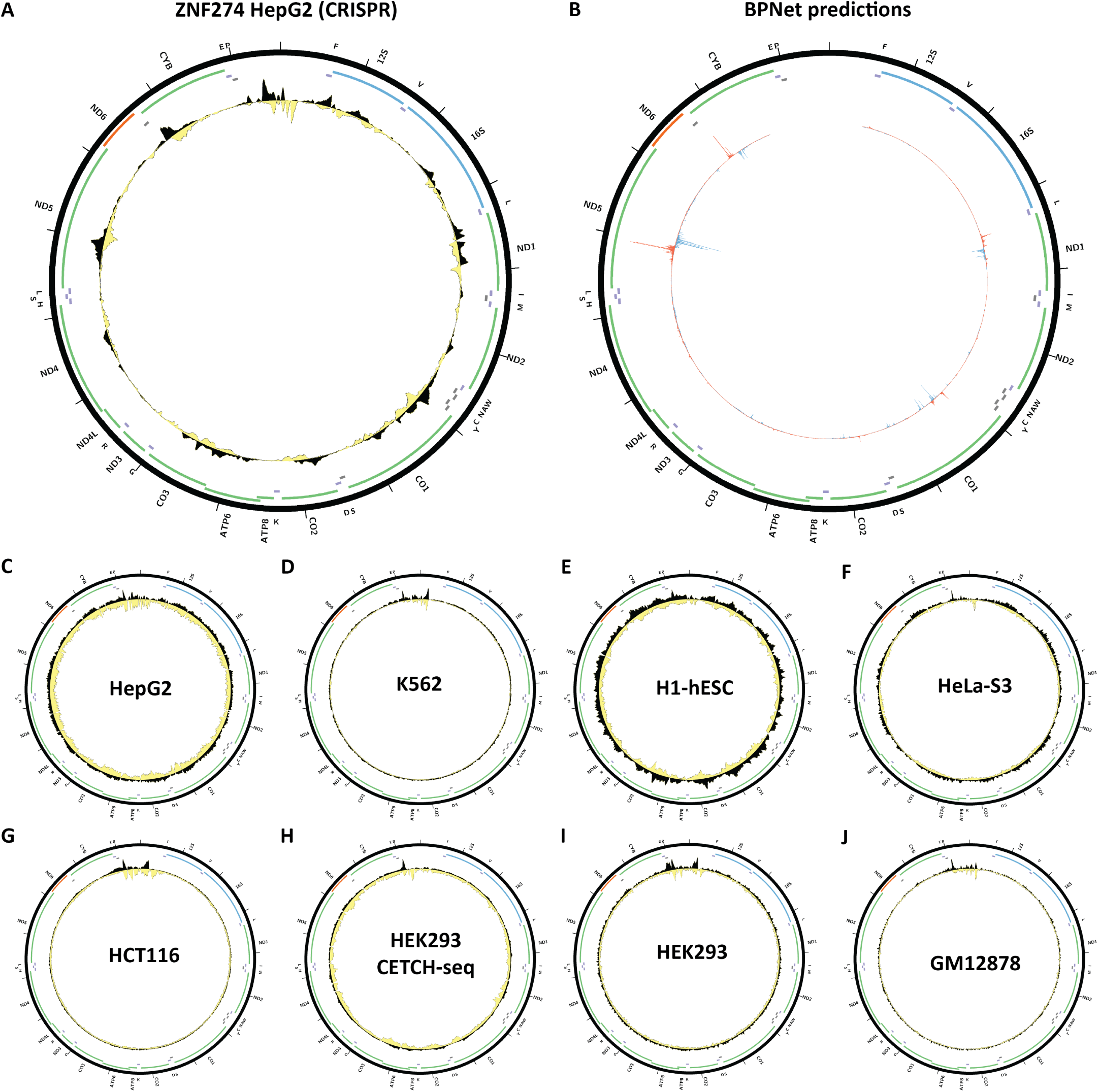
Evidence for mitochondrial genome occupancy by the ZNF274 transcription factor. Black and yellow tracks show the forward- and reverse-strand ChIP-seq coverage over chrM. (A) HepG2 CETCH-seq (ENCODE ID ENCSR871VNN); (B) BPNet predictions over chrM; (C) HepG2 ChIP-seq (ENCODE ID ENCSR000EVR; antibody: Abnova H00010782-A01, Lot ID 060729QCS1); (D) K562 ChIP-seq (ENCODE ID ENCSR000EVX; anti-body: Abnova H00010782-M01, Lot ID 08064-4C12); (E) H1-hESC ChIP-seq (ENCODE ID ENCSR000EUN; antibody: Abnova H00010782-M01, Lot ID 08064-4C12); (F) HeLa-S3 ChIP-seq (ENCODE ID ENCSR000EVG; antibody: Ab-nova H00010782-A01, Lot ID 060729QCS1); (G) HCT116 ChIP-seq (ENCODE ID ENCSR101FJM; antibody: Abnova H00010782-A01, Lot ID 060729QCS1); (H) HEK293 CETCH-seq (ENCODE ID ENCSR178QVJ); (I) HEK293 ChIP-seq (ENCODE ID ENCSR000FCI; antibody: Abnova H00010782-A01, Lot ID 060729QCS1); (J) GM12878 ChIP-seq (ENCODE ID ENCSR000EUK; antibody: Abnova H00010782-A01, Lot ID 060729QCS1).

**Supplementary Figure 38:**
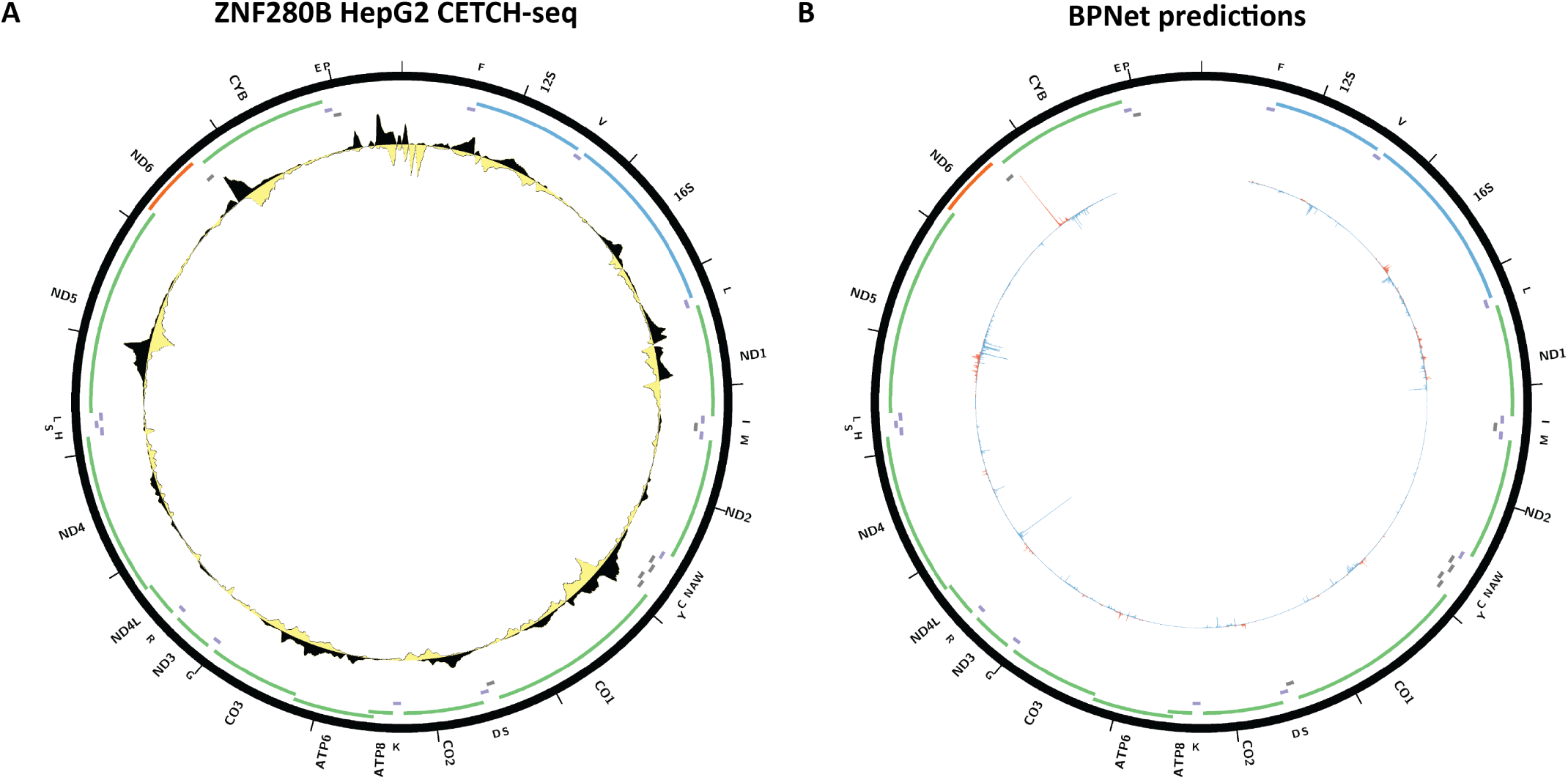
Evidence for mitochondrial genome occupancy by the ZNF280B transcription factor. Black and yellow tracks show the forward- and reverse-strand ChIP-seq coverage over chrM. (A) HepG2 CETCH- seq (ENCODE ID ENCSR940EZR); (B) BPNet predictions over chrM.

**Supplementary Figure 39:**
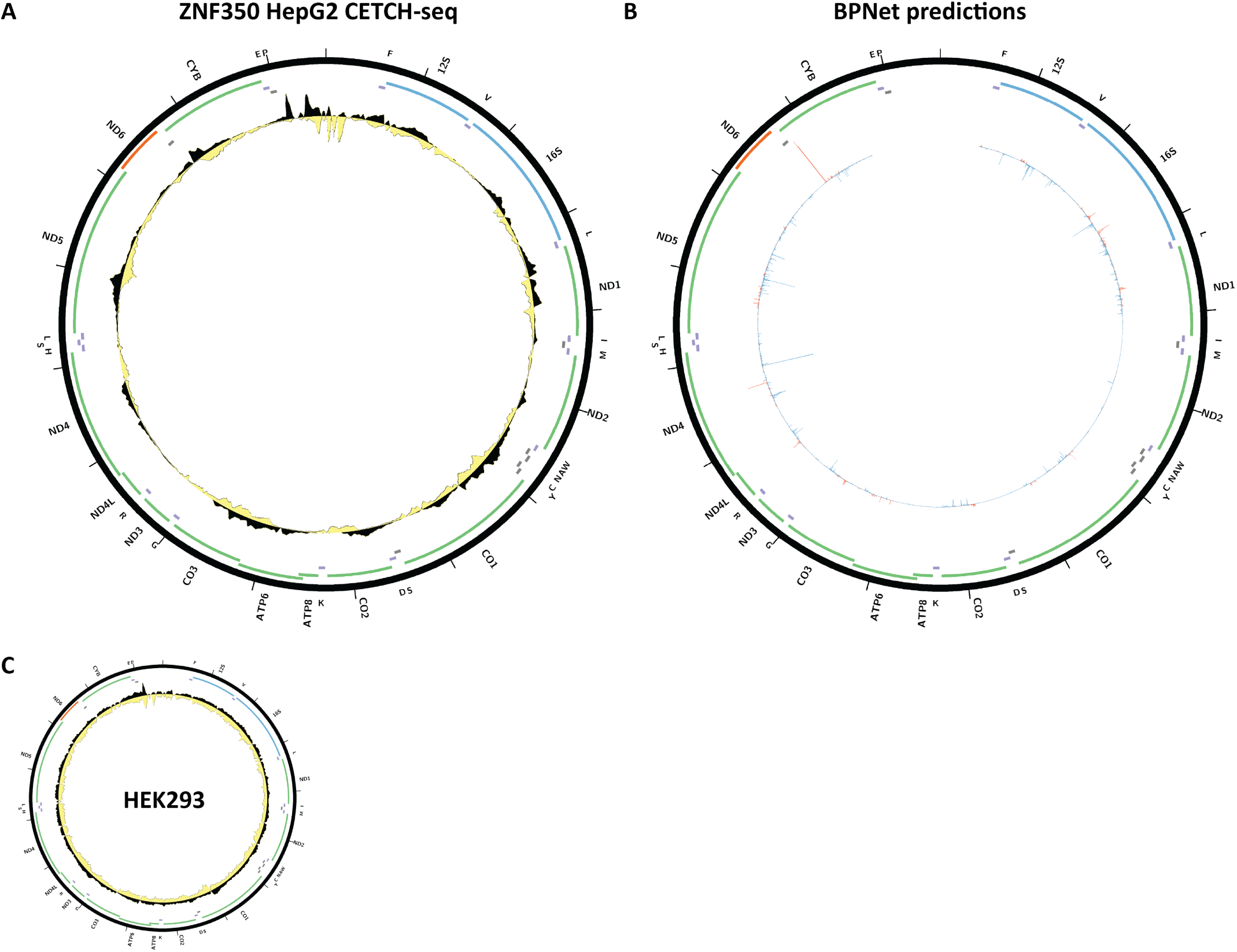
Evidence for mitochondrial genome occupancy by the ZNF350 transcription factor. Black and yellow tracks show the forward- and reverse-strand ChIP-seq coverage over chrM. (A) HepG2 CETCH-seq (ENCODE ID ENCSR842SRB); (B) BPNet predictions over chrM; (C) HEK293 CETCH-seq (ENCODE ID ENCSR854ORP).

**Supplementary Figure 40:**
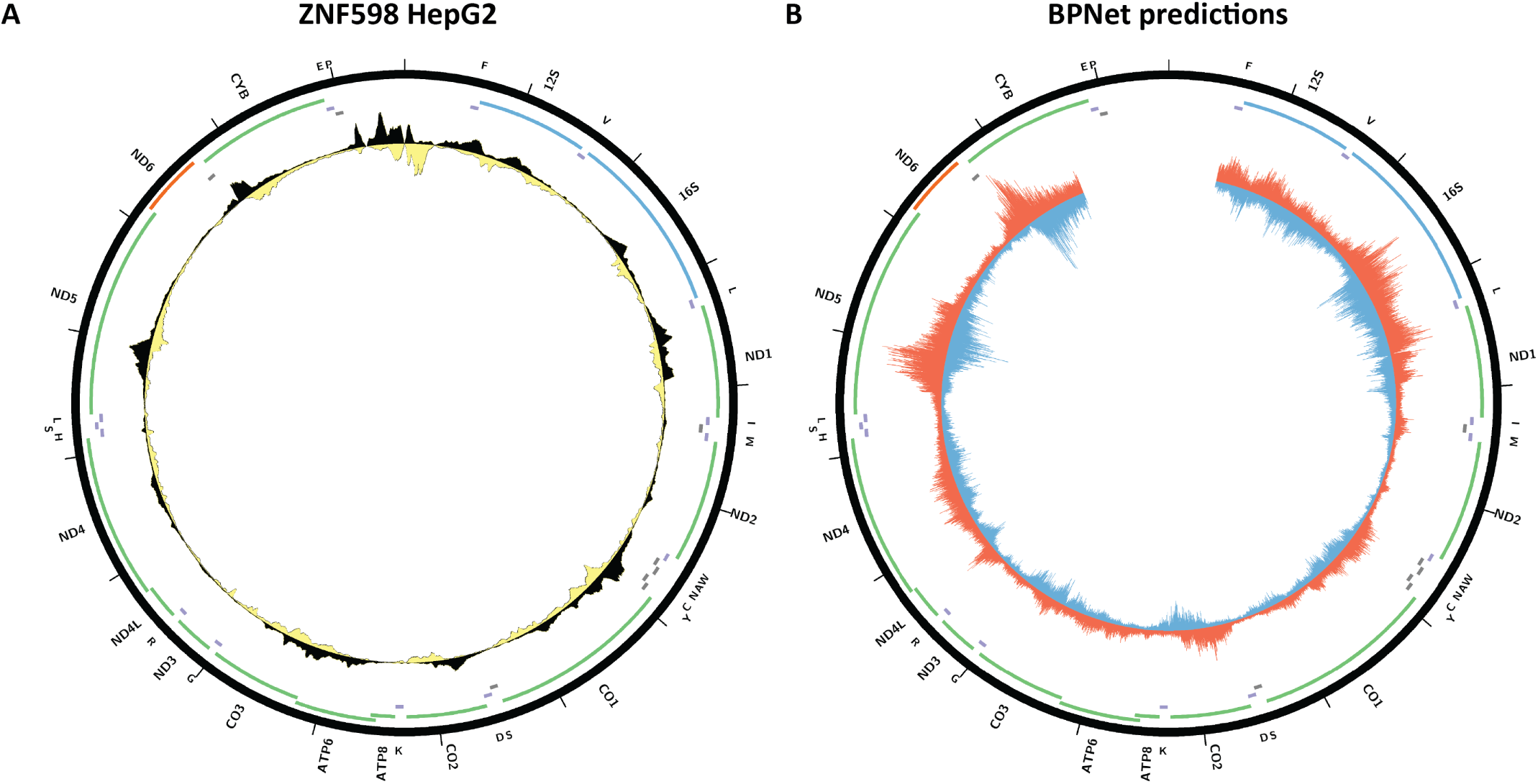
Evidence for mitochondrial genome occupancy by the ZNF598transcription factor. Black and yellow tracks show the forward- and reverse-strand ChIP-seq coverage over chrM. (A) HepG2 ChIP-seq (ENCODE ID ENCSR173NAL; antibody: Sigma F1804, Lot ID SLBK1346V); (B) BPNet predictions over chrM.

**Supplementary Figure 41:**
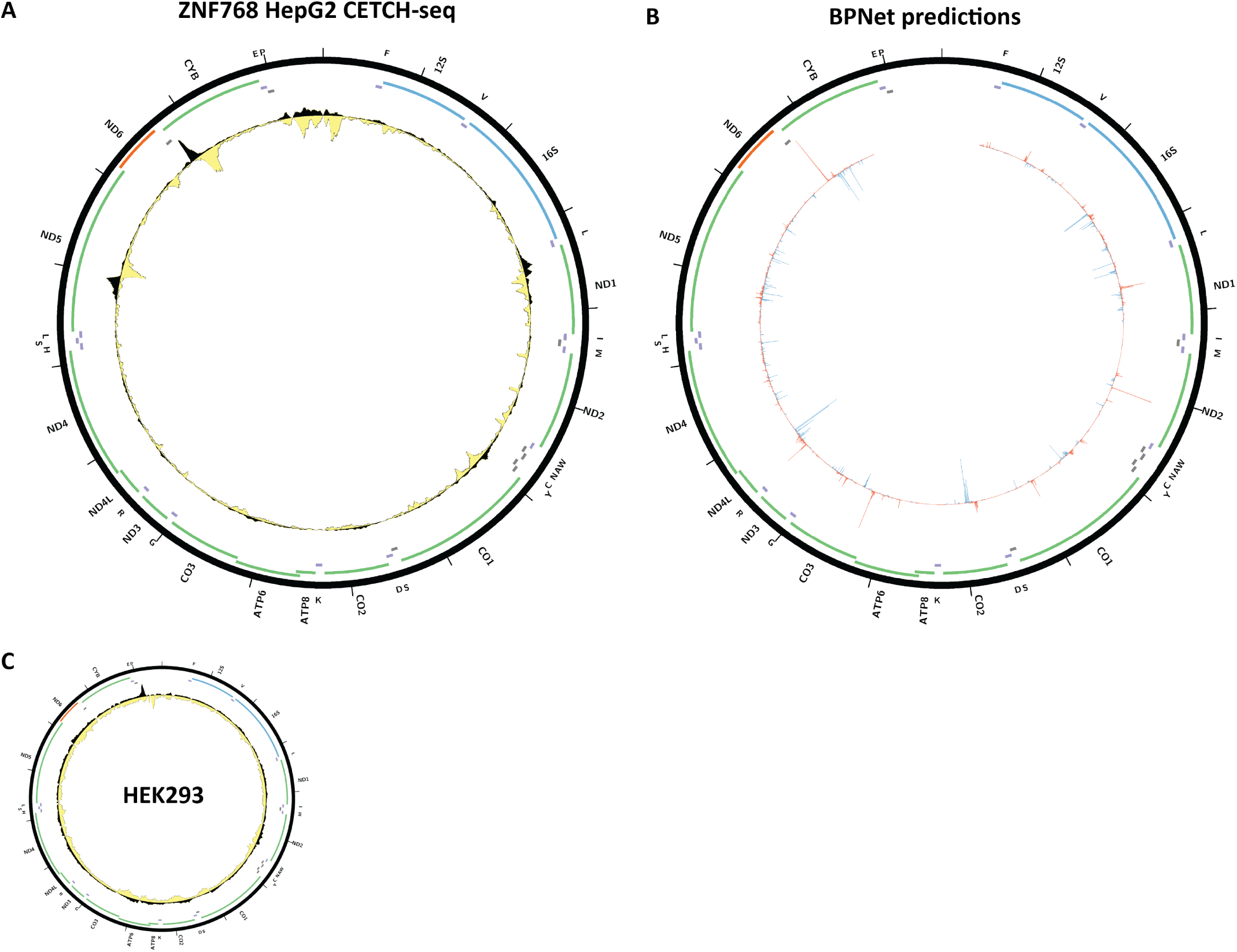
Evidence for mitochondrial genome occupancy by the ZNF768 transcription factor. Black and yellow tracks show the forward- and reverse-strand ChIP-seq coverage over chrM. (A) HepG2 CETCH-seq (ENCODE ID ENCSR181ABP); (B) BPNet predictions over chrM; (C) HEK293 CETCH-seq (ENCODE ID ENCSR070HWF).

**Supplementary Figure 42:**
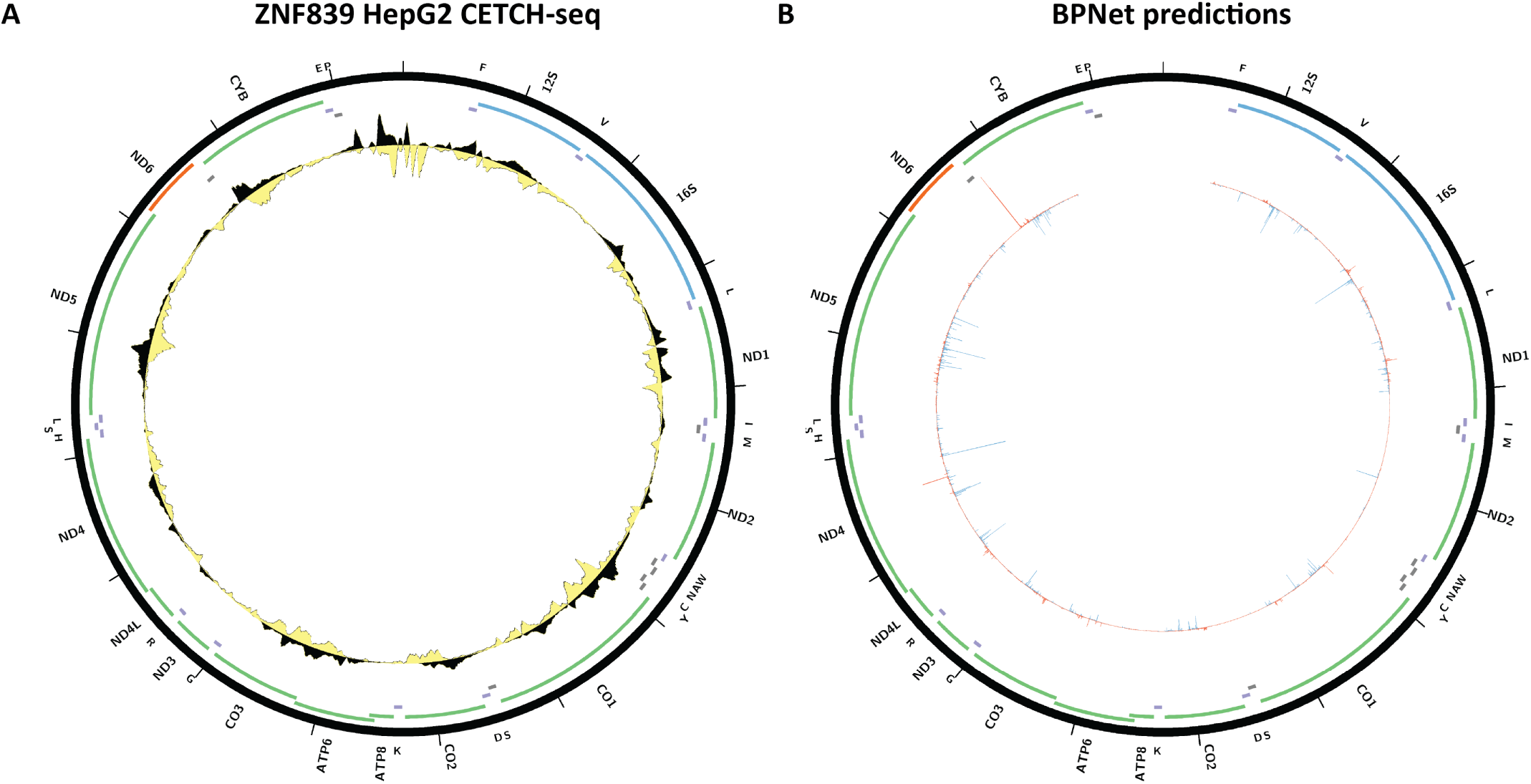
Evidence for mitochondrial genome occupancy by the ZNF839 transcription factor. Black and yellow tracks show the forward- and reverse-strand ChIP-seq coverage over chrM. (A) HepG2 CETCH- seq (ENCODE ID ENCSR540LPD). (B) BPNet predictions over chrM.

**Supplementary Figure 43:**
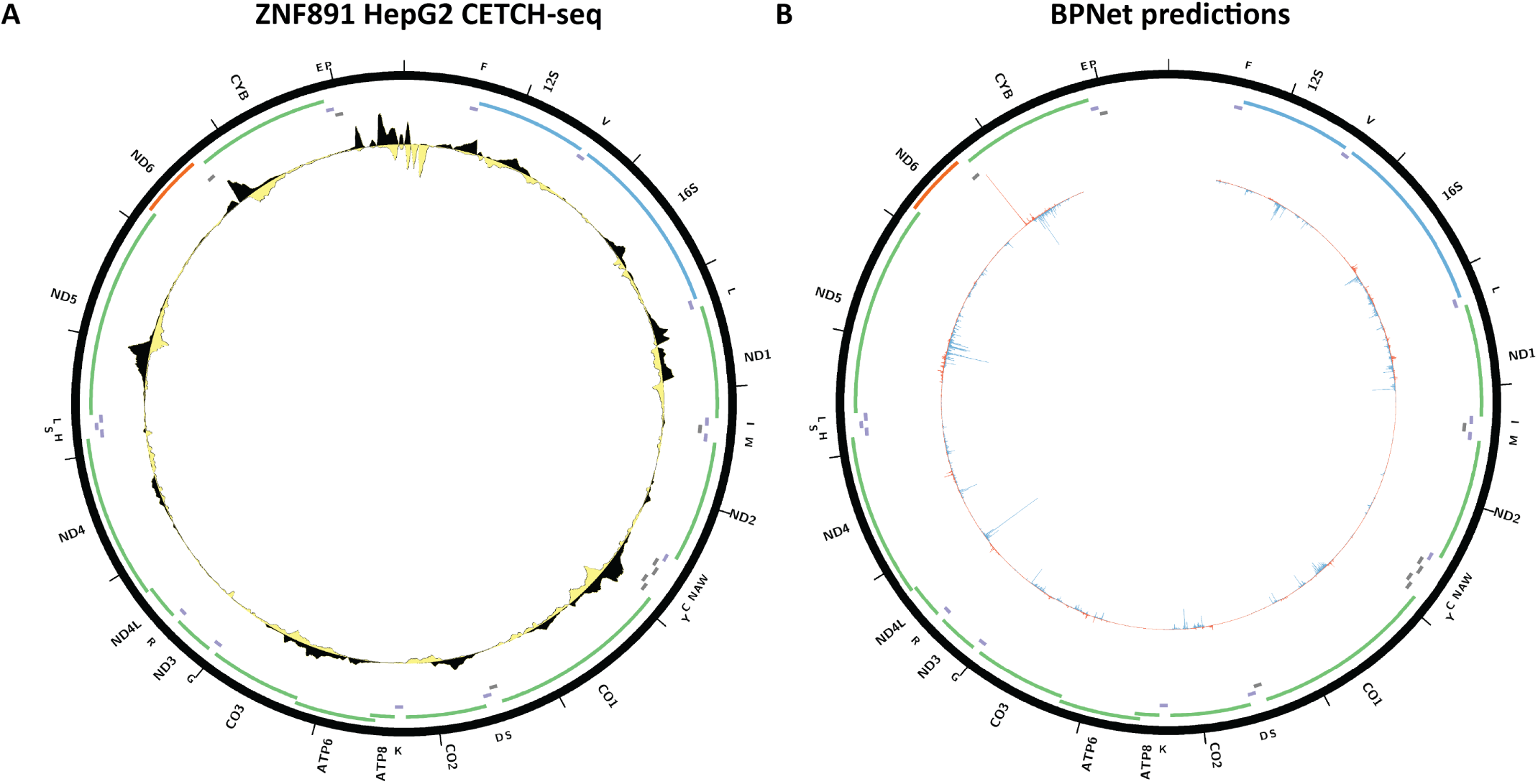
Evidence for mitochondrial genome occupancy by the ZNF891 transcription factor. Black and yellow tracks show the forward- and reverse-strand ChIP-seq coverage over chrM. (A) HepG2 CETCH- seq (ENCODE ID ENCSR020CLV). (B) BPNet predictions over chrM.

**Supplementary Figure 44:**
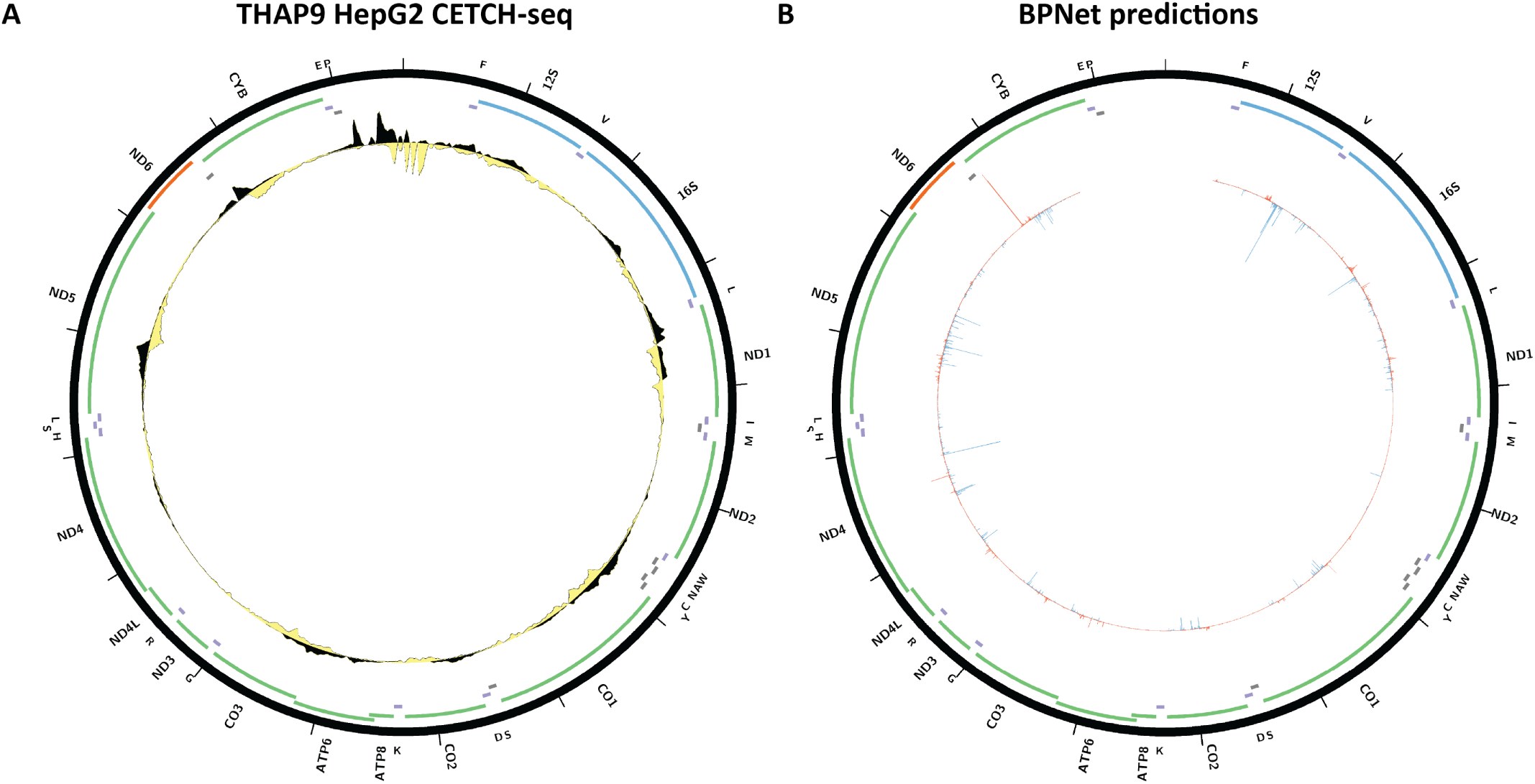
Evidence for mitochondrial genome occupancy by the THAP9 transcription factor. Black and yellow tracks show the forward- and reverse-strand ChIP-seq coverage over chrM. (A) HepG2 CETCH- seq (ENCODE ID ENCSR123GPC); (B) BPNet predictions over chrM.

**Supplementary Figure 45:**
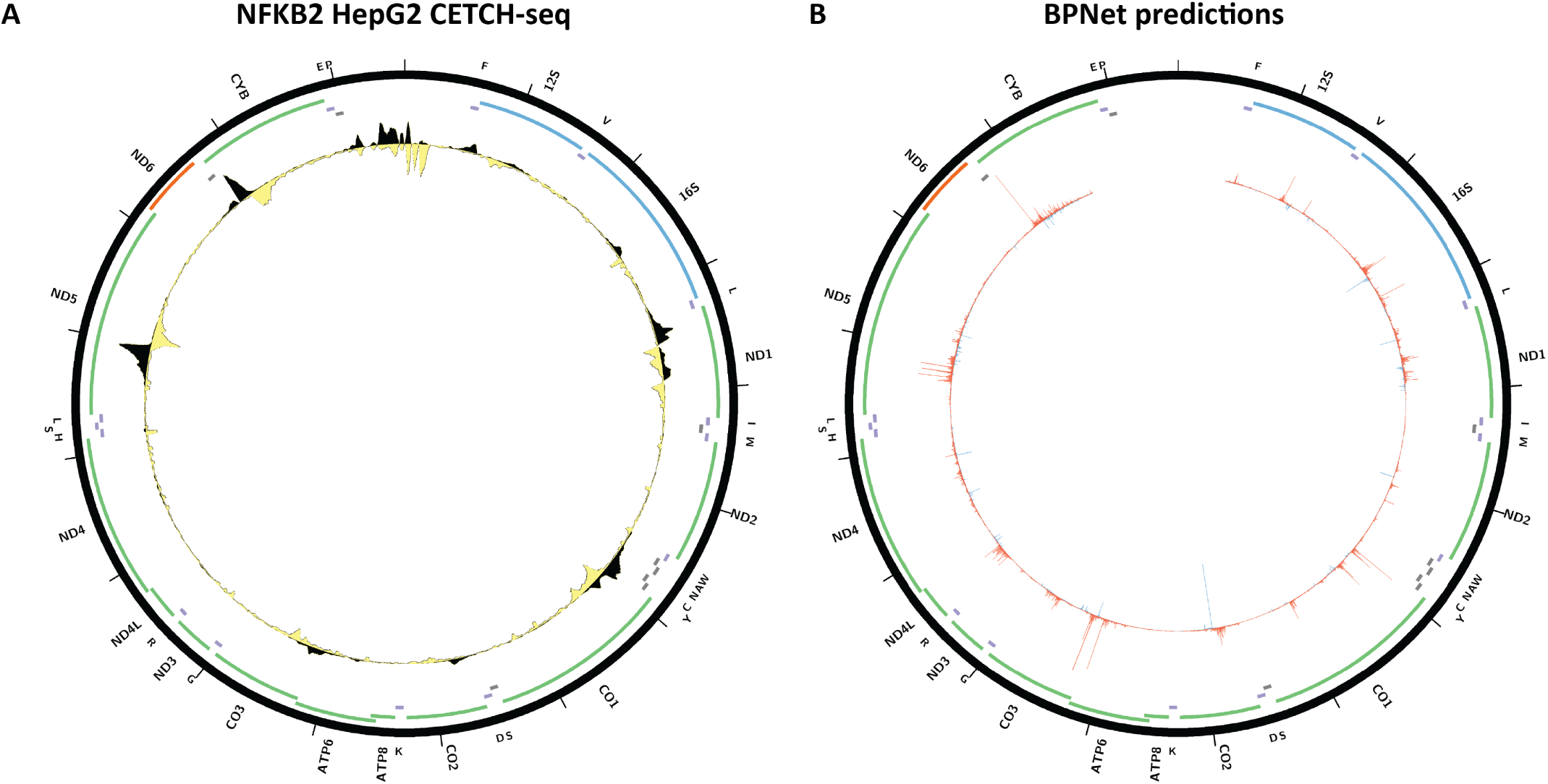
Evidence for mitochondrial genome occupancy by the NFKB2 transcription factor. Black and yellow tracks show the forward- and reverse-strand ChIP-seq coverage over chrM. (A) HepG2 CETCH- seq (ENCODE ID ENCSR164YJZ); (B) BPNet predictions over chrM.

**Supplementary Figure 46:**
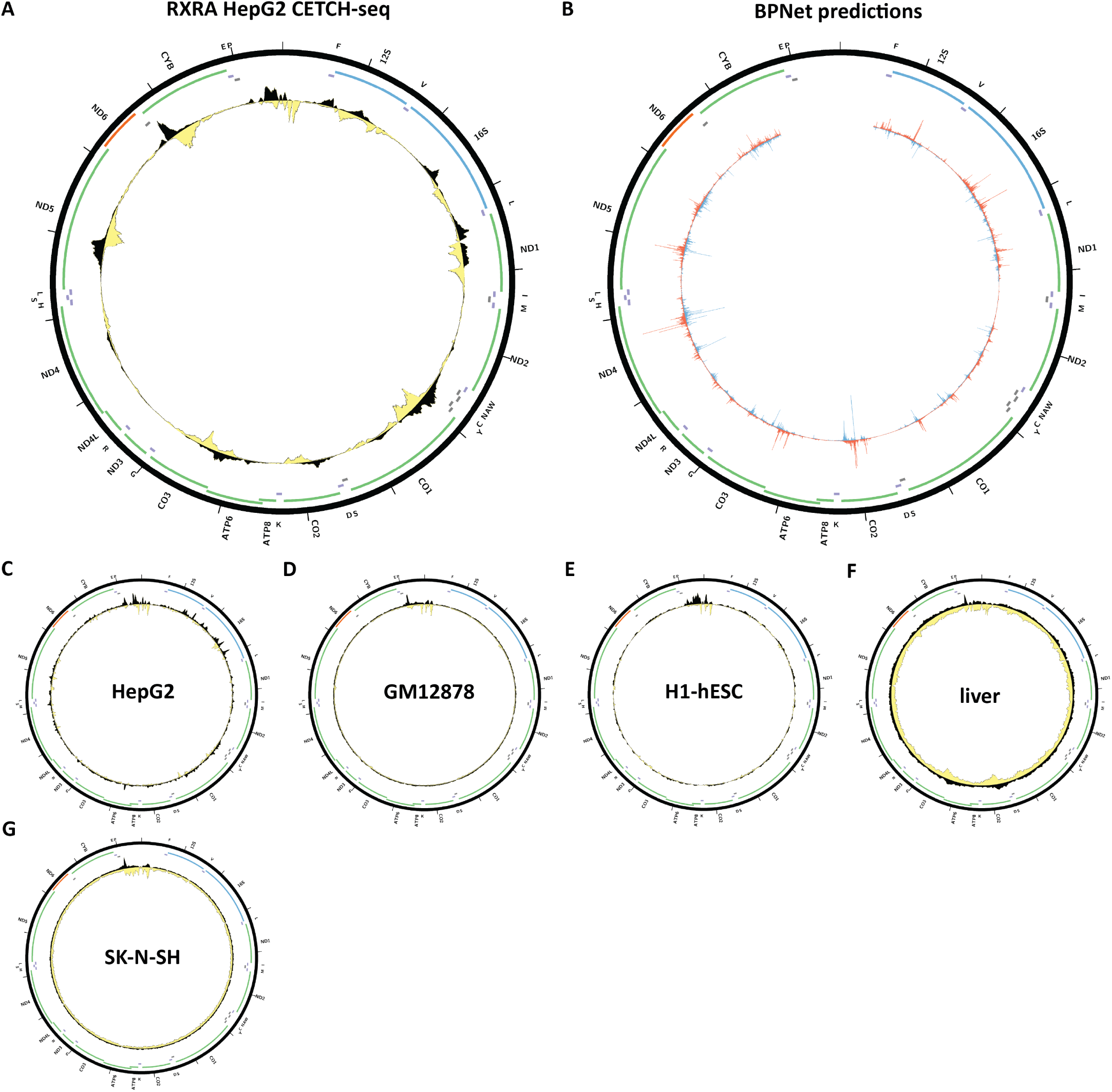
Evidence for mitochondrial genome occupancy by the RXRA transcription factor. Black and yellow tracks show the forward- and reverse-strand ChIP-seq coverage over chrM. (A) HepG2CETCH-seq (ENCODE ID ENCSR416HDG); (B) BPNet predictions over chrM (ENCODE ID ENCSR471TML); (C) HepG2 ChIP-seq (ENCODE ID ENCSR000BHU; antibody: Santa Cruz Biotech sc-553, Lot ID C1811); (D) GM12878 ChIP-seq (ENCODE ID ENCSR000BJD; antibody: Santa Cruz Biotech sc-553, Lot ID C1811); (E) H1-hESC ChIP-seq (ENCODE ID ENCSR000BJW; antibody: Santa Cruz Biotech sc-553, Lot ID C1811); (F) Liver ChIP-seq (ENCODE ID ENCSR352QSB; antibody: Santa Cruz Biotech sc-553, Lot ID C1811); (G) SK-N-SH ChIP-seq (ENCODE ID ENCSR000BVG; antibody: Santa Cruz Biotech sc-553, Lot ID C1811).

**Supplementary Figure 47:**
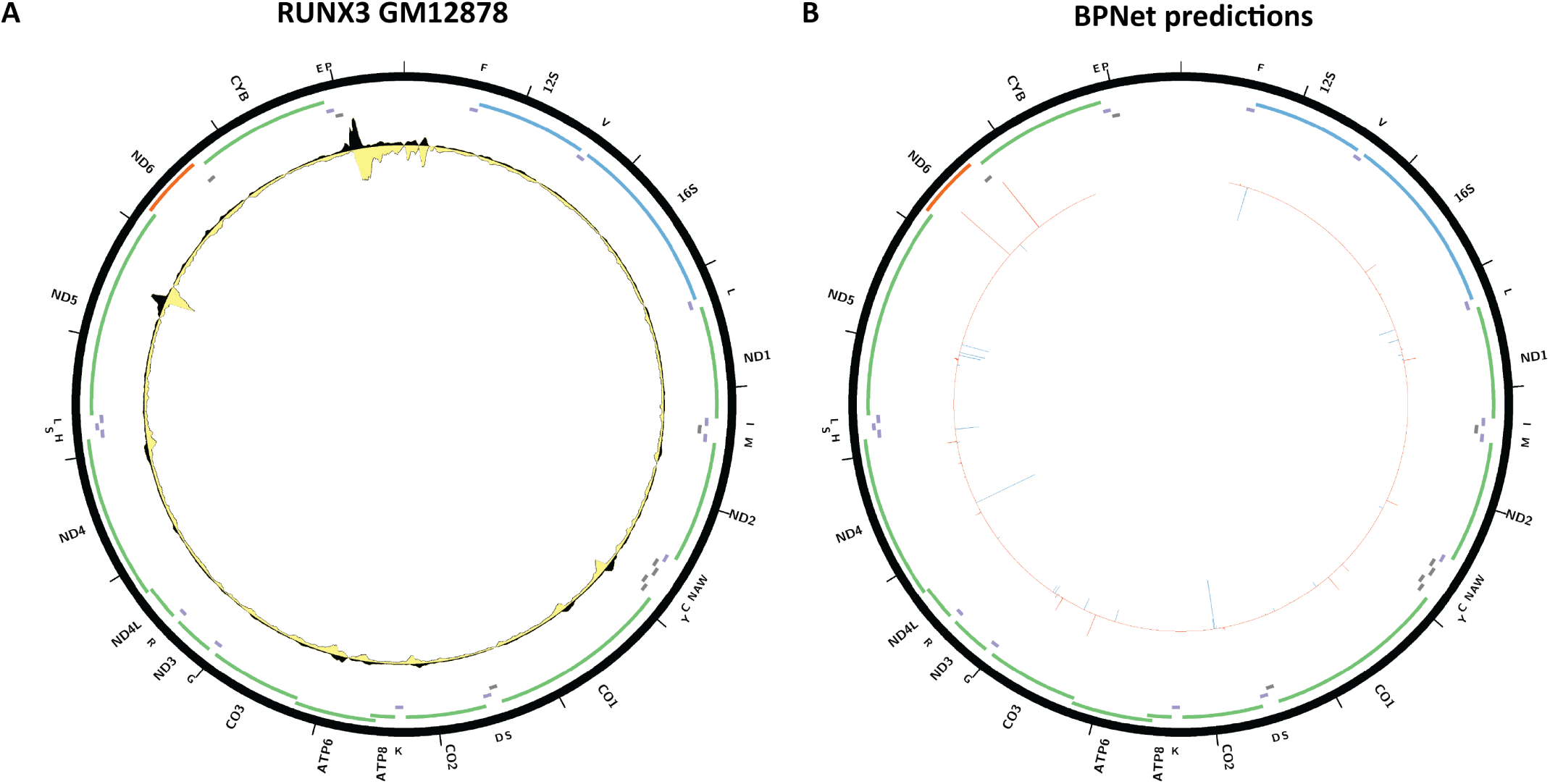
Evidence for mitochondrial genome occupancy by the RUNX3transcription factor. Black and yellow tracks show the forward- and reverse-strand ChIP-seq coverage over chrM. (A) GM12878 ChIP-seq (ENCODE ID ENCSR000BRI; antibody: Santa Cruz Biotech sc-101553, Lot ID B0909); (B) BPNet predictions over chrM (ENCODE ID ENCSR130BYX).

**Supplementary Figure 48:**
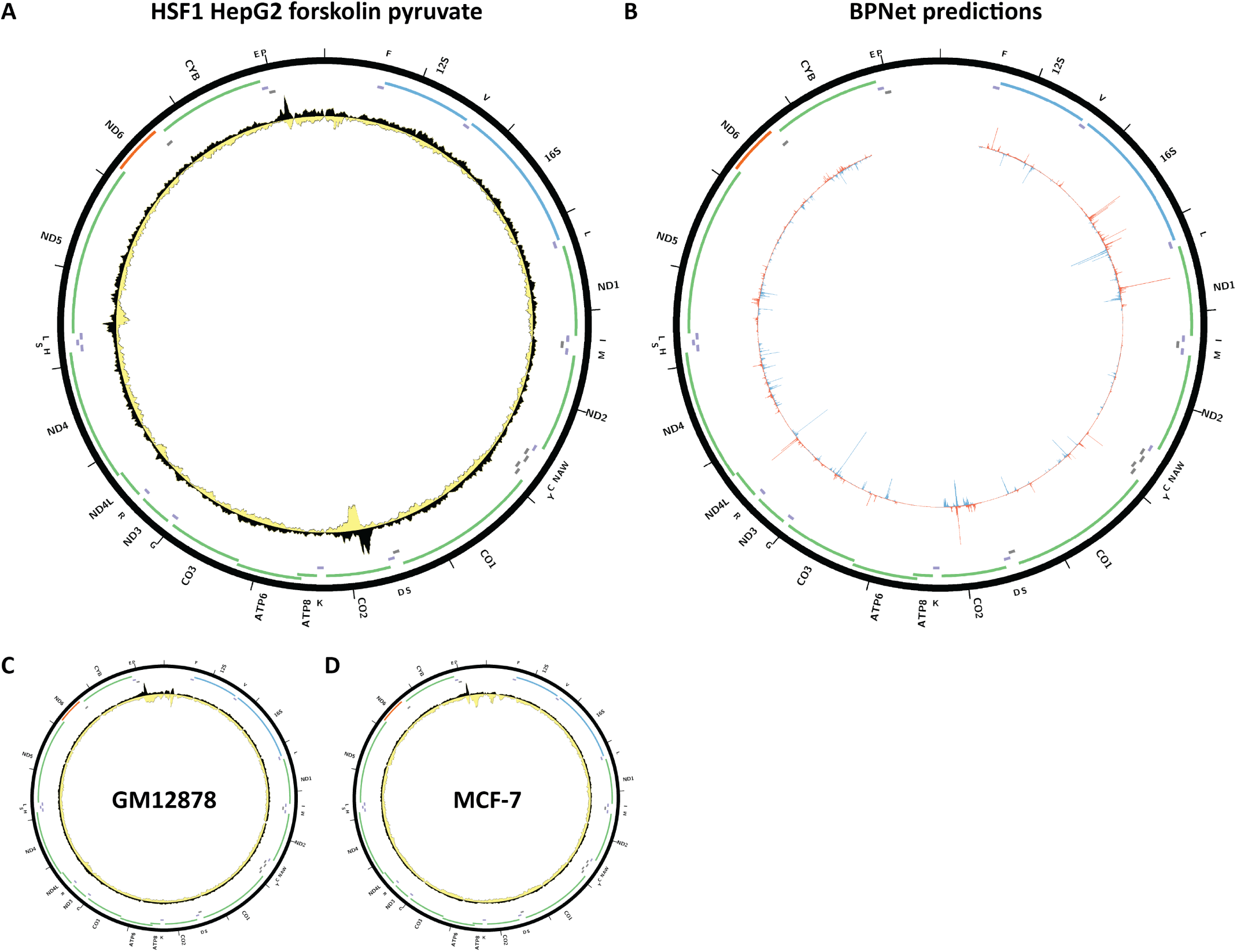
Evidence for mitochondrial genome occupancy by the HSF1 transcription factor. Black and yellow tracks show the forward- and reverse-strand ChIP-seq coverage over chrM. (A) HepG2 forskolin + 1mM pyruvate 6 hours, ChIP-seq (ENCODE ID ENCSR000EET; antibody: Santa Cruz Biotech sc-9144); (B) BPNet predictions over chrM (ENCODE ID ENCSR000EET); (C) GM12878 ChIP-seq (ENCODE ID ENCSR009MBP; antibody: Santa Cruz Biotech sc-9144); (D) MCF-7 ChIP-seq (ENCODE ID ENCSR062HDL; antibody: Santa Cruz Biotech sc- 9144).

**Supplementary Figure 49:**
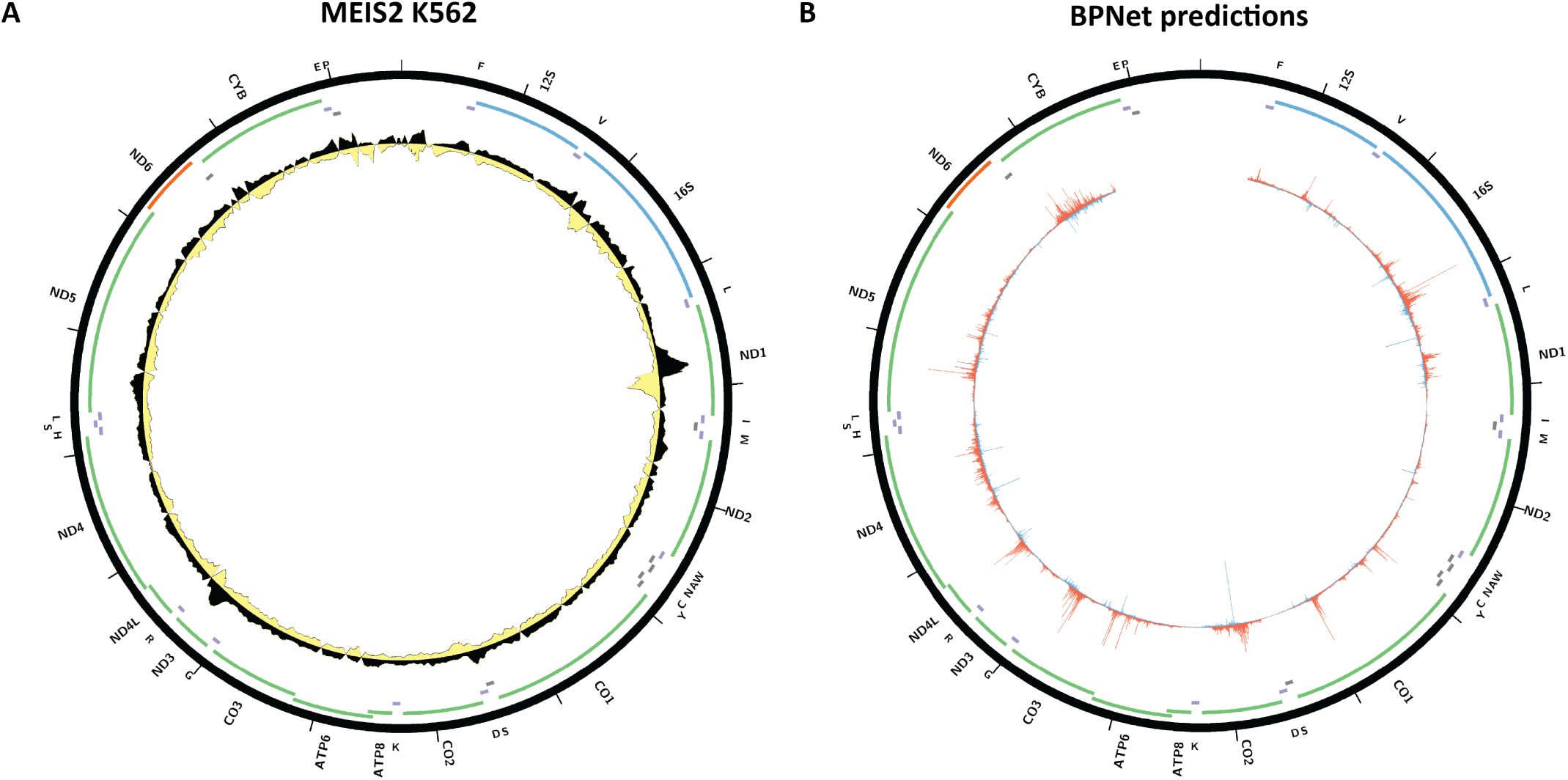
Evidence for mitochondrial genome occupancy by the MEIS2transcription factor. Black and yellow tracks show the forward- and reverse-strand ChIP-seq coverage over chrM. (A) K562 ChIP-seq (ENCODE ID ENCSR851BNE; antibody: Sigma HPA003256, Lot ID R89735); (B) BPNet predictions over chrM (ENCODE ID ENCSR603QNK).

**Supplementary Figure 50:**
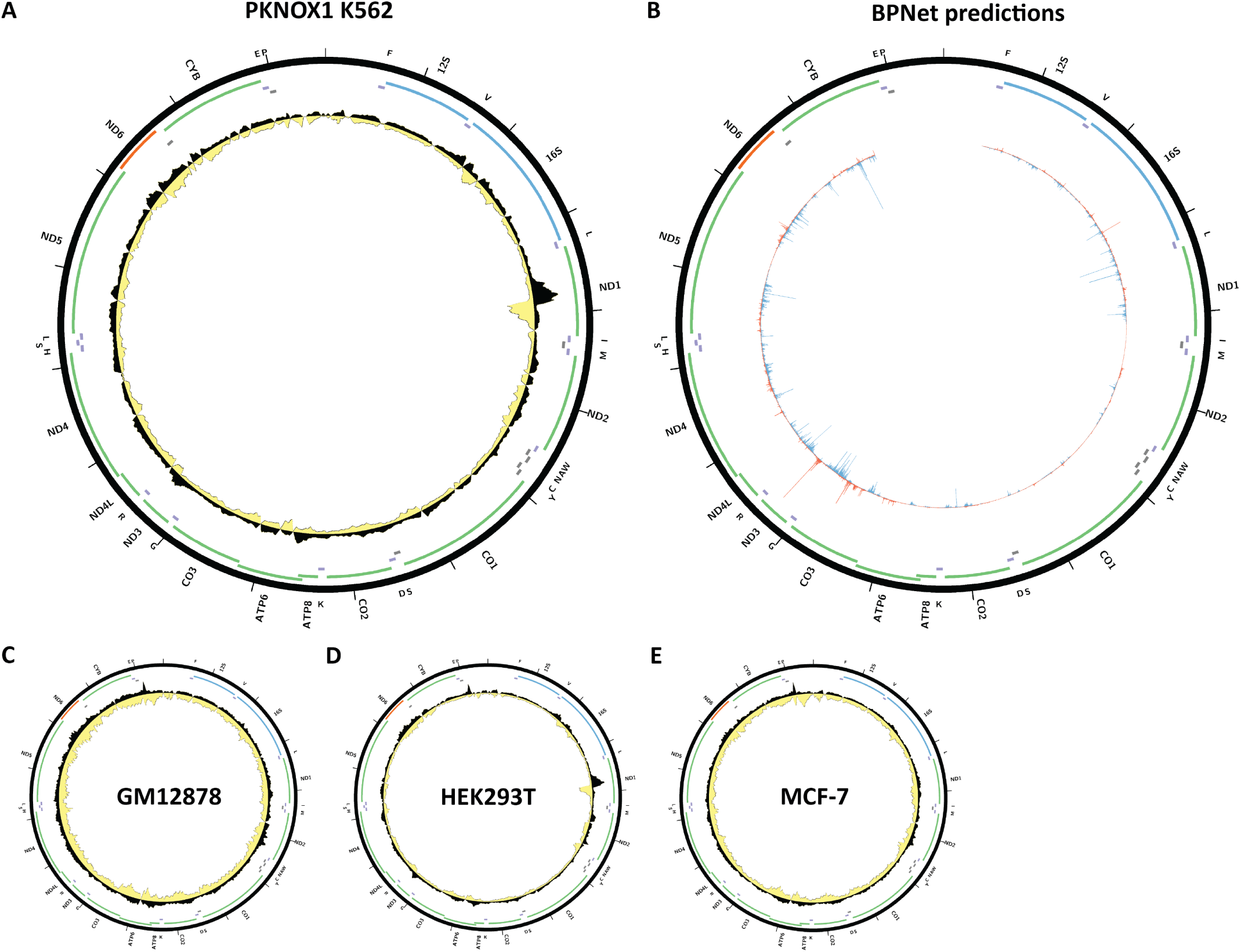
Evidence for mitochondrial genome occupancy by the PKNOX1 transcription factor. Black and yellow tracks show the forward- and reverse-strand ChIP-seq coverage over chrM. (A) K562 ChIP-seq (ENCODE ID ENCSR115SMW; antibody: GeneTex GTX114991, Lot ID 40870); (B) BPNet predictions over chrM (ENCODE ID ENCSR509JOV); (C) GM12878 ChIP-seq (ENCODE ID ENCSR711XNY; antibody: GeneTex GTX114991, Lot ID 40870); (D) HEK293T ChIP-seq (ENCODE ID ENCSR233FAG; antibody: GeneTex GTX114991, Lot ID 40870); MCF-7 ChIP-seq (ENCODE ID ENCSR986XYK; antibody: GeneTex GTX114991, Lot ID 40870).

**Supplementary Figure 51:**
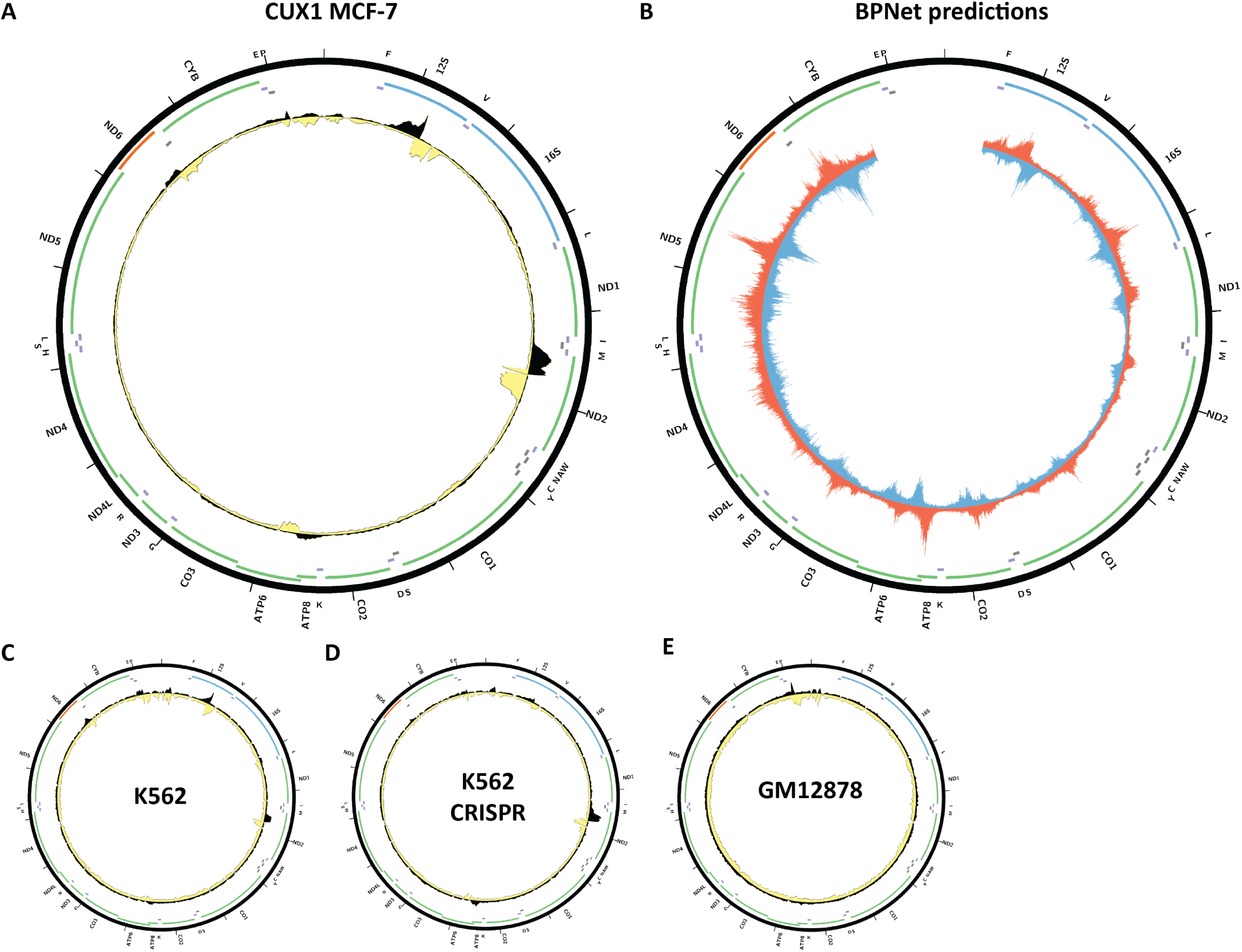
Evidence for mitochondrial genome occupancy by the CUX1transcription factor. Black and yellow tracks show the forward- and reverse-strand ChIP-seq coverage over chrM. (A) MCF-7 ChIP-seq (ENCODE ID ENCSR017CEO; antibody: Santa Cruz Biotech sc-6327, Lot ID E0709); (B) BPNet predictions over chrM (ENCODE ID ENCSR867JJN); (C) K562 ChIP-seq (ENCODE ID ENCSR000EFO; antibody: Santa Cruz Biotech sc-6327, Lot ID E0709); (D) K562 CETCH-seq (ENCODE ID ENCSR178NTX); (E) GM12878 ChIP-seq (ENCODE ID ENCSR000DYR; antibody: Santa Cruz Biotech sc-6327, Lot ID E0709).

**Supplementary Figure 52:**
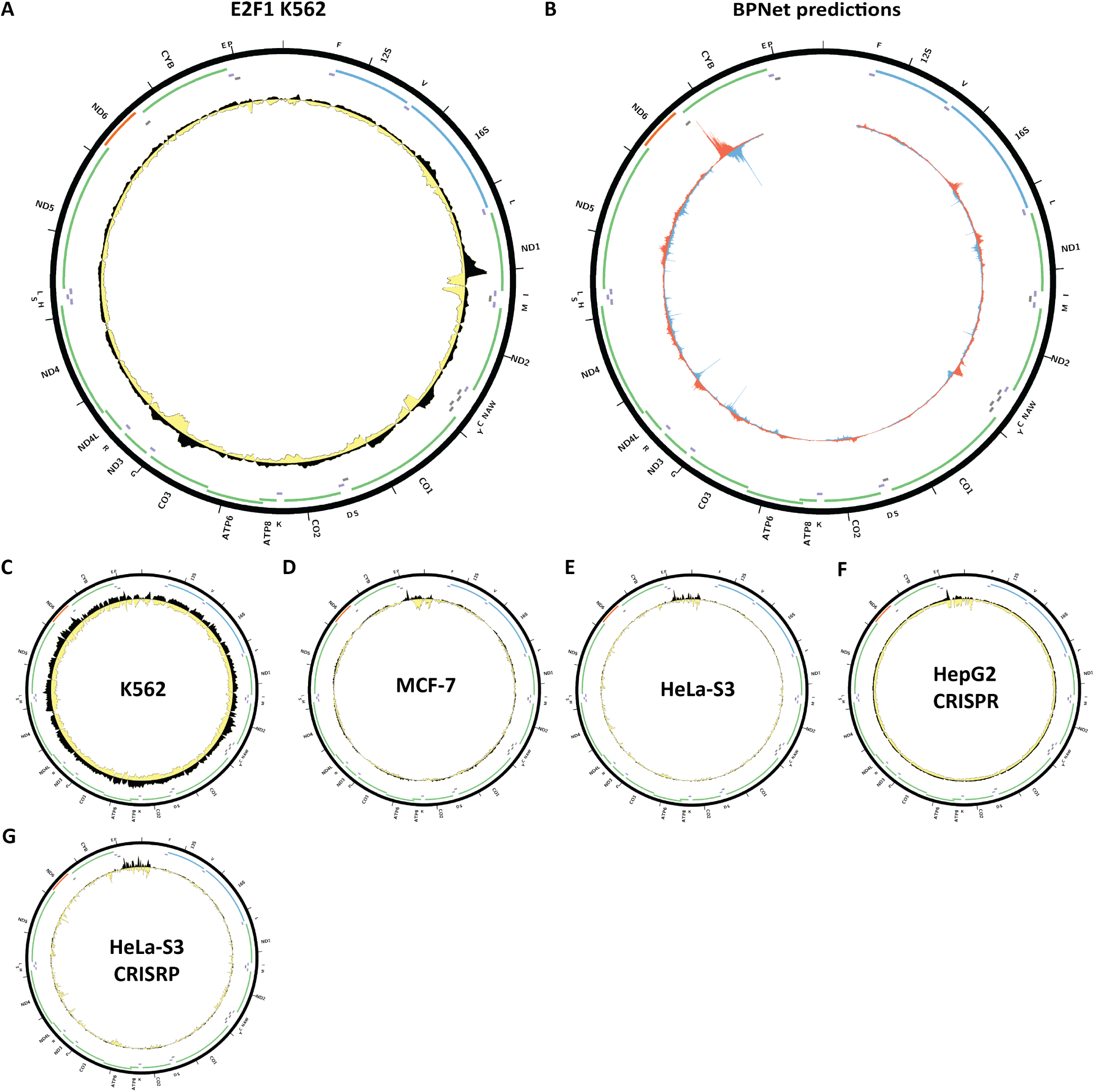
Evidence for mitochondrial genome occupancy by the E2F1 transcription factor. Black and yellow tracks show the forward- and reverse-strand ChIP-seq coverage over chrM. (A) K562 ChIP-seq (ENCODE ID ENCSR720HUL; antibody: GeneTex GTX70165, Lot ID 19267); (B) BPNet predictions over chrM; (C) K562 ChIP-seq (ENCODE ID ENCSR563LLO; antibody: Cell Signaling 3742S, Lot ID 4); (D) MCF-7 ChIP-seq (ENCODE ID ENCSR000EWX; HA-modified E2F1); (E) HeLa-S3 ChIP-seq (ENCODE ID ENCSR000EVJ; antibody: Millipore 05-379). (F) HepG2 CETCH-seq (ENCODE ID ENCSR717ZZW); (G) HeLa-S3 CETCH-seq (ENCODE ID ENCSR000EVM).

**Supplementary Figure 53:**
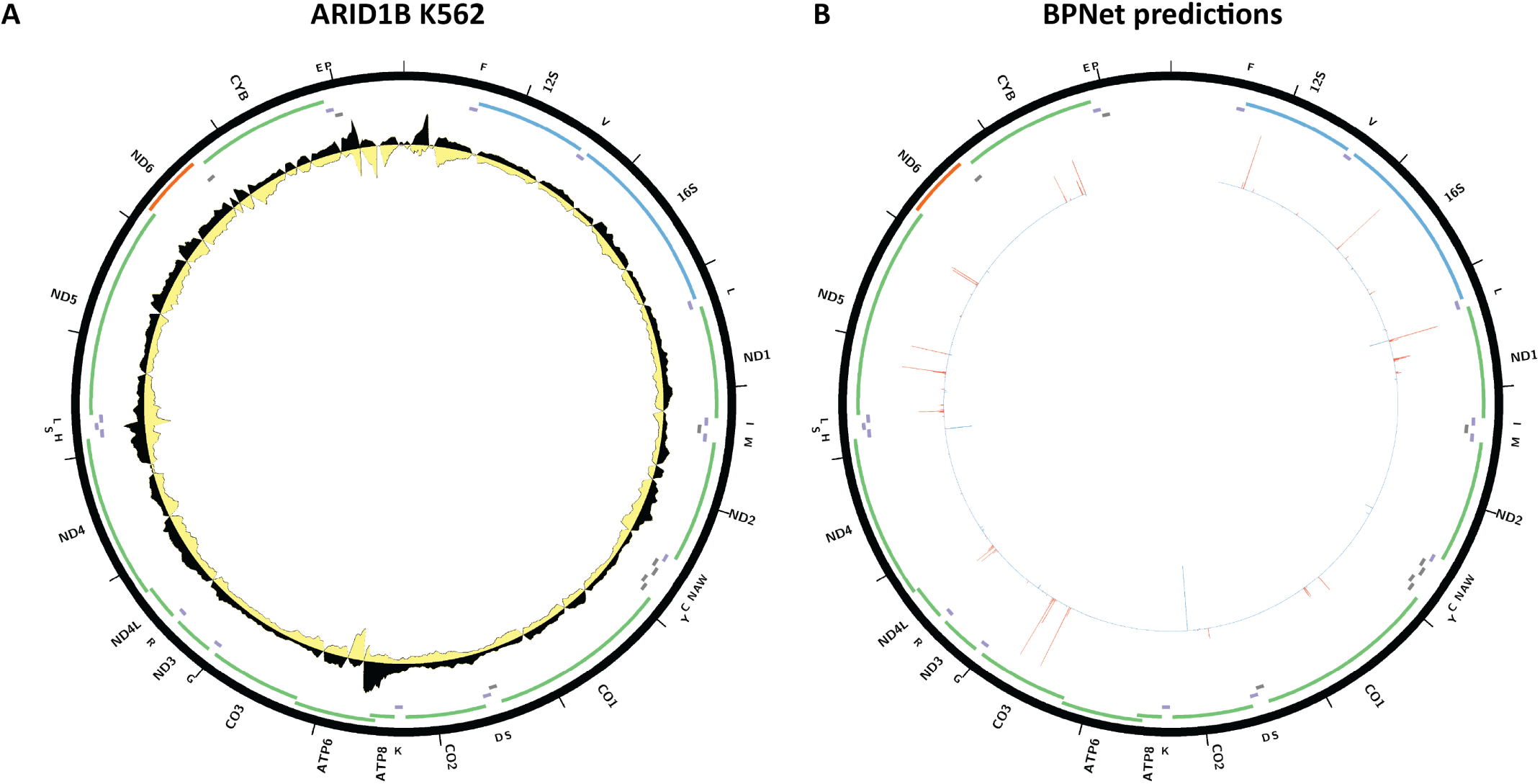
Evidence for mitochondrial genome occupancy by the ARID1Btranscription factor. Black and yellow tracks show the forward- and reverse-strand ChIP-seq coverage over chrM. (A) K562 ChIP-seq (ENCODE ID ENCSR822CCM; antibody: Bethyl Labs A301-046A); (B) BPNet predictions over chrM.

**Supplementary Figure 54:**
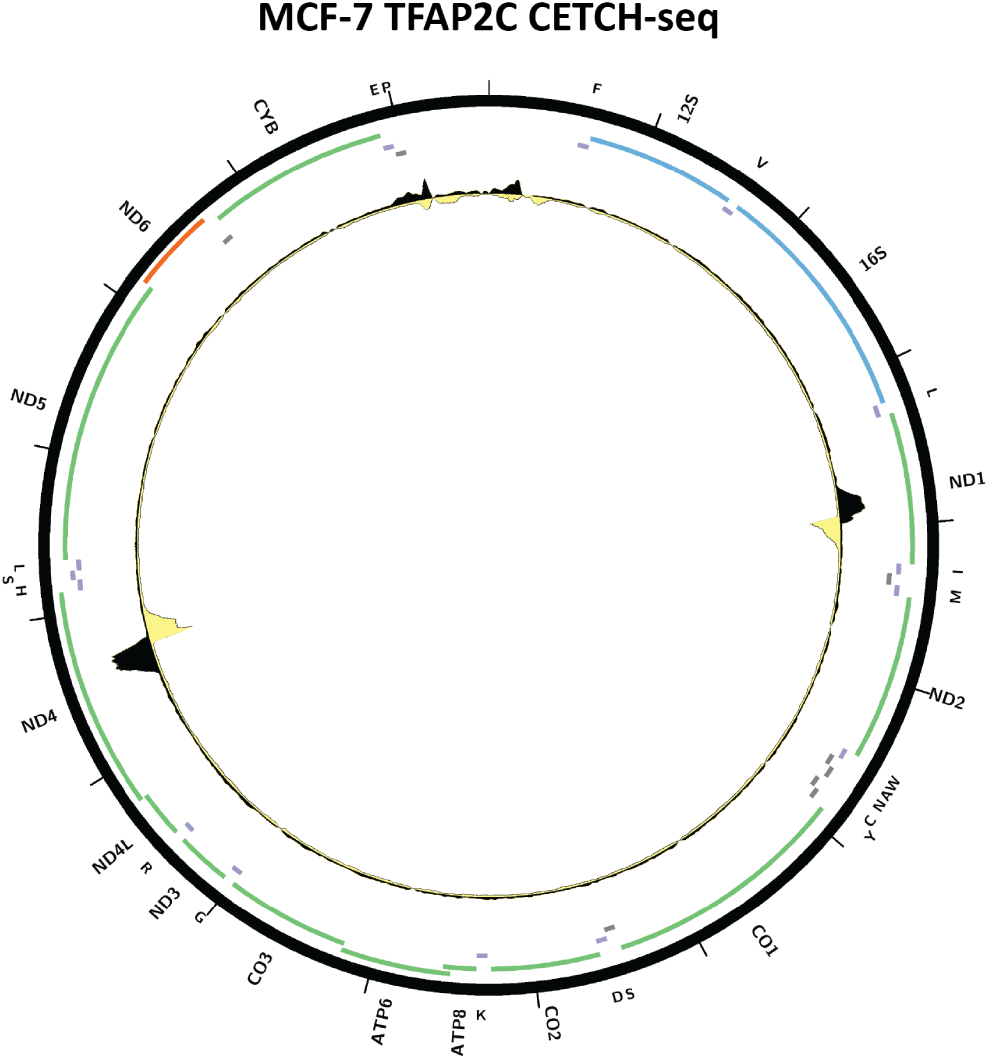
Evidence for mitochondrial genome occupancy by the TFAP2C transcription factor. Black and yellow tracks show the forward- and reverse-strand ChIP-seq coverage over chrM. CETCH-seq; ENCODE ID ENCSR742RUA.

**Supplementary Figure 55:**
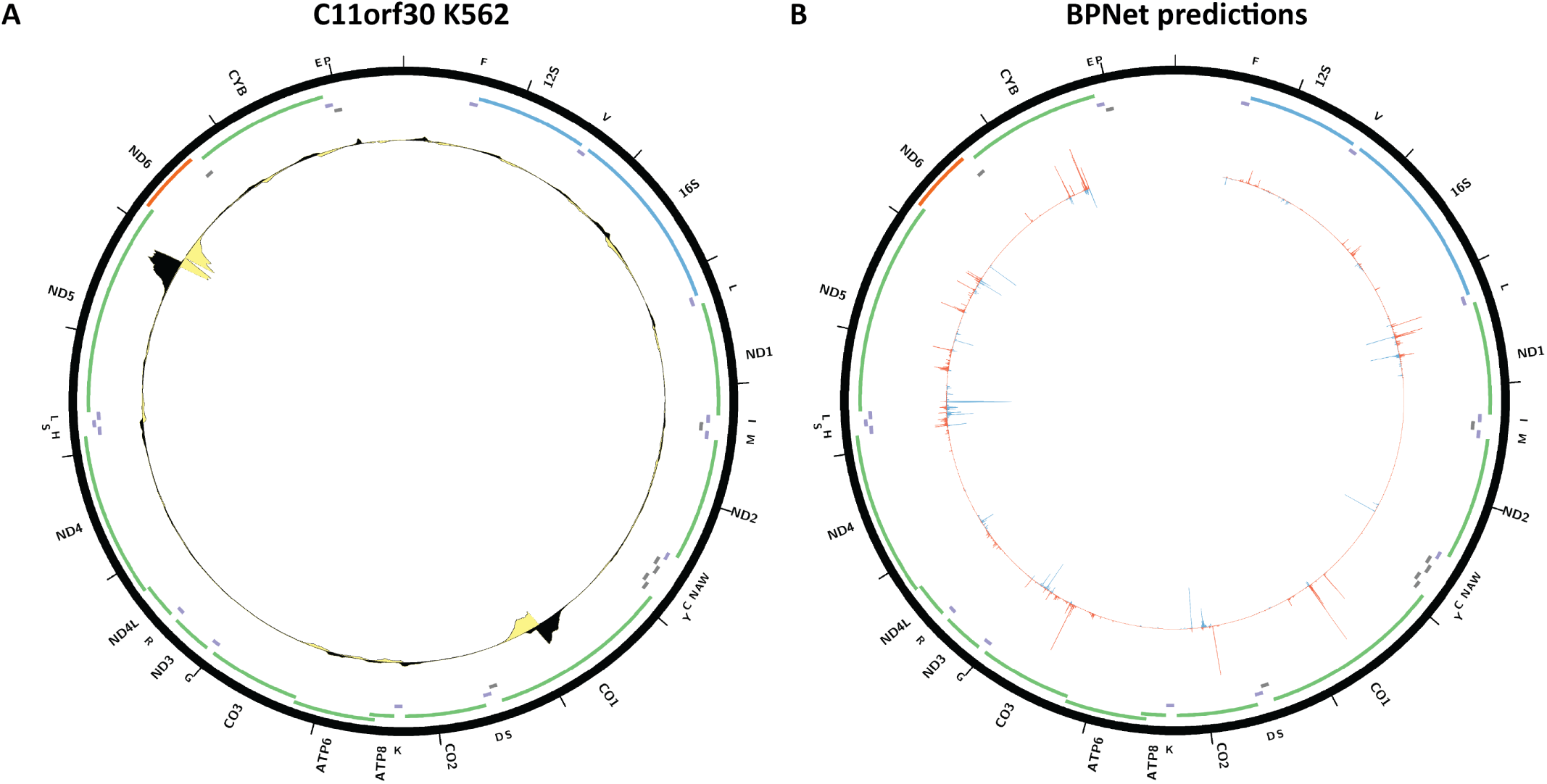
Evidence for mitochondrial genome occupancy by the C11orf30protein. Black and yellow tracks show the forward- and reverse-strand ChIP-seq coverage over chrM. (A) K562 ChIP-seq (ENCODE ID ENCSR350XWY; antibody: Bethyl Labs A300-253A, Lot ID 2); (B) BPNet predictions over chrM (ENCODE ID ENCSR350XWY).

**Supplementary Figure 56:**
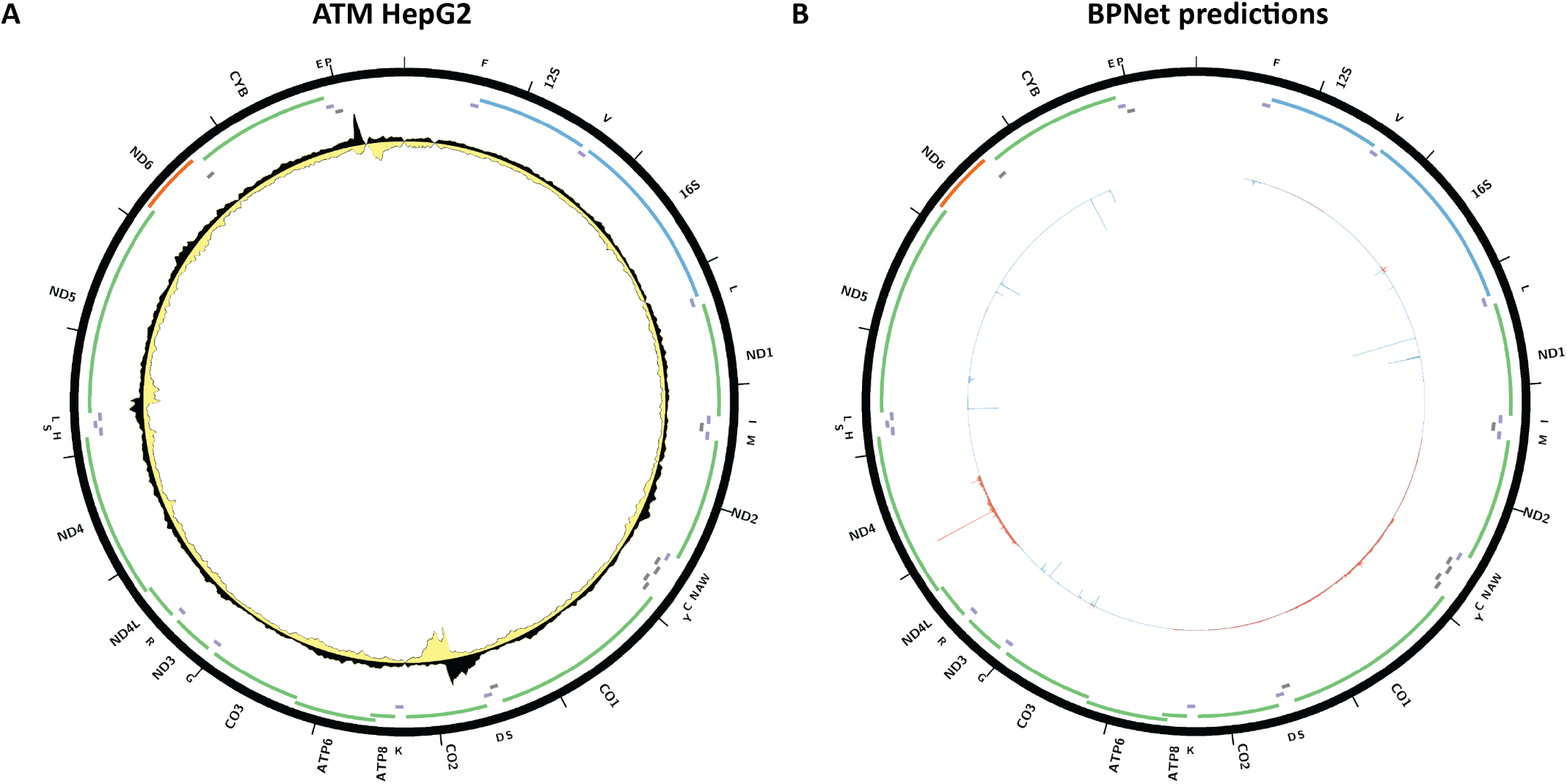
Evidence for mitochondrial genome occupancy by the ATM protein. Black and yellow tracks show the forward- and reverse-strand ChIP-seq coverage over chrM. (A) HepG2ChIP-seq (ENCODE ID ENCSR859JGF; antibody: Bethyl Labs A300-135A); (B) BPNet predictions over chrM.

## References

1. Lambert SA, Jolma A, Campitelli LF, Das PK, Yin Y, Albu M, Chen X, Taipale J, Hughes TR, Weirauch MT. 2018. The Human Transcription Factors. Cell 172(4):650–665.

2. Nass S, Nass MM, Hennix U. 1965. Deoxyribonucleic acid in isolated rat-liver mitochondria. Biochim Biophys Acta 95:426–435.

3. Anderson S, Bankier AT, Barrell BG, de Bruijn MH, Coulson AR, Drouin J, Eperon IC, Nierlich DP, Roe BA, Sanger F, Schreier PH, Smith AJ, Staden R, Young IG. 1981. Sequence and organization of the human mitochondrial genome. Nature 290(5806):457–465.

4. Bibb MJ, Van Etten RA, Wright CT, Walberg MW, Clayton DA. 1981. Sequence and gene organization of mouse mitochondrial DNA. Cell 26(2 Pt 2):167–180.

5. Shadel GS, Clayton DA. 1997. Mitochondrial DNA maintenance in vertebrates. Annu Rev Biochem 66:409–435.

6. Cantatore P, Attardi G. 1980. Mapping of nascent light and heavy strand transcripts on the physical map of HeLa cell mitochondrial DNA. Nucleic Acids Res 8(12):2605–2625.

7. Montoya J, Christianson T, Levens D, Rabinowitz M, Attardi G. 1982. Identification of initiation sites for heavy-strand and light-strand transcription in human mitochondrial DNA. Proc Natl Acad Sci 79(23):7195–7199.

8. Tiranti V, Savoia A, Forti F, D’Apolito MF, Centra M, Rocchi M, Zeviani M. 1997. Identification of the gene encoding the human mitochondrial RNA polymerase (h-mtRPOL) by cyberscreening of the Expressed Sequence Tags database. Hum Mol Genet 6(4):615–625.

9. Fisher RP, Clayton DA. 1988. Purification and characterization of human mitochondrial transcription factor 1. Mol. Cell. Biol 8(8):3496–3509.

10. Fisher RP, Clayton DA. 1985. A transcription factor required for promoter recognition by human mitochondrial RNA polymerase. Accurate initiation at the heavy- and light-strand promoters dissected and reconstituted in vitro. J Biol Chem 260(20):11330–11338.

11. Wang YE, Marinov GK, Wold BJ, Chan DC. 2013. Genome-wide analysis reveals coating of the mitochondrial genome by TFAM. PLoS ONE 8(8):e74513.

12. Falkenberg M, Gaspari M, Rantanen A, Trifunovic A, Larsson NG, Gustafsson CM. 2002. Mitochondrial transcription factors B1 and B2 activate transcription of human mtDNA. Nat Genet 31(3):289–294.

13. Gaspari M, Falkenberg M, Larsson NG, Gustafsson CM. 2004. The mitochondrial RNA polymerase contributes critically to promoter specificity in mammalian cells. EMBO J 23(23):4606–4614.

14. Shutt TE, Bestwick M, Shadel GS. 2011. The core human mitochondrial transcription initiation complex: It only takes two to tango. Transcription 2(2):55–59.

15. Mereschkowski C. 1905. Über Natur und Ursprung der Chromatophoren im Pflanzenreiche. Biol Centralbl 25:593–604.

16. Mereschkowsky K. 1910. Theorie der zwei Plasmaarten als Grundlage der Symbiogenesis, einer neuen Lehre von der Ent-stehung der Organismen. Biol Centralbl 30:353–367.

17. Sagan L. 1967. On the origin of mitosing cells. J Theor Biol 14(3):255–274.

18. Schwartz RM, Dayhoff MO. 1978. Origins of prokaryotes, eukaryotes, mitochondria, and chloroplasts. Science 199(4327):395–403.

19. Yang D, Oyaizu Y, Oyaizu H, Olsen GJ, Woese CR. 1985. Mitochondrial origins. Proc Natl Acad Sci U S A 82(13):4443–4447.

20. Gray MW, Burger G, Lang BF. 1999. Mitochondrial evolution. Science 283(5407):1476–1481.

21. Andersson SG, Zomorodipour A, Andersson JO, Sicheritz–Pontén T, Alsmark UC, Podowski RM, Näslund AK, Eriksson AS, Winkler HH, Kurland CG. 1998. The genome sequence of *Rickettsia prowazekii* and the origin of mitochondria. Nature 396(6707):133–140.

22. Esser C, Ahmadinejad N, Wiegand C, Rotte C, Sebastiani F, Gelius–Dietrich G, Henze K, Kretschmann E, Richly E, Leister D, Bryant D, Steel MA, Lockhart PJ, Penny D, Martin W. 2004. A genome phylogeny for mitochondria among *α*-proteobacteria and a predominantly eubacterial ancestry of yeast nuclear genes. Mol Biol Evol 21(9):1643–1660.

23. Fitzpatrick DA, Creevey CJ, McInerney JO. 2006. Genome phylogenies indicate a meaningful *α*-proteobacterial phylogeny and support a grouping of the mitochondria with the Rickettsiales. Mol Biol Evol 23(1):74–85.

24. Leigh-Brown S, Enriquez JA, Odom DT. 2010. Nuclear transcription factors in mammalian mitochondria. Genome Biol 11(7):215

25. Rubalcava-Gracia D, García-Villegas R, Larsson NG. 2023. No role for nuclear transcription regulators in mammalian mitochondria? Mol Cell 83(6):832–842.

26. Marinov GK, Wang YE, Chan D, Wold BJ. 2014. Evidence for site-specific occupancy of the mitochondrial genome by nuclear transcription factors. PLoS One 9(1):e84713.

27. Blumberg A, Sri Sailaja B, Kundaje A, Levin L, Dadon S, Shmorak S, Shaulian E, Meshorer E, Mishmar D. 2014. Transcription factors bind negatively selected sites within human mtDNA genes. Genome Biol Evol 6(10):2634–2646.

28. Demonacos C, Tsawdaroglou NC, Djordjevic-Markovic R, Papalopoulou M, Galanopoulos V, Papadogeorgaki S, Sekeris CE. 1993. Import of the glucocorticoid receptor into rat liver mitochondria in vivo and in vitro. J Steroid Biochem Mol Biol 46(3):401–13.

29. Demonacos C, Djordjevic-Markovic R, Tsawdaroglou N, Sekeris CE. 1995. The mitochondrion as a primary site of action of glucocorticoids: the interaction of the glucocorticoid receptor with mitochondrial DNA sequences showing partial similarity to the nuclear glucocorticoid responsive elements. J Steroid Biochem Mol Biol 55(1):43–55.

30. Koufali MM, Moutsatsou P, Sekeris CE, Breen KC. 2003. The dynamic localization of the glucocorticoid receptor in rat C6 glioma cell mitochondria. Mol Cell Endocrinol 209(1-2):51–60.

31. Psarra AM, Solakidi S, Sekeris CE. 2006. The mitochondrion as a primary site of action of steroid and thyroid hormones: presence and action of steroid and thyroid hormone receptors in mitochondria of animal cells. Mol Cell Endocrinol 246(1-2):21–33.

32. Wrutniak C, Cassar-Malek I, Marchal S, Rascle A, Heusser S, Keller JM, Fĺechon J, Dauça M, Samarut J, Ghysdael J, Cabello, G. 1995. A 43-kDa protein related to c-Erb A *α*1 is located in the mitochondrial matrix of rat liver. J Biol Chem 270(27):16347–54.

33. Casas F, Rochard P, Rodier A, Cassar-Malek I, Marchal-Victorion S, Wiesner RJ, Cabello G, Wrutniak C. 1999. A variant form of the nuclear triiodothy- ronine receptor c-ErbA*α*1 plays a direct role in regulation of mitochondrial RNA synthesis. Mol Cell Biol 19(12):7913–24.

34. Enríquez JA, Fernández-Sílva P, Montoya J. 1999a. Autonomous regulation in mammalian mitochondrial DNA transcription. Biol Chem 380(7-8):737–47.

35. Enríquez JA, Fernández-Silva P, Garrido-Pérez N, Lóopez-Pérez MJ, Pérez-Martos A, Montoya J. 1999b, Direct regulation of mitochondrial RNA synthesis by thyroid hormone. Mol Cell Biol 19(1):657–70.

36. Cammarota M, Paratcha G, Bevilaqua LR, Levi de Stein M, Lopez M, Pellegrino de Iraldi A, Izquierdo I, Medina JH. 1999. Cyclic AMP-responsive element binding protein in brain mitochondria. J Neurochem 72(6):2272–7.

37. Lee J, Kim CH, Simon DK, Aminova LR, Andreyev AY, Kushnareva YE, Murphy AN, Lonze BE, Kim KS, Ginty DD, Ferrante RJ, Ryu H, Ratan RR. 2005. Mitochondrial cyclic AMP response element-binding protein (CREB) mediates mitochondrial gene expression and neuronal survival. J Biol Chem 280(49):40398–401.

38. Ryu H, Lee J, Impey S, Ratan RR, Ferrante RJ. 2005. Antioxidants modulate mitochondrial PKA and increase CREB binding to D-loop DNA of the mitochondrial genome in neurons. Proc Natl Acad Sci U S A 102(39):13915–20.

39. De Rasmo D, Signorile A, Roca E, Papa S. 2009. cAMP response element-binding protein (CREB) is imported into mitochondria and promotes protein synthesis. FEBS J 276(16):4325–33.

40. Marchenko ND, Zaika A, Moll UM. 2000. Death signal-induced localization of p53 protein to mitochondria. A potential role in apoptotic signaling. J Biol Chem 275(21):16202–12.

41. Yoshida Y, Izumi H, Torigoe T, Ishiguchi H, Itoh H, Kang D, Kohno K. 2003. P53 physically interacts with mitochondrial transcription factor A and differentially regulates binding to damaged DNA. Cancer Res 63(13):3729–34.

42. Heyne K, Mannebach S, Wuertz E, Knaup KX, Mahyar-Roemer M, Roemer K. 2004. Identification of a putative p53 binding sequence within the human mitochondrial genome. FEBS Lett 578(1-2):198–202.

43. Achanta G, Sasaki R, Feng L, Carew JS, Lu W, Pelicano H, Keating MJ, Huang P. 2005. Novel role of p53 in maintaining mitochondrial genetic stability through interaction with DNA Pol gamma. EMBO Journal 24(19):3482–92.

44. Monje P, Boland R. 2001. Subcellular distribution of native estrogen receptor *α* and *β* isoforms in rabbit uterus and ovary. J Cell Biochem 82(3):467–79.

45. Chen JQ, Delannoy M, Cooke C, Yager JD. 2004. Mitochondrial localization of ER*α* and ER*β* in human MCF7 cells. Am J Physiol Endocrinol Metab 286(6):E1011–22.

46. Wegrzyn J, Potla R, Chwae YJ, Sepuri NB, Zhang Q, Koeck T, Derecka M, Szczepanek K, Szelag M, Gornicka A, Moh A, Moghaddas S, Chen Q, Bobbili S, Cichy J, Dulak J, Baker DP, Wolfman A, Stuehr D, Hassan MO, Fu XY, Avadhani N, Drake JI, Fawcett P, Lesnefsky EJ, Larner AC. 2009. Function of mitochondrial Stat3 in cellular respiration. Science 323(5915):793–7.

47. Casas F, Domenjoud L, Rochard P, Hatier R, Rodier A, Daury L, Bianchi A, Kremarik-Bouillaud P, Becuwe P, Keller J, Schohn H, Wrutniak-Cabello C, Cabello G, Dauça M. 2000. A 45 kDa protein related to PPAR*γ*2, induced by peroxisome proliferators, is located in the mitochondrial matrix. FEBS Lett 478(1-2):4–8.

48. Ogita K, Okuda H, Kitano M, Fujinami Y, Ozaki K, Yoneda Y. 2002. Localization of activator protein-1 complex with DNA binding activity in mitochondria of murine brain after in vivo treatment with kainate. J Neurosci 22(7):2561–70.

49. Ogita K, Fujinami Y, Kitano M, Yoneda Y. 2003. Transcription factor activator protein-1 expressed by kainate treatment can bind to the non-coding region of mitochondrial genome in murine hippocampus. J Neurosci Res 73(6):794–802.

50. She H, Yang Q, Shepherd K, Smith Y, Miller G, Testa C, Mao Z. 2011. Direct regulation of complex I by mitochondrial MEF2D is disrupted in a mouse model of Parkinson disease and in human patients. J Clin Invest 121(3):930–940.

51. Johnson DS, Mortazavi A, Myers RM, Wold B. 2007. Genome-wide mapping of in vivo protein-DNA interactions. Science 316(5830):1497–502.

52. Mikkelsen TS, Ku M, Jaffe DB, Issac B, Lieberman E, Giannoukos G, Alvarez P, Brockman W, Kim TK, Koche RP, Lee W, Mendenhall E, O’Donovan A, Presser A, Russ C, Xie X, Meissner A, Wernig M, Jaenisch R, Nusbaum C, Lander ES, Bernstein BE. 2007. Genome-wide maps of chromatin state in pluripotent and lineage-committed cells. Nature 448(7153):553–560.

53. Barski A, Cuddapah S, Cui K, Roh TY, Schones DE, Wang Z, Wei G, Chepelev I, Zhao K. 2007. High-resolution profiling of histone methylations in the human genome. Cell 129(4):823–837.

54. Robertson G, Hirst M, Bainbridge M, Bilenky M, Zhao Y, Zeng T, Euskirchen G, Bernier B, Varhol R, Delaney A, Thiessen N, Griffith OL, He A, Marra M, Snyder M, Jones S. 2007. Genome-wide profiles of STAT1 DNA association using chromatin immunoprecipitation and massively parallel sequencing. Nat Methods 4(8):651–657.

55. ENCODE Project Consortium. 2011. A user’s guide to the encyclopedia of DNA elements (ENCODE). PLoS Biol 9(4):e1001046.

56. ENCODE Project Consortium, Dunham I, Kundaje A, Aldred SF, Collins PJ, Davis CA, Doyle F, Epstein CB, Frietze S, Harrow J, Kaul R, Khatun J, Lajoie BR, Landt SG, Lee BK, Pauli F, Rosenbloom KR, Sabo P, Safi A, Sanyal A, Shoresh N, Simon JM, Song L, Trinklein ND, Altshuler RC, Birney E, Brown JB, Cheng C, Djebali S, Dong X, Dunham I, Ernst J, Furey TS, Gerstein M, Giardine B, Greven M, Hardison RC, Harris RS, Herrero J, Hoffman MM, Iyer S, Kelllis M, Khatun J, Kheradpour P, Kundaje A, Lassman T, Li Q, Lin X, Marinov GK, Merkel A, Mortazavi A, Parker SC, Reddy TE, Rozowsky J, Schlesinger F, Thurman RE, Wang J, Ward LD, Whitfield TW, Wilder SP, Wu W, Xi HS, Yip KY, Zhuang J, Bernstein BE, Birney E, Dunham I, Green ED, Gunter C, Snyder M, Pazin MJ, Lowdon RF, Dillon LA, Adams LB, Kelly CJ, Zhang J, Wexler JR, Green ED, Good PJ, Feingold EA, Bernstein BE, Birney E, Crawford GE, Dekker J, Elinitski L, Farnham PJ, Gerstein M, Giddings MC, Gingeras TR, Green ED, Guigó R, Hardison RC, Hubbard TJ, Kellis M, Kent WJ, Lieb JD, Margulies EH, Myers RM, Snyder M, Starnatoyannopoulos JA, Tennebaum SA, Weng Z, White KP, Wold B, Khatun J, Yu Y, Wrobel J, Risk BA, Gunawardena HP, Kuiper HC, Maier CW, Xie L, Chen X, Giddings MC, Bernstein BE, Epstein CB, Shoresh N, Ernst J, Kheradpour P, Mikkelsen TS, Gillespie S, Goren A, Ram O, Zhang X, Wang L, Issner R, Coyne MJ, Durham T, Ku M, Truong T, Ward LD, Altshuler RC, Eaton ML, Kellis M, Djebali S, Davis CA, Merkel A, Dobin A, Lassmann T, Mortazavi A, Tanzer A, Lagarde J, Lin W, Schlesinger F, Xue C, Marinov GK, Khatun J, Williams BA, Zaleski C, Rozowsky J, Röder M, Kokocinski F, Abdelhamid RF, Alioto T, Antoshechkin I, Baer MT, Batut P, Bell I, Bell K, Chakrabortty S, Chen X, Chrast J, Curado J, Derrien T, Drenkow J, Dumais E, Dumais J, Duttagupta R, Fastuca M, Fejes-Toth K, Ferreira P, Foissac S, Fullwood MJ, Gao H, Gonzalez D, Gordon A, Gunawardena HP, Howald C, Jha S, Johnson R, Kapranov P, King B, Kingswood C, Li G, Luo OJ, Park E, Preall JB, Presaud K, Ribeca P, Risk BA, Robyr D, Ruan X, Sammeth M, Sandu KS, Schaeffer L, See LH, Shahab A, Skancke J, Suzuki AM, Takahashi H, Tilgner H, Trout D, Walters N, Wang H, Wrobel J, Yu Y, Hayashizaki Y, Harrow J, Gerstein M, Hubbard TJ, Reymond A, Antonarakis SE, Hannon GJ, Giddings MC, Ruan Y, Wold B, Carninci P, Guigó R, Gingeras TR, Rosenbloom KR, Sloan CA, Learned K, Malladi VS, Wong MC, Barber GP, Cline MS, Dreszer TR, Heitner SG, Karolchik D, Kent WJ, Kirkup VM, Meyer LR, Long JC, Maddren M, Raney BJ, Furey TS, Song L, Grasfeder LL, Giresi PG, Lee BK, Battenhouse A, Sheffield NC, Simon JM, Showers KA, Safi A, London D, Bhinge AA, Shestak C, Schaner MR, Kim SK, Zhang ZZ, Mieczkowski PA, Mieczkowska JO, Liu Z, McDaniell RM, Ni Y, Rashid NU, Kim MJ, Adar S, Zhang Z, Wang T, Winter D, Keefe D, Birney E, Iyer VR, Lieb JD, Crawford GE, Li G, Sandhu KS, Zheng M, Wang P, Luo OJ, Shahab A, Fullwood MJ, Ruan X, Ruan Y, Myers RM, Pauli F, Williams BA, Gertz J, Marinov GK, Reddy TE, Vielmetter J, Partridge EC, Trout D, Varley KE, Gasper C, Bansal A, Pepke S, Jain P, Amrhein H, Bowling KM, Anaya M, Cross MK, King B, Muratet MA, Antoshechkin I, Newberry KM, McCue K, Nesmith AS, Fisher-Aylor KI, Pusey B, DeSalvo G, Parker SL, Balasubramanian S, Davis NS, Meadows SK, Eggleston T, Gunter C, Newberry JS, Levy SE, Absher DM, Mortazavi A, Wong WH, Wold B, Blow MJ, Visel A, Pennachio LA, Elnitski L, Margulies EH, Parker SC, Petrykowska HM, Abyzov A, Aken B, Barrell D, Barson G, Berry A, Bignell A, Boychenko V, Bussotti G, Chrast J, Davidson C, Derrien T, Despacio-Reyes G, Diekhans M, Ezkurdia I, Frankish A, Gilbert J, Gonzalez JM, Griffiths E, Harte R, Hendrix DA, Howald C, Hunt T, Jungreis I, Kay M, Khurana E, Kokocinski F, Leng J, Lin MF, Loveland J, Lu Z, Manthravadi D, Mariotti M, Mudge J, Mukherjee G, Notredame C, Pei B, Rodriguez JM, Saunders G, Sboner A, Searle S, Sisu C, Snow C, Steward C, Tanzer A, Tapanan E, Tress ML, van Baren MJ, Walters N, Washieti S, Wilming L, Zadissa A, Zhengdong Z, Brent M, Haussler D, Kellis M, Valencia A, Gerstein M, Raymond A, Guigó R, Harrow J, Hubbard TJ, Landt SG, Frietze S, Abyzov A, Addleman N, Alexander RP, Auerbach RK, Balasubramanian S, Bettinger K, Bhardwaj N, Boyle AP, Cao AR, Cayting P, Charos A, Cheng Y, Cheng C, Eastman C, Euskirchen G, Fleming JD, Grubert F, Habegger L, Hariharan M, Harmanci A, Iyenger S, Jin VX, Karczewski KJ, Kasowski M, Lacroute P, Lam H, Larnarre-Vincent N, Leng J, Lian J, LindahlAllen M, Min R, Miotto B, Monahan H, Moqtaderi Z, Mu XJ, O’Geen H, Ouyang Z, Patacsil D, Pei B, Raha D, Ramirez L, Reed B, Rozowsky J, Sboner A, Shi M, Sisu C, Slifer T, Witt H, Wu L, Xu X, Yan KK, Yang X, Yip KY, Zhang Z, Struhl K, Weissman SM, Gerstein M, Farnham PJ, Snyder M, Tenebaum SA, Penalva LO, Doyle F, Karmakar S, Landt SG, Bhanvadia RR, Choudhury A, Domanus M, Ma L, Moran J, Patacsil D, Slifer T, Victorsen A, Yang X, Snyder M, White KP, Auer T, Centarin L, Eichenlaub M, Gruhl F, Heerman S, Hoeckendorf B, Inoue D, Kellner T, Kirchmaier S, Mueller C, Reinhardt R, Schertel L, Schneider S, Sinn R, Wittbrodt B, Wittbrodt J, Weng Z, Whitfield TW, Wang J, Collins PJ, Aldred SF, Trinklein ND, Partridge EC, Myers RM, Dekker J, Jain G, Lajoie BR, Sanyal A, Balasundaram G, Bates DL, Byron R, Canfield TK, Diegel MJ, Dunn D, Ebersol AK, Ebersol AK, Frum T, Garg K, Gist E, Hansen RS, Boatman L, Haugen E, Humbert R, Jain G, Johnson AK, Johnson EM, Kutyavin TM, Lajoie BR, Lee K, Lotakis D, Maurano MT, Neph SJ, Neri FV, Nguyen ED, Qu H, Reynolds AP, Roach V, Rynes E, Sabo P, Sanchez ME, Sandstrom RS, Sanyal A, Shafer AO, Stergachis AB, Thomas S, Thurman RE, Vernot B, Vierstra J, Vong S, Wang H, Weaver MA, Yan Y, Zhang M, Akey JA, Bender M, Dorschner MO, Groudine M, MacCoss MJ, Navas P, Stamatoyannopoulos G, Kaul R, Dekker J, Stamatoyannopoulos JA, Dunham I, Beal K, Brazma A, Flicek P, Herrero J, Johnson N, Keefe D, Lukk M, Luscombe NM, Sobral D, Vaquerizas JM, Wilder SP, Batzoglou S, Sidow A, Hussami N, Kyriazopoulou-Panagiotopoulou S, Libbrecht MW, Schaub MA, Kundaje A, Hardison RC, Miller W, Giardine B, Harris RS, Wu W, Bickel PJ, Banfai B, Boley NP, Brown JB, Huang H, Li Q, Li JJ, Noble WS, Bilmes JA, Buske OJ, Hoffman MM, Sahu AO, Kharchenko PV, Park PJ, Baker D, Taylor J, Weng Z, Iyer S, Dong X, Greven M, Lin X, Wang J, Xi HS, Zhuang J, Gerstein M, Alexander RP, Balasubramanian S, Cheng C, Harmanci A, Lochovsky L, Min R, Mu XJ, Rozowsky J, Yan KK, Yip KY, Birney E. 2012. An integrated encyclopedia of DNA elements in the human genome. Nature 489(7414):57–74.

57. Mouse ENCODE Consortium, Stamatoyannopoulos JA, Snyder M, Hardison R, Ren B, Gingeras T, Gilbert DM, Groudine M, Bender M, Kaul R, Canfield T, Giste E, Johnson A, Zhang M, Balasundaram G, Byron R, Roach V, Sabo PJ, Sandstrom R, Stehling AS, Thurman RE, Weissman SM, Cayting P, Hariharan M, Lian J, Cheng Y, Landt SG, Ma Z, Wold BJ, Dekker J, Crawford GE, Keller CA, Wu W, Morrissey C, Kumar SA, Mishra T, Jain D, Byrska-Bishop M, Blankenberg D, Lajoie BR, Jain G, Sanyal A, Chen KB, Denas O, Taylor J, Blobel GA, Weiss MJ, Pimkin M, Deng W, Marinov GK, Williams BA, Fisher-Aylor KI, Desalvo G, Kiralusha A, Trout D, Amrhein H, Mortazavi A, Edsall L, McCleary D, Kuan S, Shen Y, Yue F, Ye Z, Davis CA, Zaleski C, Jha S, Xue C, Dobin A, Lin W, Fastuca M, Wang H, Guigo R, Djebali S, Lagarde J, Ryba T, Sasaki T, Malladi VS, Cline MS, Kirkup VM, Learned K, Rosenbloom KR, Kent WJ, Feingold EA, Good PJ, Pazin M, Lowdon RF, Adams LB. 2012. An encyclopedia of mouse DNA elements (Mouse ENCODE). Genome Biol 13(8):418.

58. Gerstein MB, Lu ZJ, Van Nostrand EL, Cheng C, Arshinoff BI, Liu T, Yip KY, Robilotto R, Rechtsteiner A, Ikegami K, Alves P, Chateigner A, Perry M, Morris M, Auerbach RK, Feng X, Leng J, Vielle A, Niu W, Rhrissorrakrai K, Agarwal A, Alexander RP, Barber G, Brdlik CM, Brennan J, Brouillet JJ, Carr A, Cheung MS, Clawson H, Contrino S, Dannenberg LO, Dernburg AF, Desai A, Dick L, Dosé AC, Du J, Egelhofer T, Ercan S, Euskirchen G, Ewing B, Feingold EA, Gassmann R, Good PJ, Green P, Gullier F, Gutwein M, Guyer MS, Habegger L, Han T, Henikoff JG, Henz SR, Hinrichs A, Holster H, Hyman T, Iniguez AL, Janette J, Jensen M, Kato M, Kent WJ, Kephart E, Khivansara V, Khurana E, Kim JK, Kolasinska-Zwierz P, Lai EC, Latorre I, Leahey A, Lewis S, Lloyd P, Lochovsky L, Lowdon RF, Lubling Y, Lyne R, MacCoss M, Mackowiak SD, Mangone M, McKay S, Mecenas D, Merrihew G, Miller DM 3rd, Muroyama A, Murray JI, Ooi SL, Pham H, Phippen T, Preston EA, Rajewsky N, Rätsch G, Rosenbaum H, Rozowsky J, Rutherford K, Ruzanov P, Sarov M, Sasidharan R, Sboner A, Scheid P, Segal E, Shin H, Shou C, Slack FJ, Slightam C, Smith R, Spencer WC, Stinson EO, Taing S, Takasaki T, Vafeados D, Voronina K, Wang G, Washington NL, Whittle CM, Wu B, Yan KK, Zeller G, Zha Z, Zhong M, Zhou X; modEN-CODE Consortium, Ahringer J, Strome S, Gunsalus KC, Micklem G, Liu XS, Reinke V, Kim SK, Hillier LW, Henikoff S, Piano F, Snyder M, Stein L, Lieb JD, Waterston RH. 2010. Integrative analysis of the *Caenorhabditis elegans* genome by the modENCODE project. Science 330(6012):1775–1787.

59. Négre N, Brown CD, Ma L, Bristow CA, Miller SW, Wagner U, Kheradpour P, Eaton ML, Loriaux P, Sealfon R, Li Z, Ishii H, Spokony RF, Chen J, Hwang L, Cheng C, Auburn RP, Davis MB, Domanus M, Shah PK, Morrison CA, Zieba J, Suchy S, Senderowicz L, Victorsen A, Bild NA, Grundstad AJ, Hanley D, MacAlpine DM, Mannervik M, Venken K, Bellen H, White R, Gerstein M, Russell S, Grossman RL, Ren B, Posakony JW, Kellis M, White KP. 2011. A *cis*-regulatory map of the *Drosophila* genome. Nature 471(7339):527–531.

60. modENCODE Consortium, Roy S, Ernst J, Kharchenko PV, Kheradpour P, Negre N, Eaton ML, Landolin JM, Bristow CA, Ma L, Lin MF, Washietl S, Arshinoff BI, Ay F, Meyer PE, Robine N, Washington NL, Di Stefano L, Berezikov E, Brown CD, Candeias R, Carlson JW, Carr A, Jungreis I, Marbach D, Sealfon R, Tolstorukov MY, Will S, Alekseyenko AA, Artieri C, Booth BW, Brooks AN, Dai Q, Davis CA, Duff MO, Feng X, Gorchakov AA, Gu T, Henikoff JG, Kapranov P, Li R, MacAlpine HK, Malone J, Minoda A, Nordman J, Okamura K, Perry M, Powell SK, Riddle NC, Sakai A, Samsonova A, Sandler JE, Schwartz YB, Sher N, Spokony R, Sturgill D, van Baren M, Wan KH, Yang L, Yu C, Feingold E, Good P, Guyer M, Lowdon R, Ahmad K, Andrews J, Berger B, Brenner SE, Brent MR, Cherbas L, Elgin SC, Gingeras TR, Grossman R, Hoskins RA, Kaufman TC, Kent W, Kuroda MI, Orr-Weaver T, Perrimon N, Pirrotta V, Posakony JW, Ren B, Russell S, Cherbas P, Graveley BR, Lewis S, Micklem G, Oliver B, Park PJ, Celniker SE, Henikoff S, Karpen GH, Lai EC, MacAlpine DM, Stein LD, White KP, Kellis M. 2010. Identification of functional elements and regulatory circuits by Drosophila modENCODE. Science 330(6012):1787–1797.

61. Yue F, Cheng Y, Breschi A, Vierstra J, Wu W, Ryba T, Sandstrom R, Ma Z, Davis C, Pope BD, Shen Y, Pervouchine DD, Djebali S, Thurman RE, Kaul R, Rynes E, Kirilusha A, Marinov GK, Williams BA, Trout D, Amrhein H, Fisher-Aylor K, Antoshechkin I, DeSalvo G, See LH, Fastuca M, Drenkow J, Zaleski C, Dobin A, Prieto P, Lagarde J, Bussotti G, Tanzer A, Denas O, Li K, Bender MA, Zhang M, Byron R, Groudine MT, McCleary D, Pham L, Ye Z, Kuan S, Edsall L, Wu YC, Rasmussen MD, Bansal MS, Kellis M, Keller CA, Morrissey CS, Mishra T, Jain D, Dogan N, Harris RS, Cayting P, Kawli T, Boyle AP, Euskirchen G, Kundaje A, Lin S, Lin Y, Jansen C, Malladi VS, Cline MS, Erickson DT, Kirkup VM, Learned K, Sloan CA, Rosenbloom KR, Lacerda de Sousa B, Beal K, Pignatelli M, Flicek P, Lian J, Kahveci T, Lee D, Kent WJ, Ramalho Santos M, Herrero J, Notredame C, Johnson A, Vong S, Lee K, Bates D, Neri F, Diegel M, Canfield T, Sabo PJ, Wilken MS, Reh TA, Giste E, Shafer A, Kutyavin T, Haugen E, Dunn D, Reynolds AP, Neph S, Humbert R, Hansen RS, De Bruijn M, Selleri L, Rudensky A, Josefowicz S, Samstein R, Eichler EE, Orkin SH, Levasseur D, Papayannopoulou T, Chang KH, Skoultchi A, Gosh S, Disteche C, Treuting P, Wang Y, Weiss MJ, Blobel GA, Cao X, Zhong S, Wang T, Good PJ, Lowdon RF, Adams LB, Zhou XQ, Pazin MJ, Feingold EA, Wold B, Taylor J, Mortazavi A, Weissman SM, Stamatoyannopoulos JA, Snyder MP, Guigo R, Gingeras TR, Gilbert DM, Hardison RC, Beer MA, Ren B; Mouse ENCODE Consortium. 2014. A comparative encyclopedia of DNA elements in the mouse genome. Nature 515(7527):355–364.

62. ENCODE Project Consortium, Moore JE, Purcaro MJ, Pratt HE, Epstein CB, Shoresh N, Adrian J, Kawli T, Davis CA, Dobin A, Kaul R, Halow J, Van Nostrand EL, Freese P, Gorkin DU, Shen Y, He Y, Mackiewicz M, Pauli-Behn F, Williams BA, Mortazavi A, Keller CA, Zhang XO, Elhajjajy SI, Huey J, Dickel DE, Snetkova V, Wei X, Wang X, RiveraMulia JC, Rozowsky J, Zhang J, Chhetri SB, Zhang J, Victorsen A, White KP, Visel A, Yeo GW, Burge CB, LÃfÂ©cuyer E, Gilbert DM, Dekker J, Rinn J, Mendenhall EM, Ecker JR, Kellis M, Klein RJ, Noble WS, Kundaje A, Guigó R, Farnham PJ, Cherry JM, Myers RM, Ren B, Graveley BR, Gerstein MB, Pennacchio LA, Snyder MP, Bernstein BE, Wold B, Hardison RC, Gingeras TR, Stamatoyannopoulos JA, Weng Z. 2020. Expanded encyclopaedias of DNA elements in the human and mouse genomes. Nature 583(7818):699–710.

63. Ching T, Himmelstein DS, Beaulieu-Jones BK, Kalinin AA, Do BT, Way GP, Ferrero E, Agapow PM, Zietz M, Hoffman MM, Xie W, Rosen GL, Lengerich BJ, Israeli J, Lanchantin J, Woloszynek S, Carpenter AE, Shrikumar A, Xu J, Cofer EM, Lavender CA, Turaga SC, Alexandari AM, Lu Z, Harris DJ, DeCaprio D, Qi Y, Kundaje A, Peng Y, Wiley LK, Segler MHS, Boca SM, Swamidass SJ, Huang A, Gitter A, Greene CS. 2018. Opportunities and obstacles for deep learning in biology and medicine. J R Soc Interface 15(141). pii: 20170387.

64. Savic D, Partridge EC, Newberry KM, Smith SB, Meadows SK, Roberts BS, Mackiewicz M, Mendenhall EM, Myers RM. 2015. CETCh-seq: CRISPR epitope tagging ChIP-seq of DNA-binding proteins. Genome Res 25(10):1581–1589.

65. O’Gorman S, Fox DT, Wahl GM. 1991. Recombinasemediated gene activation and site-specific integration in mammalian cells. Science 251(4999):1351–1355

66. Dudek J, Rehling P, van der Laan M. 2013. Mitochondrial protein import: common principles and physiological networks. Biochim Biophys Acta 1833(2):274– 285.

67. Busch JD, Fielden LF, Pfanner N, Wiedemann N. 2023. Mitochondrial protein transport: Versatility of translocases and mechanisms. Mol Cell 83(6):890– 991.

68. Bogenhagen D, Clayton DA. 1974. The number of mitochondrial deoxyribonucleic acid genomes in mouse L and human HeLa cells. Quantitative isolation of mitochondrial deoxyribonucleic acid. J Biol Chem 249(24):7991–5.

69. Hazkani-Covo E, Zeller RM, Martin W. 2010. Molecular poltergeists: mitochondrial DNA copies (numts) in sequenced nuclear genomes. PLoS Genet 6(2):e1000834.

70. Avsec Z^̌^, Weilert M, Shrikumar A, Krueger S, Alexandari A, Dalal K, Fropf R, McAnany C, Gagneur J, Kundaje A, Zeitlinger J. 2021. Base-resolution models of transcription-factor binding reveal soft motif syntax. Nat Genet 53(3):354–366.

71. Kharchenko PV, Tolstorukov MY, Park PJ. 2008. Design and analysis of ChIP-seq experiments for DNAbinding proteins. Nat. Biotechnol 26: 1351–1359.

72. Landt SG, Marinov GK, Kundaje A, Kheradpour P, Pauli F, Batzoglou S, Bernstein BE, Bickel P, Brown JB, Cayting P, Chen Y, DeSalvo G, Epstein C, Fisher-Aylor KI, Euskirchen G, Gerstein M, Gertz J, Hartemink AJ, Hoffman MM, Iyer VR, Jung YL, Karmakar S, Kellis M, Kharchenko PV, Li Q, Liu T, Liu XS, Ma L, Milosavljevic A, Myers RM, Park PJ, Pazin MJ, Perry MD, Raha D, Reddy TE, Rozowsky J, Shoresh N, Sidow A, Slattery M, Stamatoyannopoulos JA, Tolstorukov MY, White KP, Xi S, Farnham PJ, Lieb JD, Wold BJ, Snyder M. 2012. ChIP-seq guidelines and practices of the ENCODE and modENCODE consortia. Genome Res 22(9):1813–1831.

73. Marinov GK, Kundaje A, Park PJ, Wold BJ. 2014. Large-scale quality analysis of published ChIP-seq data. G3 (Bethesda) 4(2):209–223.

74. Marinov GK. 2017. Identification of Candidate Functional Elements in the Genome from ChIP-seq Data. Methods Mol Biol 1543:19–43.

75. Hess J, Angel P, Schorpp-Kistner M. 2004. AP-1 subunits: quarrel and harmony among siblings. J Cell Sci 117(Pt 25):5965–5973.

76. Luo Y, Hitz BC, Gabdank I, Hilton JA, Kagda MS, Lam B, Myers Z, Sud P, Jou J, Lin K, Baymuradov UK, Graham K, Litton C, Miyasato SR, Strattan JS, Jolanki O, Lee JW, Tanaka FY, Adenekan P, O’Neill E, Cherry JM. 2020. New developments on the Encyclopedia of DNA Elements (ENCODE) data portal. Nucleic Acids Res 48(D1):D882–D889.

77. Langmead B, Trapnell C, Pop M, Salzberg SL. 2009. Ultrafast and memory-efficient alignment of short DNA sequences to the human genome. Genome Biol 10(3):R25.

78. Krzywinski M, Schein J, Birol I, Connors J, Gascoyne R, Horsman D, Jones SJ, Marra MA. 2009. Circos: an information aesthetic for comparative genomics. Genome Res 19(9):1639–45.

79. Li Q, Brown JB, Huang H, Bickel PJ. 2011. Measuring reproducibility of high-throughput experiments. Ann Appl Stat 5:1752–1779

